# Peroxisome dysfunction alters metabolism of photoreceptor outer segments in human retinal pigment epithelium

**DOI:** 10.64898/2026.02.01.701576

**Authors:** Constantin Mouzaaber, Carly B. Feldman, Suzette M. Huguenin, John Y.S. Han, Erin Trombly, Qitao Zhang, Aja Rieger, Hamed Hojjat, Brandon C. Huynh, Ehsan Misaghi, Alina Radziwon, Temesgen D. Fufa, Robert B. Hufnagel, Jason M.L. Miller, Matthew D. Benson

## Abstract

Peroxisomes are ubiquitous organelles that compartmentalize metabolic reactions including lipid catabolism and cellular detoxification. Pathogenic variants in *PEX1* and *PEX6* disrupt essential peroxisome functions and cause profound neurodegenerative diseases called peroxisome biogenesis disorders (PBDs). Despite retinal degeneration and blindness occurring frequently in PBDs, precisely how impaired peroxisome activity disrupts retinal function remains to be fully explored. To address this, we differentiated *PEX1^-/-^*, *PEX6^-/-^*, and wildtype human induced pluripotent stem cells into retinal pigment epithelium (iRPE) to study the consequences of peroxisome dysfunction in this disease-relevant cell type. Despite exhibiting impaired peroxisome matrix protein import, *PEX1^-/-^* and *PEX6^-/-^*iRPE had comparable morphology, tight junctions, and expression of proteins characteristic of RPE compared to wildtype iRPE. Targeted lipid profiling revealed reduced docosahexaenoic acid, a polyunsaturated fatty acid (PUFA) essential for retinal function, and elevated lipid species exclusively metabolized by peroxisomes in *PEX1^-/-^* and *PEX6^-/-^* iRPE. Following a photoreceptor outer segment (POS) challenge, *PEX1^-/-^* and *PEX6^-/-^* iRPE demonstrated disrupted PUFA retroconversion and lipid droplet accumulation. Additionally, *PEX1^-/-^* and *PEX6^-/-^*iRPE had impaired rhodopsin degradation, lysosomal dysfunction, and reduced transepithelial electrical resistance. These findings suggest that dysregulated POS metabolism in the RPE is a potential mechanism driving retinal degeneration in patients with PBDs.

**Graphical abstract:** 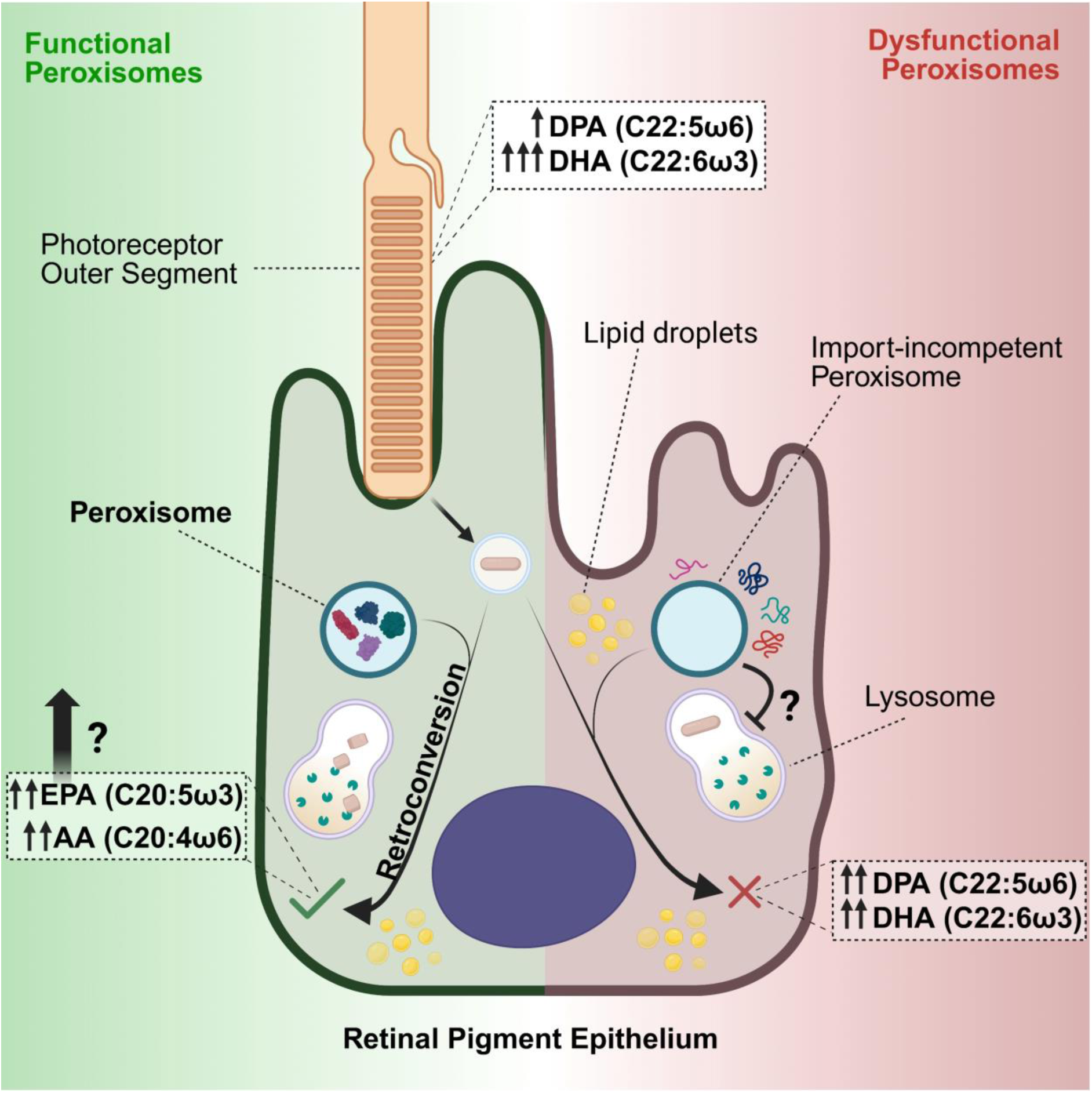

Schematic summarizing the consequences of PEX1 and PEX6 knockout on iRPE biology, including the presence of import-incompetent peroxisomes, impaired ω3 and ω6 fatty acid retroconversion, lipid droplet accumulation, and defective photoreceptor outer segment phagocytosis.

## Introduction

As a highly bioenergetic tissue, the retina is especially susceptible to local and systemic metabolic disturbances. Indeed, retinal degeneration occurs in a myriad of metabolic diseases, including peroxisomal biogenesis disorders (PBDs).^1^ The PBDs are a group of recessively inherited diseases that result in impaired function of the brain, retina, liver, and kidneys, among other organs, due to the disrupted synthesis and activity of peroxisomes. Peroxisomes are ubiquitous subcellular organelles that have essential roles in lipid metabolism and redox homeostasis via unique enzymes including ACOX1, MFP2 and ACAA1 that are guided into the peroxisome lumen using peroxisome targeting sequences 1 (PTS1) and 2 (PTS2). Two of the major reactions occurring in the peroxisome lumen are the β- and α-oxidation of lipid species such as very long-chain (≥ C22) and branched-chain fatty acids, to produce shortened or straightened FAs (≤C20) that may then undergo mitochondrial β-oxidation.

In the retina, peroxisomes metabolize very long (≥ C22)-chain polyunsaturated fatty acids (PUFAs), which are highly enriched in photoreceptor outer segments.^2^ These outer segments are continuously phagocytosed and metabolized by the retinal pigment epithelium (RPE), suggesting a critical role for intact peroxisome function in the RPE. Mice with global deficiencies in peroxisomal activity have defective phagocytosis and perturbed fatty acid β-oxidation in the RPE, which is associated with the development of retinal degeneration.^3–6^ However, the effect of peroxisome dysfunction in human RPE is understudied. To this end, we have developed cell-autonomous models to explore the function of peroxisomes in human RPE.

Pathogenic loss-of-function variants in 14 different *PEX* genes, encoding for peroxins essential for peroxisome assembly and activity, result in PBDs in humans, with *PEX1* and *PEX6* together accounting for 75% of cases.^1,7^ PEX1 and PEX6 are members of the AAA ATPase (ATPases associated with diverse cellular activities) family and form a heterohexameric complex on the peroxisome membrane. This complex facilitates the recycling of the cargo-carrying shuttle, PEX5, from the peroxisome membrane to the cytosol to enable subsequent rounds of peroxisomal matrix protein import.^1^ Despite retinal degeneration and blindness occurring in the majority of patients with *PEX1*- and *PEX6*-related PBDs,^8^ precisely how peroxisomal dysfunction impairs retinal function, and specifically induces RPE atrophy in humans, is largely unknown. To address this, we have generated and characterized the first human induced pluripotent stem cell-derived retinal pigment epithelium (iRPE) models, allowing us to study peroxisome dysfunction in a disease-relevant cell type. Our results demonstrate that *PEX1*-deficient and *PEX6*-deficient iPSCs can be differentiated into mature RPE but have cell-autonomous defects in lipid metabolism and impaired phagocytosis of photoreceptor outer segments, highlighting a novel disease mechanism in the RPE in PBDs. In addition, our iRPE models serve as novel *in vitro* platforms to broaden our understanding of the integral roles of peroxisomes in the RPE.

## Results

### *PEX1^-/-^* and *PEX6^-/-^* iRPE Exhibit Properties of Mature RPE

To study the consequences of impaired peroxisome function in human RPE, *PEX1^-/-^*, *PEX6^-/-^*, and isogenic wildtype iPSCs were concurrently differentiated into RPE (Figure 1A). All three iRPE lines achieved differentiation milestones (RPE progenitor, committed RPE, immature RPE, and mature RPE) at similar time points based on daily manual inspection with light microscopy throughout differentiation. At the mature RPE stage, all three iRPE lines expressed proteins consistent with differentiated RPE, including TYRP1, PAX6, PMEL17, BEST1, and MITF (Figure 1B) quantified by flow cytometry. In addition, there were no differences in the proportion of cells expressing RPE signature proteins across iRPE lines. All three iRPE lines developed pigment (Supplementary Figures 2A and B) and were observed to have small regions in the RPE monolayer with focal detachment from the culture surface (Supplementary Figure 2C). This latter phenomenon is due to the intrinsic fluid-pumping ability of the RPE, and is consistent with effective RPE polarization.^9,10^ In addition, all three iRPE lines formed zona occludins-1-positive tight junctions evident by immunofluorescence microscopy (Figure 1C). There were no morphological differences between the three iRPE lines in terms of cell area or number of neighboring cells using REShAPE,^11^ (Figures 1D and E) and the iRPE cell area was consistent with human RPE cell area in vivo.^12^ Finally, all three iRPE lines achieved comparable transepithelial electrical resistance (TEER) (Figure 1F).

**Figure 1:**
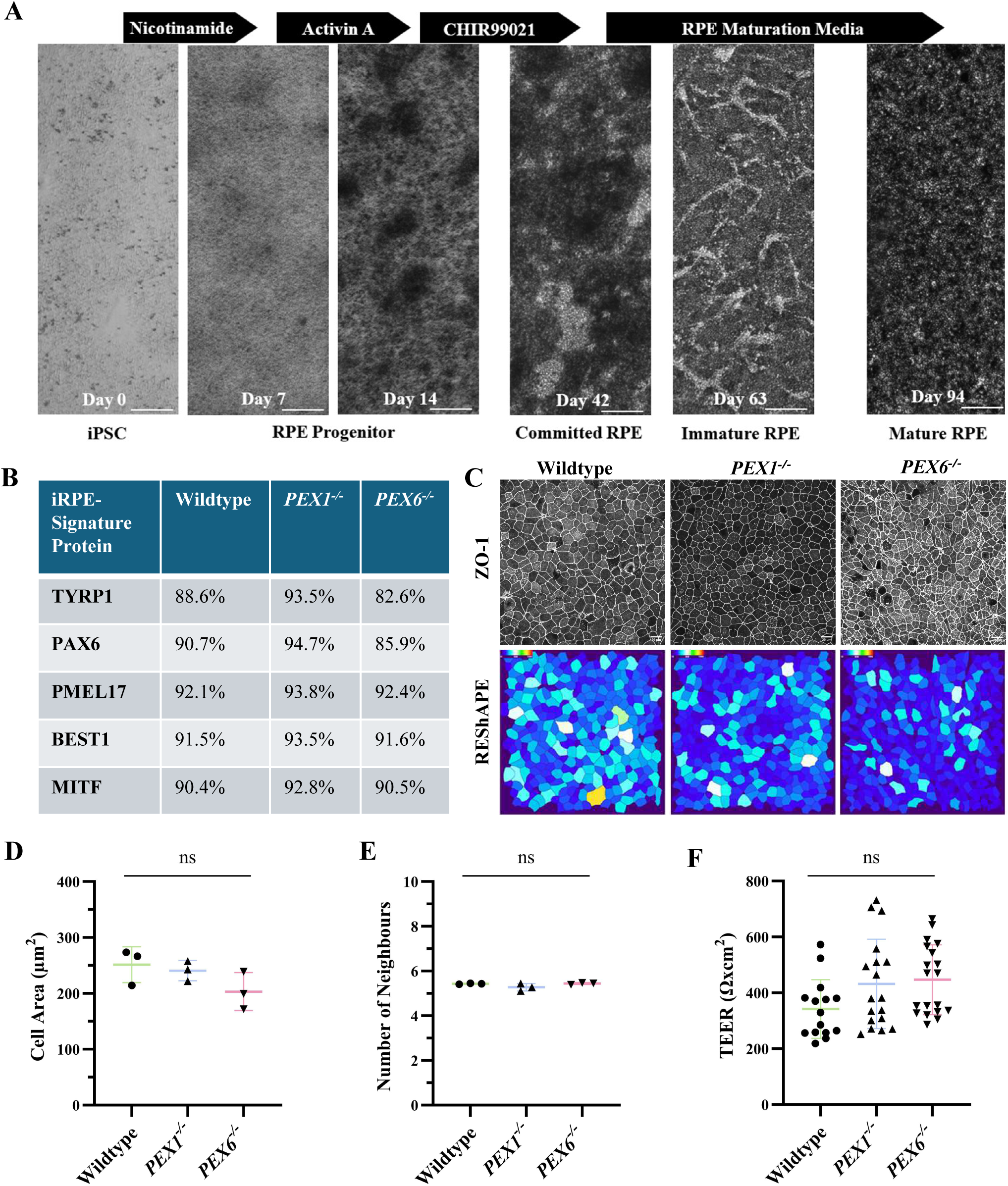
Differentiation and characterization of iPSC-RPE (iRPE) with PEX mutations. A. Representative brightfield images highlighting the differentiation timeline and media supplements. Scale bar = 350 μm. B. RPE signature proteins detected from iRPE populations assayed using flow cytometry. At least 10,000 events were captured for each sample. C. Representative confocal microscopy images of iRPE showing maximum intensity projections of anti-Zonula Occludens-1 (ZO-1), a tight junction protein, and corresponding REShAPE cell area heat maps. Scale bar = 20 μm. D. Mean (± standard deviation) iRPE area following REShAPE analysis. One-way ANOVA p=0.18 (n=3). E. Mean (± standard deviation) number of neighboring iRPE cells following REShAPE analysis. One-way ANOVA p=0.16 (n=3). F. Transepithelial electrical resistance (TEER) of wildtype (n=15), PEX1-/- (n=18), and PEX6-/- (n=19) iRPE cultured

### *PEX1^-/-^* and *PEX6^-/-^* iRPE Have Defective Import of Peroxisome Matrix Proteins

Knockout of PEX1 and PEX6 was confirmed by immunoblot using *PEX1^-/-^* and *PEX6^-/-^* iRPE lysates, respectively (Figures 2A and B). In addition, *PEX6^-/-^* iRPE had reduced PEX1 protein compared to wildtype iRPE, and PEX6 protein was not detected in *PEX1^-/-^* iRPE. PEX1 and PEX6 are tethered to the peroxisome membrane by PEX26 and facilitate the import of PTS1-and PTS2-containing cargo into the peroxisome matrix by enabling the recycling of the cargos’ cytosolic receptor, PEX5. Both PEX26 and PEX5 abundance were reduced in *PEX1^-/-^*and *PEX6^-/-^* iRPE compared to wildtype iRPE (Figures 2C and D). There was no significant difference in the number of peroxisomes per cell across all three iRPE lines assessed by quantifying peroxisomal membrane protein-70 (PMP70) abundance using immunoblotting and fluorescence microscopy (Figures 2E, F, and G).

**Figure 2:**
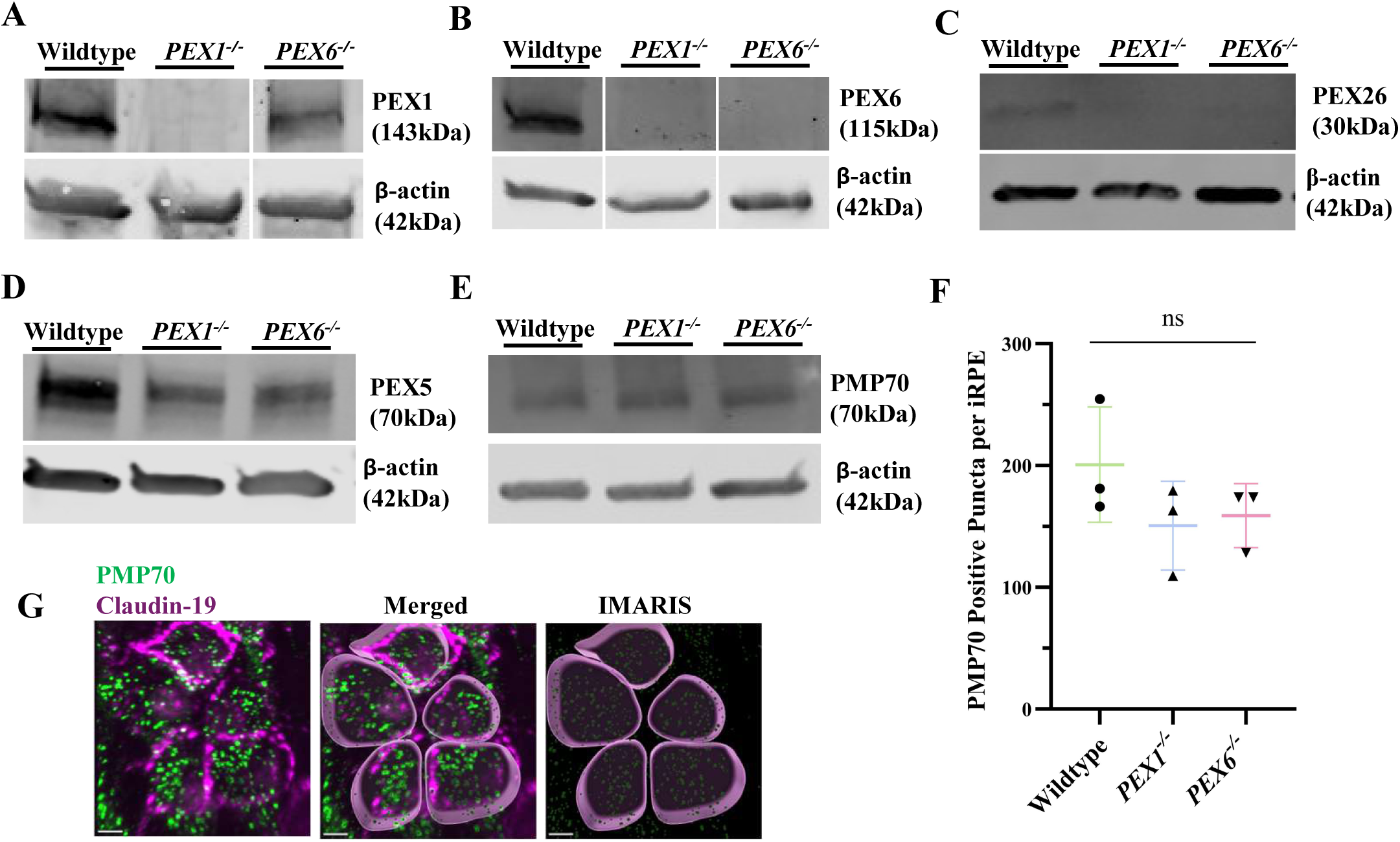
Consequences of knockout of PEX1 and PEX6 on peroxisome abundance and associated PEX proteins in iRPE. A. Representative immunoblot of iRPE lysates using anti-PEX1 and anti-β-actin antibodies. 50 μg of protein was loaded per lane (n=3). B. Representative immunoblot of iRPE lysates using anti-PEX6 and anti-β-actin antibodies. 50 μg of protein was loaded per lane (n=3). C. Representative immunoblot of iRPE lysates using anti-PEX26 and anti-β-actin antibodies. 30 μg of protein was loaded per lane (n=3). D. Representative immunoblot of iRPE lysates using anti-PEX5 and anti-β-actin antibodies. 30 μg of protein was loaded per lane (n=3). E. Representative immunoblot of iRPE lysates using anti-PMP70 and anti-β-actin antibodies. 30 μg of protein was loaded per lane (n=3). F. Mean (± standard deviation) number of PMP70 spots detected per iRPE. Three wells were analyzed per cell line. Five fields of view were randomly captured from each well, and 385 cells were analyzed in total. One-way ANOVA p=0.29 (n=3). G. Representative confocal microscopy images of iRPE showing IMARIS rendering used for the quantification of PMP70 positive puncta, a marker for peroxisomes. Scale bar = 5 μm.

To assess the integrity of peroxisome matrix protein import, *PEX1^-/-^, PEX6^-/-^,* and wildtype iPSCs were transfected with a GFP-PTS1 reporter (GFP-SKL). For this experiment, iPSCs were studied given their substantially higher transfection efficiency compared to post-mitotic iRPE.^13^ Forty-eight hours post-transfection, wildtype iPSCs displayed numerous intracellular GFP-PTS1-positive puncta, suggesting successful trafficking of GFP to peroxisomes (Figure 3A). However, both transfected *PEX1^-/-^* and *PEX6^-/-^* iPSCs demonstrated a diffuse cytosolic pattern of GFP fluorescence with no discernible GFP-PTS1-positive puncta, suggesting impaired GFP trafficking to peroxisomes (Figure 3A). The integrity of PTS1- and PTS2-mediated peroxisome matrix protein import was also assessed in iRPE by interrogating the proteolytic processing of peroxisomal β-oxidation enzymes, ACOX1, MFP2, and ACAA1 on an immunoblot. ACOX1 encodes acyl-CoA oxidase-1 and catalyzes the first and rate-limiting step of peroxisomal β-oxidation.^14,15^ ACOX1 is a 72-kDa protein and contains a C-terminal PTS1 sequence that targets the protein to the peroxisome. Once imported into the peroxisome matrix, ACOX1 undergoes proteolytic cleavage by the peroxisomal protease, TYSND1, into active 50-kDa and 22-kDa fragments.^16^ Wildtype iRPE, but not *PEX1^-/-^* and *PEX6^-/-^* iRPE, demonstrated a prominent 22-kDa ACOX1 band on immunoblot, suggesting impaired PTS1-mediated peroxisome matrix protein import in *PEX1^-/-^* and *PEX6^-/-^* iRPE (Figures 3B and C). MFP2 encodes multifunctional protein-2, an enzyme that catalyzes the second and third steps of peroxisomal β-oxidation.^17^ Similarly, once inside the peroxisome lumen, MFP2 is cleaved into an active form by TYSND1.^16^ Only wildtype iRPE demonstrated a prominent 45-kDa MFP2 band, representing the cleaved/processed form of the enzyme (Figures 3D and E).

**Figure 3:**
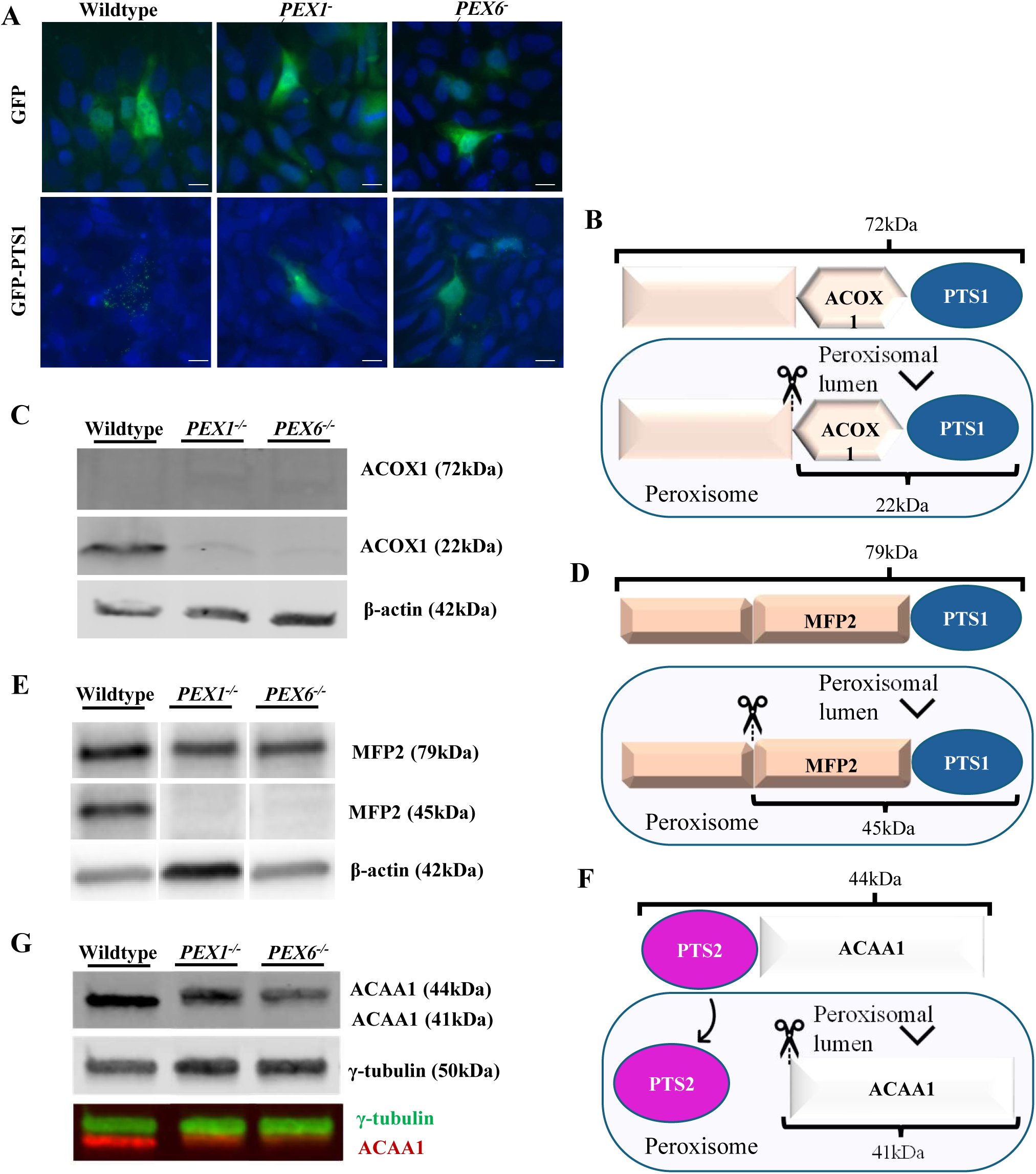
Knockout of PEX1 and PEX6 in iRPE results in peroxisomes with defective protein import. A. Representative widefield microscopy images of iPSCs transfected with GFP only or GFP-PTS1 expression plasmids. PTS1 targets proteins to the peroxisome matrix. Scale bar = 5μm. B. Schematic demonstrating the import of ACOX1 into peroxisomes, followed by its subsequent cleavage. C. Representative immunoblot of iRPE lysates using anti-ACOX1 and anti-β-actin antibodies. 50 μg of protein was loaded per lane (n=3). D. Schematic demonstrating the import of MFP2 into peroxisomes, followed by its subsequent cleavage. E. Representative immunoblot of iRPE lysates using anti-MFP2 and anti-β-actin antibodies. 10 μg of protein was loaded per lane (n=3). F. Schematic demonstrating the import of ACAA1 into peroxisomes, followed by its subsequent processing. G. Representative immunoblot of iRPE lysates using anti-ACAA1 and anti-γ-tubulin antibodies. 30 μg of protein was loaded per lane (n=3).

ACAA1 encodes acetyl-CoA acyltransferase-1 and catalyzes the final step of peroxisomal β-oxidation.^18,19^. ACAA1 is a 44-kDa protein that contains an N-terminal PTS2 sequence that is enzymatically cleaved in the peroxisome matrix by TYSND1.^16^ Similar to the ACOX1 and MFP2 results, wildtype iRPE, but not *PEX1^-/-^*and *PEX6^-/-^* iRPE, demonstrated a cleaved form of ACAA1 on immunoblot, suggesting impaired PTS2-mediated peroxisome matrix protein import in *PEX1^-/-^*and *PEX6^-/-^* iRPE (Figures 3F and G).

### Lipidomic Profiles of *PEX1^-/-^* and *PEX6^-/-^* iRPE Recapitulate Biochemical Abnormalities in PBDs

We performed lipidomic profiling of the three iRPE lines (n=3 replicates) using both gas and liquid chromatography-mass spectrometry approaches to elucidate specific lipid metabolic perturbations in iRPE with dysfunctional peroxisomes. Since plasmalogen synthesis requires functional peroxisomes, we measured phosphoethanolamine (PE)-containing plasmalogens,.^20^ Both *PEX1^-/-^* and *PEX6^-/-^* iRPE had reduced levels for all PE-plasmalogen species measured, C18:0(plasm)-Total-PE displayed 8.8-fold decrease (*p*=0.001) and 9.9-fold decrease (*p*=0.0026), respectively compared to wildtype iRPE (Figure 4A).

**Figure 4:**
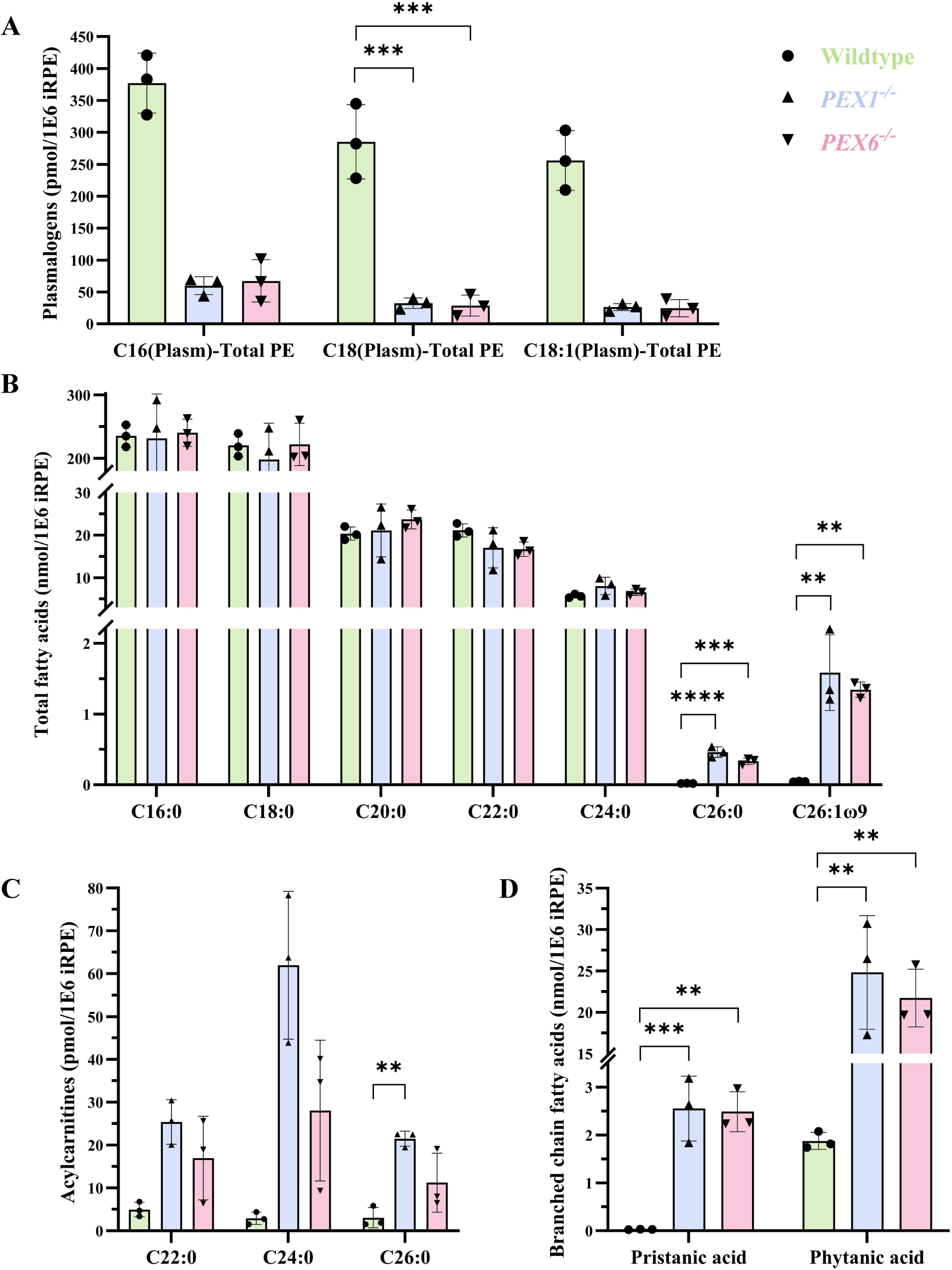
Quantitative analysis of fatty acids in PEX1 and PEX6 knockout iRPE at baseline. A. Quantification and comparison of plasmalogens in iRPE lysates measured by LC-MS. One-way ANOVA was performed on C18(Plasm)-Total PE only because the other species lacked specific standards; p=0.0002 (n=3). Dunnett’s multiple comparisons test was performed. B. Quantification and comparison of very long-chain fatty acids (VLCFA) in iRPE lysates measured by GC-MS. Dunnett’s multiple comparisons test was performed following a one-way ANOVA for each fatty acid (n=3). C. Quantification and comparison of acylcarnitines in iRPE lysates measured by LC-MS. One-way ANOVA was performed on C26:0 only because this was the only species with a standard available; p=0.0059 (n=3). Dunnett’s multiple comparisons test was performed. D. Quantification and comparison of branched chain fatty acids in iRPE lysates measured by GC-MS. Dunnett’s multiple comparisons test was performed following a one-way ANOVA for each fatty acid (n=3).

Clinically, VLCFAs are used as a biomarker for peroxisomal disorders^21^. *PEX1^-/-^* and *PEX6^-/-^*iRPE had significantly increased C26:0 compared to wildtype (24-fold increase; *p*=6.9x10^-5^ and 17-fold increase; *p*=0.0005, respectively) and dramatically increased C26:1ω9 (34-fold increase; *p*=0.0018 and 29-fold increase; *p*=0.0043, respectively) (Figure 4B). The C24:0 and C22:0 species were not significantly different across the three iRPE lines. In addition, there were no significant differences in any VLCFA species when comparing *PEX1^-/-^* and *PEX6^-/-^*iRPE. In general, the long-chain FAs C16:0 and C18:0 were approximately one order of magnitude more abundant than the VLCFA species in iRPE. There were no significant differences in C16:0 and C18:0 levels across all three iRPE lines (Figure 4B). Compared to wildtype, the ratio of C26:0 to C22:0, a biochemical diagnostic test for patients with PBDs, was 27-fold higher in *PEX1^-/-^* iRPE and 19-fold higher in *PEX6^-/-^* iRPE. Similarly, the ratio of C24:0 to C22:0, another biochemical PBD test, was 1.8-fold higher in *PEX1^-/-^*iRPE and 1.5-fold higher in *PEX6^-/-^* iRPE compared to wildtype. Additionally, C26:0 acylcarnitine species were 7.0-fold higher (*p*=0.0036) in *PEX1^-/-^*iRPE relative to wildtype (Figure 4C). While C26:0 acylcarnitine species were 3.7-fold higher in *PEX6^-/-^* iRPE relative to wildtype, this did not reach statistical significance (*p*=0.10).

The branched-chain fatty acids (BCFAs) pristanic acid and phytanic acid require intact peroxisome function for their metabolism by β-oxidation and α-oxidation, respectively.^22,23^ For pristanic acid, in comparison to wildtype iRPE, *PEX1^-/-^* iRPE had a 96.9-fold increased level (*p*=0.0009) and *PEX6^-/-^* iRPE had a 94.4-fold increased level (*p*=0.0011). For phytanic acid, relative to wildtype iRPE, *PEX1^-/-^* iRPE had a 13.2-fold increased level (*p*=0.0013) and *PEX6^-/-^* iRPE had a 11.6-fold increased level (*p*=0.0028). There were no significant differences in BCFA levels between *PEX1^-/-^* and *PEX6^-/-^* iRPE (Figure 4D).

### Intracellular Lipid Accumulates in *PEX1^-/-^* and *PEX6^-/-^*iRPE

Given the essential role of peroxisomes in lipid metabolism, we investigated whether intracellular lipids accumulate in *PEX1^-/-^* and *PEX6^-/-^* iRPE relative to wildtype iRPE. *PEX1^-/-^*iRPE had a 1.5-fold greater LipidTOX Deep Red Mean Fluorescence Intensity (MFI) relative to wildtype iRPE measured by flow cytometry, indicating increased intracellular neutral lipids (*p*=0.018). Similarly, *PEX6^-/-^* iRPE had a 1.8-fold greater LipidTOX Deep Red MFI relative to wildtype iRPE (*p*=0.0025), suggesting increased intracellular neutral lipids (Figure 5A). Both immunofluorescence microscopy and immunoblotting using antibodies against perilipin-2 (PLIN2), one of the most abundant lipid droplet (LD)-related proteins,^24^ revealed increased PLIN2 in *PEX1^-/-^*and *PEX6^-/-^* iRPE relative to wildtype at baseline (Figures 5B, C, and D). By immunoblot analysis, *PEX1^-/-^* and *PEX6^-/-^* iRPE had 1.8-fold (*p=*0.038) and 2.3- fold (*p*=0.0009) increases in PLIN2, respectively, compared to wildtype iRPE at baseline (Figures 5B and D). Next, all three iRPE lines were subjected to a 1, 12, or 24-hour continuous POS challenge (0.5 POS per cell). After the 1-hour POS challenge, PLIN2 increased in wildtype, *PEX1^-/-^* and *PEX6^-/-^* iRPE. However, relative to PLIN2 levels at 1 hour, wildtype iRPE demonstrated a 34% reduction in PLIN2 at 12 hours, whereas *PEX1^-/-^* and *PEX6^-/-^* iRPE demonstrated only 13% and 11% reductions in PLIN2 at 12 hours, respectively. At 24 hours, PLIN2 levels remained persistently higher in the *PEX1^-/-^* (1.9-fold increase; p=0.029) and *PEX6^-/-^*iRPE (2.5-fold increase; *p*=0.0006) relative to wildtype (Figures 5D and E).

**Figure 5:**
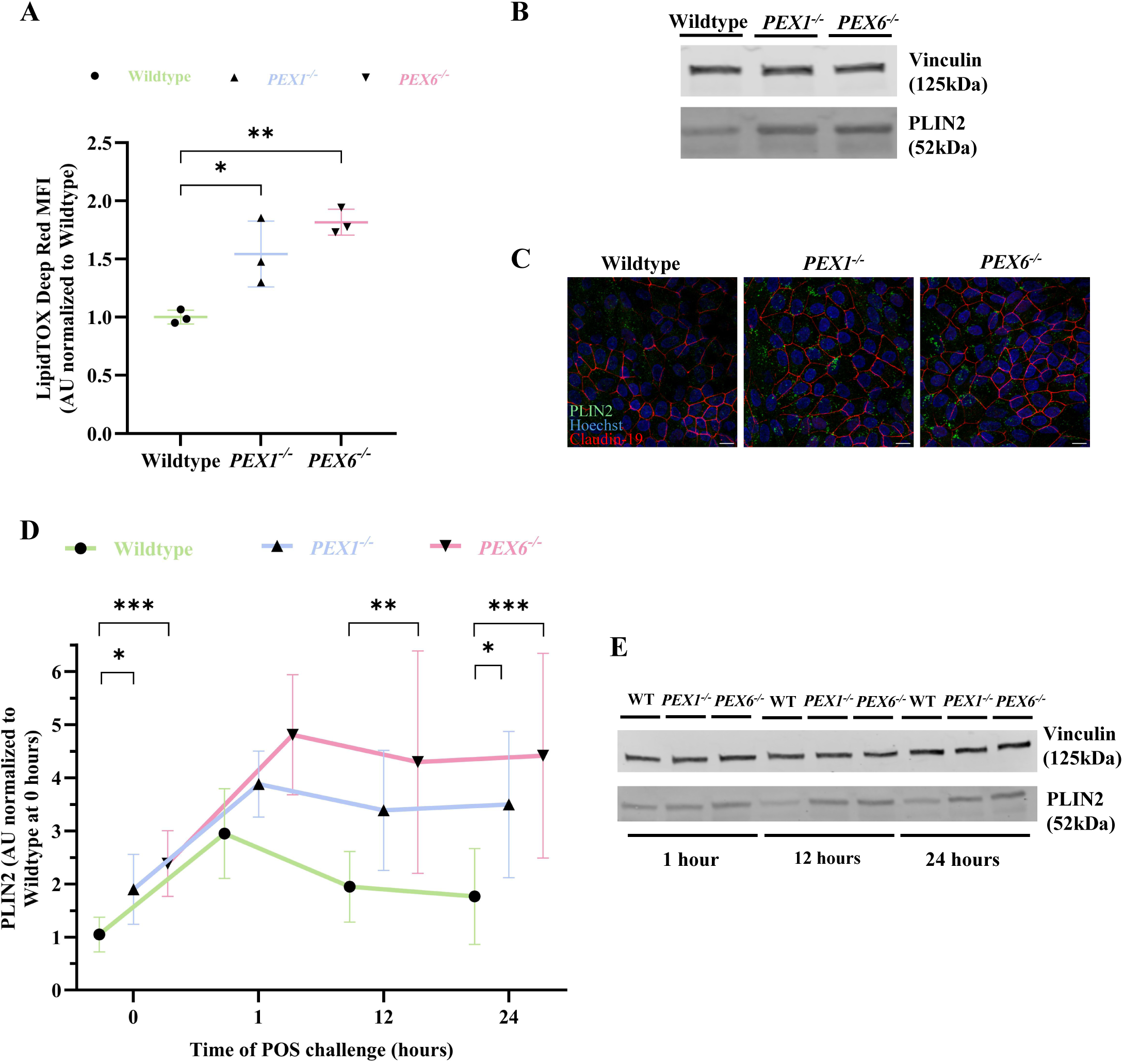
Lipid droplet perturbations in PEX1 and PEX6 knockout iRPE. A. Mean fluorescence intensity (MFI) of LipidTOX Deep Red, a neutral lipid dye, in iRPE captured by flow cytometry. At least 100,000 events were captured for each sample. One-way ANOVA p=0.0038 (n=3). Dunnett’s multiple comparisons test was performed. B. Representative immunoblot of baseline lipid droplets in iRPE lysates using anti-PLIN2, a lipid droplet protein. 30 μg of protein was loaded per lane (n=6). C. Representative confocal microscopy images of iRPE showing maximum intensity projections of Hoechst, PLIN2, and Claudin-19, a tight junction protein, at baseline. Scale bar = 10 μm. D. Immunoblot quantification of PLIN2 densitometry (normalized to vinculin) in iRPE lysates following a continuous POS challenge (0.5 POS per cell). Two-way ANOVA yielded p=6.7x10-6 for genotype variation, and p=6.3x10-7 for time variation (n=6). Dunnett’s multiple comparisons test was performed. E. Representative immunoblot of lipid droplets (anti-PLIN2) from iRPE lysates following a continuous POS challenge (0.5 POS per cell). 30 μg of protein was loaded per lane (n=6). *p<0.05; **p<0.01; ****p<0.0001

### *PEX1^-/-^* and *PEX6^-/-^* iRPE Have Reduced DHA and Impaired Retroconversion of ω3 and ω6 PUFAs

To evaluate the effect of peroxisome dysfunction on the metabolism of specific lipid species present in POS, we measured the lipid profiles of the three iRPE lines before and after a POS challenge using GC-MS. First, we identified and quantified the FA lipid species present in isolated bovine POS. The most abundant PUFA lipid species was DHA (C22:6ω3) at 151.8 pmol/μg of total protein, representing 36.6% of all measured FA lipid species (Figure 6A). The second and third most abundant PUFAs were C22:5ω6 (DPAω6) (41.5 pmol/μg of total protein) and C20:4ω6 (AA) (26.3 pmol/μg of total protein), respectively. EPA (C20:5ω3) existed in only trace amounts in bovine POS. The most abundant saturated FA lipid species were C18:0 (99.8 pmol/μg of total protein) and C16:0 (75.4 pmol/μg of total protein) (Figure 6A). A comprehensive dataset of measured lipid species in bovine POS is available in Supplementary Table 2.

**Figure 6:**
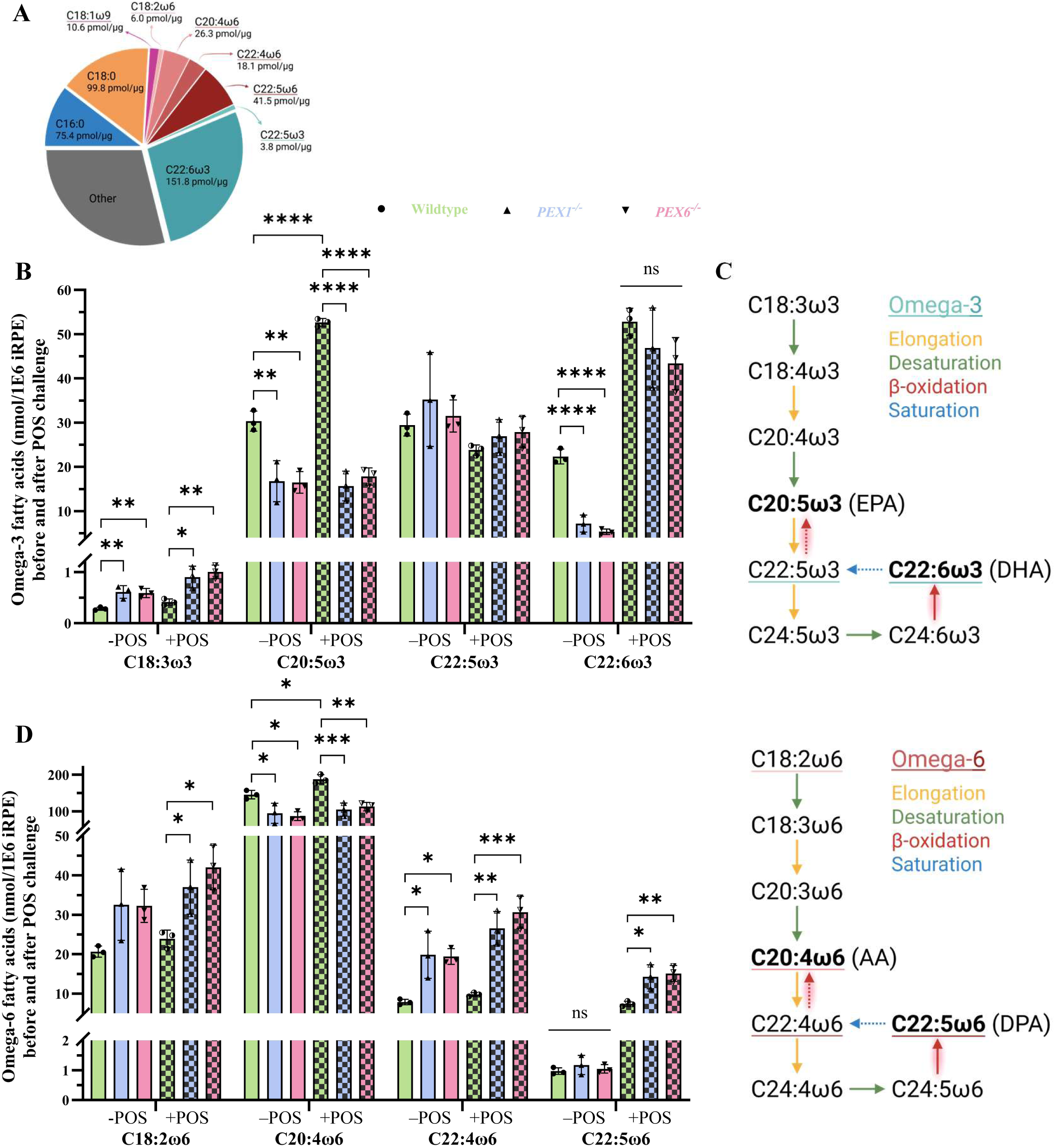
Quantitative analysis of fatty acid profile changes in iRPE following a POS challenge. A. The most abundant fatty acid species detected by GC-MS from isolated bovine POS (n=3). Refer to Supplementary Table 2 for a complete list of detected fatty acid species. B. Quantification of omega-3 (ω3) fatty acids in iRPE lysates by GC-MS before and after a 7-day POS challenge (3 POS per cell were added with daily media changes). Dunnett’s multiple comparisons test was performed following a one-way ANOVA for each fatty acid (n=3). C. The omega-3 fatty acid metabolic pathway adapted from Yu et al., 2012. Reactions that are presumed to require intact peroxisome β-oxidation are highlighted in red. D. Quantification of omega-6 (ω6) fatty acids in iRPE lysates by GC-MS before and after a 7-day POS challenge (3 POS per cell were added with daily media changes). Dunnett’s multiple comparisons test was performed following a one-way ANOVA for each fatty acid (n=3). E. The omega-6 fatty acid metabolic pathway adapted from Yu et al., 2012. Reactions that are presumed to require in_4_ta_6_ct peroxisome β-oxidation are highlighted in red. *p<0.05; **p<0.01; ***p<0.001; ****p<0.0001

The ω3 PUFAs are essential lipids since they cannot be synthesized *de novo* in humans, yet they are highly abundant in the brain and retina^25^. These lipids have critical roles in POS membrane structure and support phototransduction, and they impart cellular anti-inflammatory, anti-oxidative, and neuroprotective properties.^26^ The biosynthesis of ω3 PUFAs begins with dietary α-linolenic acid (ALA; C18:3ω3) and proceeds through a series of elongation and desaturation reactions to generate EPA (C20:5ω3) and DHA (C22:6ω3). Both DHA and its precursor, C24:6ω3, can be retroconverted to EPA through a series of β-oxidation steps that occur in peroxisomes (Figure 6C).^27^ The relative contribution of the retroconversion pathway for generating intracellular EPA pools has been debated, and it has not been studied in the RPE in the context of peroxisome dysfunction.^28–30^ To address this, we measured ω3 PUFAs before and after a 7-day POS challenge (3 POS per iRPE cell were added with daily media changes) in all three iRPE lines.

At baseline, *PEX1^-/-^* and *PEX6^-/-^* iRPE had significantly elevated ALA relative to wildtype iRPE (2.2-fold increase (*p*=0.0073) and 2.1-fold increase (*p*=0.0098), respectively). In contrast, *PEX1^-/-^* and *PEX6^-/-^* iRPE had significantly reduced EPA (1.8-fold decrease (*p*=0.0043) and 1.8-fold decrease (*p*=0.0038), respectively) and DHA (3.1-fold decrease (*p*=1.8x10^-5^) and 4.2-fold decrease (*p*=4.6x10^-6^), respectively) relative to wildtype iRPE (Figure 6B). Following the POS challenge, DHA significantly increased across all iRPE lines relative to baseline. Despite only trace amounts of EPA (and other ω3 species aside from DHA) in bovine POS, there was a 1.7-fold increase (*p*=4.0x10^-5^) in EPA, the retroconversion product of DHA, post-POS challenge in wildtype iRPE. In stark contrast, *PEX1^-/-^* and *PEX6^-/-^*iRPE did not demonstrate an increase in EPA following the POS challenge (Figure 6B). To determine whether the increase in EPA in wildtype iRPE, but not *PEX1^-/-^* and *PEX6^-/-^* iRPE, was from the DHA in POS, we challenged all three iRPE lines with 30 uM DHA conjugated to BSA. We observed the same result as the POS challenge, suggesting impaired DHA retroconversion in *PEX1^-/-^* and *PEX6^-/-^* iRPE (Supplementary Figures 3A and B).

We simultaneously evaluated changes in the ω6 PUFA species in all three iRPE lines before and after a POS challenge. The metabolic pathway for the ω6 PUFAs mirrors that of the ω3 PUFAs, and they compete for the same set of enzymes (Figure 6E).^31^ Consistent with the ω3 PUFAs results, following the POS challenge, all three iRPE lines accumulated C22:5ω6 (DPAω6), which is the most common of the ω6 PUFA detected in bovine POS and is the ω6 PUFA equivalent of DHA (Figure 6D). However, the *PEX1^-/-^* and *PEX6^-/-^* iRPE accumulated approximately twice the amount of C22:5ω6 relative to wildtype iRPE following the POS challenge (*p*=0.014 and *p*=0.0082, respectively). In addition, wildtype iRPE had a 1.3-fold increase (*p*=0.041) in arachidonic acid (C20:4ω6), the ω6 PUFA equivalent of EPA and retroconversion product of C22:5ω6, following a POS challenge. Both *PEX1^-/-^* and *PEX6^-/-^*iRPE failed to significantly accumulate C20:4ω6 following a POS challenge (Figure 6D). Similar to the ω3 PUFA species, we challenged all three iRPE lines with 30 uM DPAω6 conjugated to BSA and observed the same result as the POS challenge, suggesting impaired DPAω6 retroconversion in *PEX1^-/-^* and *PEX6^-/-^*iRPE (Supplementary Figures 3C and D).

### *PEX1^-/-^* and *PEX6^-/-^* iRPE Have Disrupted Phagocytosis of Photoreceptor Outer Segments

Emerging evidence suggests a critical role of peroxisomes in phagocytosis, especially for immune cells.^32,33^ To determine the consequence of peroxisome dysfunction on phagocytosis in the RPE, the presence of rhodopsin, a protein that is not expressed in the RPE but is the most abundant protein in POS, was evaluated using an antibody recognizing an epitope on the N-terminus of the protein. Since alterations in phagocytosis may be attributed to differences in POS uptake, degradation, or a combination of both, we first utilized an approach that more broadly measures total POS consumption by the RPE called the total RPE consumption capacity.^34^ This assay measures the amount of POS persisting both inside the RPE (engulfed but not degraded) and within the media (not engulfed) by rhodopsin immunoblotting of harvested cells and media together after a continuous POS challenge.^34^.

Following a 1-hour POS challenge, there was no significant difference between total rhodopsin (iRPE lysates and media) in wildtype, *PEX1^-/-^* or *PEX6^-/-^* iRPE (Figures 7A and B). However, by 24 hours, wildtype iRPE had the largest reduction in rhodopsin, with only 19.0% of initial rhodopsin remaining, compared to 32.7% and 38.9% of initial rhodopsin remaining in *PEX1^-/-^* and *PEX6^-/-^* iRPE, respectively. By 24 hours, *PEX1^-/-^* and *PEX6^-/-^* iRPE had 1.7-fold higher (*p*=0.032) and 1.9-fold higher (*p*=0.0026) levels of rhodopsin relative to wildtype respectively, suggesting impaired rhodopsin degradation and disrupted POS phagocytosis in *PEX1^-/-^* and *PEX6^-/-^* iRPE (Figures 7A and B).

**Figure 7:**
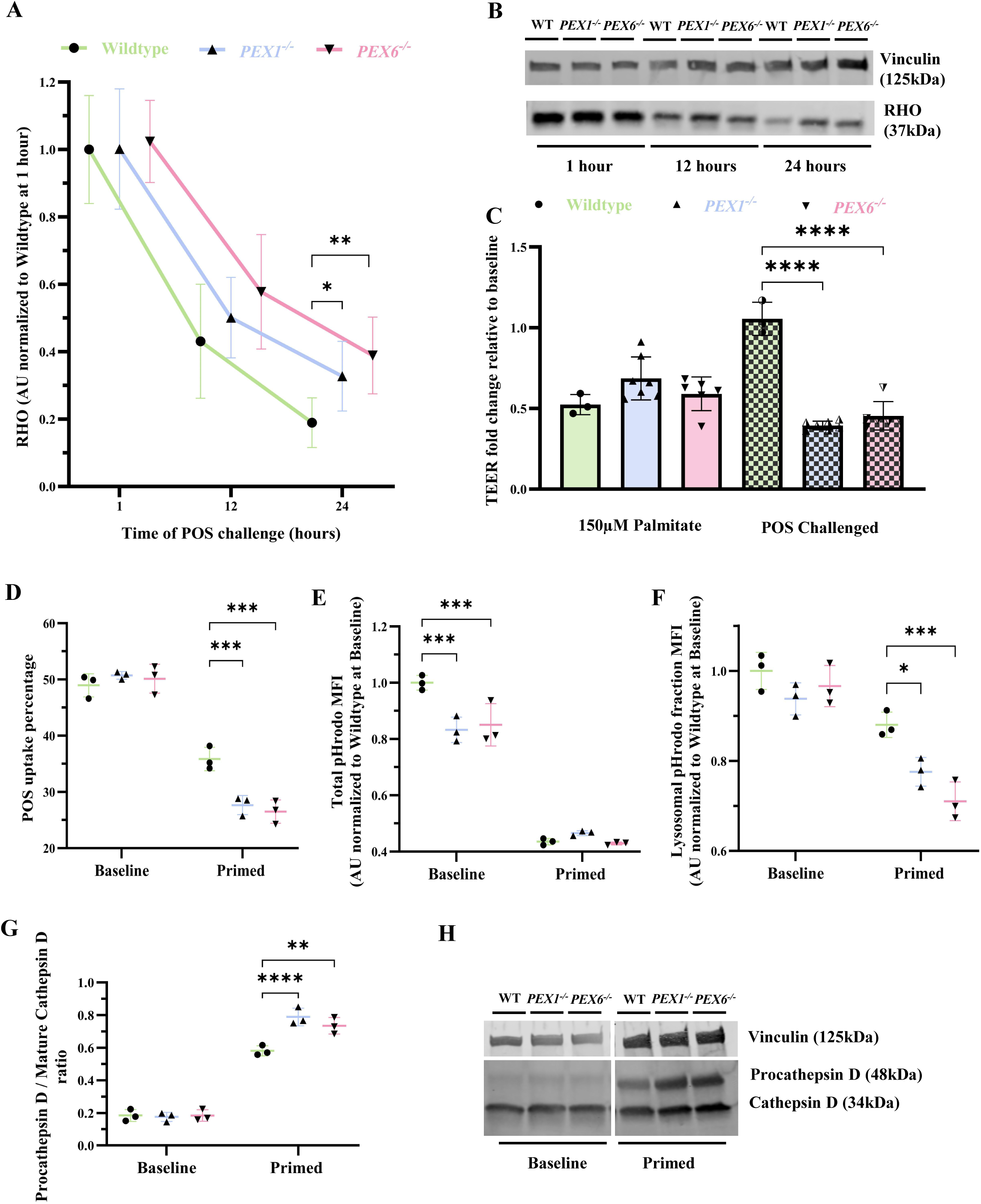
Interrogating POS phagocytosis in PEX1 and PEX6 knockout iRPE. A. Quantification of changes in RHO densitometry by immunoblot in iRPE lysates following a continuous POS challenge (0.5 POS per cell). Two-way ANOVA yielded p=0.021 for genotype and p=2.1x10-22 for time variations (n=7). Dunnett’s multiple comparisons test was performed. B. Representative immunoblots of iRPE lysates probed with anti-rhodopsin (RHO) and anti-vinculin antibodies (loading control) following a continuous POS challenge (0.5 POS per cell) (n=7). C. Relative fold change (compared to baseline) in transepithelial electrical resistance (TEER) of iRPE cultured on 0.33cm2 transwells following a 24-hour incubation with 150 µM palmitate or POS (10 POS per cell). Two-way ANOVA yielded p=1.2x10-5 for genotype variation, and p=1.0x10-8 for the interaction (n=7). Dunnett’s multiple comparisons test was performed. D. Frequency of live iRPE cells that were pHrodo-positive following a 1.5 pHrodo-labeled POS per cell challenge over one hour, measured by flow cytometry. At least 100,000 events were captured for each sample. Cells were primed with 3 unlabeled POS per cell added with daily media changes over 7 days. Two-way ANOVA yielded p=0.0083 for genotype variation, p=5.4x10-11 for the priming, and p=0.0008 for the interaction (n=3). Dunnett’s multiple comparisons test was performed. E. Mean fluorescence intensity (MFI) of pHrodo-labeled POS internalized by live iRPE and measured by flow cytometry. At least 100,000 events were captured for each sample. Two-way ANOVA yielded p=0.0077 for genotype variation, p=9.5x10-12 for priming, and p=0.0021 for the interaction (n=3). Dunnett’s multiple comparisons test was performed. F. Mean fluorescence intensity (MFI) of pHrodo-labeled POS that reached acidic environments in live iRPE and measured by flow cytometry. At least 100,000 events were captured for each sample. Two-way ANOVA yielded p=0.0013 for genotype variation, p=3.7x10-7 for priming, and p=0.026 for the interaction (n=3). Dunnett’s multiple comparisons test was performed. G. The ratio of procathepsin D to mature cathepsin D densitometry in iRPE lysates following a one-hour challenge (1.5 POS per cell) before (baseline) or after priming (3 POS per cell added with daily media changes over 7 days). Two-way ANOVA yielded p=6.7x10-6 for genotype variation and p=6.3x10-7 for time variation (n=3). Dunnett’s multiple comparisons test was performed. H. Representative immunoblots of iRPE lysates probed with anti-cathepsin D and anti-vinculin (loading control) antibodies following a one-hour challenge (1.5 POS per cell) before (baseline) or after priming (3 POS per cell added with daily media changes over 7 days) (n=3). *p<0.05; **p<0.01; ***p<0.001; ****p<0.0001

To assess the effect of a POS challenge on RPE integrity, we measured transepithelial electrical resistance (TEER) of the iRPE lines following a 24-hour POS challenge. Incubating the iRPE for 24 hours with 150 μM of palmitate, a lipotoxic concentration of the fatty acid that reduces cellular metabolic activity,^35^ resulted in an approximately 2-fold reduction in TEER from baseline across all three iRPE lines (Figure 7C). Following a 24-hour POS challenge, there was no change in TEER in wildtype iRPE compared to baseline. However, following a 24-hour POS challenge, *PEX1^-/-^*iRPE had a 2.5-fold reduction in TEER compared to baseline (*p*=2.0x10^-9^). Similarly, *PEX6^-/-^* iRPE had a 2.2-fold reduction in TEER following a 24-hour POS challenge compared to baseline (*p*=7.2x10^-9^) (Figure 7C).

To elucidate whether impaired rhodopsin degradation in *PEX1^-/-^* and *PEX6^-/-^* iRPE was due to impaired POS uptake and/or compromised trafficking through the phagolysosomal pathway, we incubated iRPE with pHrodo-labeled POS (pHrodo-POS), a pH-sensitive dye that fluoresces more intensely in acidic environments, for 1 hour and measured MFI by flow cytometry at baseline and following a 7-day challenge (3 unlabeled POS/cell delivered daily). The 7-day POS challenge generated “primed” iRPE that recapitulated the continuous POS burden encountered by RPE in vivo. At baseline, approximately 50% of iRPE in each line were positive for pHrodo-POS after 1 hour, implying similar POS uptake (*p*=0.56) (Figure 7D). However, the total pHrodo-POS MFI in *PEX1^-/-^* and *PEX6^-/-^*iRPE was 16.7% (*p*=0.0003) and 15.0% (*p*=0.0008) lower than wildtype, respectively, suggesting a deficiency in the trafficking of pHrodo-POS to acidic intracellular compartments (Figure 7E). Following the 7-day unlabeled POS challenge, all iRPE lines demonstrated a substantial reduction in the percent of cells positive for pHrodo-POS after 1 hour; however, *PEX1^-/-^* and *PEX6^-/-^*iRPE had 22.8% (*p*=0.0005) and 26.2% (*p*=0.0001) fewer pHrodo-POS positive cells compared to wildtype at this timepoint, respectively, implying a reduced capacity for POS uptake in primed *PEX1^-/-^* and *PEX6^-/-^* iRPE (Figure 7D). The total pHrodo-POS MFI following the 7-day POS challenge was substantially reduced from baseline, but similar across all three iRPE lines, likely resulting from an already congested phagolysosomal pathway with unlabeled POS (Figure 7E). Finally, we utilized a gating strategy developed with bafilomycin, a drug that blocks the lysosomal V-ATPase and prevents lysosomal acidification, to interrogate the pHrodo-POS MFI in a fluorescence fraction corresponding to the terminal compartments of the phagolysosomal pathway. For the purposes of this assay, we termed this fraction the lysosomal fraction (Supplementary Figure 5). At baseline, there was no difference in pHrodo-POS MFI in the lysosomal fraction across all three iRPE lines (*p*=0.26); however, following the 7-day POS challenge, *PEX1^-/-^* and *PEX6^-/-^* iRPE had 11.9% (*p*=0.011) and 19.3% (*p*=0.0003) reduction in pHrodo-POS MFI in the lysosomal fraction, respectively, reflecting a POS uptake deficiency and/or suggesting impaired lysosomal acidification compared to wildtype (Figure 7F).

Finally, we probed the maturation of cathepsin D, the major lysosomal protease for rhodopsin degradation in the RPE.^36,37^ Cathepsin D undergoes a series of post-translational modifications and proteolytic cleavages that culminates in the generation of mature and enzymatically active cathepsin D (14 kDa and 34 kDa fragments) from procathepsin D (48 kDa) in lysosomes.^38^ Following the same pHrodo-POS challenge, the ratio of procathepsin D to mature cathepsin D was the same across all iRPE lines at baseline (Figures 7G and H).

However, after the same 7-day POS challenge, *PEX1^-/-^*and *PEX6^-/-^* iRPE had a 1.4-fold (*p*=1.5x10^-7^) and a 1.3-fold (*p*=0.001) increase, respectively, in the ratio of procathepsin D to mature cathepsin D compared to wildtype, indicating a relative impairment in the maturation of a critical lysosomal protease (Figures 7G and H).

Together, our data suggest that *PEX1^-/-^* and *PEX6^-/-^*iRPE have altered phagocytosis evident by the disrupted processing of POS, likely due to phagolysosomal trafficking defects, altered lysosomal function, and secondary perturbation of POS uptake, which result in a reduction in RPE cellular health, as assessed by TEER.

## Discussion

We have developed the first human iPSC-derived RPE models to study the effects of peroxisome dysfunction in the RPE, an essential retinal layer that is disrupted in patients with PBDs.^39,40^ The *PEX1^-/-^* and *PEX6^-/-^* iRPE recapitulated features of differentiated RPE as demonstrated by the development of pigmentation, hexagonal morphology with tight junctions, transepithelial electrical resistance, and expression of RPE signature proteins (Figure 1). These properties closely resemble those of other iRPE models and cultured primary human RPE.^41–43^ We did not appreciate any differences in the development of these RPE features when comparing *PEX1^-/-^* and *PEX6^-/-^* iRPE to wildtype iRPE, suggesting that intact peroxisome function may not be essential for iPSC generation and RPE maturation *in vitro*.

As expected, the *PEX1^-/-^* and *PEX6^-/-^* iRPE demonstrated impaired proteolytic processing of ACOX1, MFP2, and ACAA1, suggesting that these enzymes did not effectively reach the peroxisome matrix where they are normally cleaved (Figure 3).^16^ Moreover, GFP-tagged SKL, the C-terminal PTS1 tripeptide that targets proteins to the peroxisome matrix, was mislocalized to the cytosol in *PEX1^-/-^* and *PEX6^-/-^* iPSCs, confirming a global defect in matrix protein import (Figure 3). Consistent with studies in fibroblasts from patients with *PEX1*-, *PEX6*-, and *PEX26*-related PBDs demonstrating reduced PEX5 protein abundance,^44,45^ the *PEX1^-/-^* and *PEX6^-/-^* iRPE also had decreased PEX5 protein compared to wildtype iRPE (Figure 2), likely due to a quality control mechanism that reduces the accumulation of inoperable PEX5 on peroxisome membranes.^46^ Interestingly, knockout of either PEX1 or PEX6 reduced the abundance of the other protein, an effect especially pronounced in *PEX6^-/-^*iRPE where PEX1 was not detectable (Figure 2). This finding is in agreement with a recent study that demonstrated reduced PEX1 when PEX6 was knocked-down in HCT116 cells,^47^ and likely occurs due to the process by which cells degrade unpaired subunits in protein complexes.^48^ While the import of matrix proteins was impaired in *PEX1^-/-^* and *PEX6^-/-^* iRPE, the number of peroxisomes per cell was not significantly different from wildtype iRPE (Figure 2). This differs from the reduced peroxisome abundance reported in *PEX6^-/-^*HEK293T cells^39^ and *PEX1^-/-^* patient fibroblasts,^49^ where the latter study mechanistically linked reduced peroxisome abundance to increased peroxisome degradation by pexophagy.^49^ We suspect that the effect of *PEX1* and *PEX6* mutation on peroxisome abundance could be cell-type specific, and the highly metabolically active RPE may upregulate peroxisome biogenesis in an attempt to offset peroxisome degradation and dysfunction.

Lipid biochemical analyses of iRPE confirmed significantly elevated VLCFAs, BCFAs, and reduced plasmalogens in *PEX1^-/-^* and *PEX6^-/-^* iRPE compared to wildtype iRPE (Figure 4), recapitulating lipid abnormalities in both plasma and dermal fibroblasts from patients with severe PBDs^50–52^ and in the RPE of *Pex1*-p.G844D mutant mice.^53^ Elevated VLCFAs implies impaired peroxisomal β-oxidation in *PEX1^-/-^* and *PEX6^-/-^* iRPE, consistent with the perturbed proteolytic processing of ACOX1, MFP2, and ACAA1, three essential enzymes in peroxisomal β-oxidation (Figure 3). Intriguingly, the *PEX1^-/-^*and *PEX6^-/-^* iRPE each demonstrated a nearly 100-fold increase in pristanic acid and more than 10-fold increase in phytanic acid compared to wildtype iRPE, despite the relatively low abundance of these BCFAs in FBS-containing media.^54^ Patients with PBDs typically do not demonstrate such a profound increase in BCFAs in their plasma.^50,52^ The accumulation of phytanic acid and pristanic acid in *PEX1^-/-^* and *PEX6^-/-^* iRPE may contribute to the retinal degeneration phenotype in patients with PBDs. In support of this hypothesis, patients with adult Refsum disease accumulate massive amounts of phytanic acid due to loss-of-function of the peroxisomal enzyme phytanoyl-CoA hydroxylase and develop retinal degeneration that resembles that seen in patients with PBDs.^55^ Both pristanic acid and phytanic acid induce mitochondrial dysfunction and disrupt calcium homeostasis in rat neurons and glia.^56^ In primary human RPE, phytanic acid accumulation leads to ultrastructural disturbances including vacuolation and loss of apical microvilli.^57^

One of the major roles of the RPE in maintaining normal vision is the continuous phagocytosis and metabolism of POS. The POS are enriched in very long-chain-PUFAs (VLC-PUFAs) and various long-chain PUFAs (LC-PUFAs), including DHA, and their daily phagocytosis places a large metabolic burden on the RPE.^58^ We identified an accumulation of LipidTOX Deep Red-positive intracellular neutral lipids in *PEX1^-/-^* and *PEX6^-/-^* iRPE compared to wildtype iRPE at baseline, which correlated with PLIN2-positive lipid droplet formation before and after a POS challenge (Figure 5). Since peroxisomes import and oxidize fatty acids released from lipid droplets,^59^ differences in neutral lipid accumulation in *PEX1^-/-^*and *PEX6^-/-^* iRPE compared to wildtype iRPE at baseline could be due to compromised peroxisome-lipid droplet interactions.^60^ Following a POS challenge in *PEX1^-/-^*and *PEX6^-/-^*iRPE, fatty acids that would normally be metabolized by peroxisomes may instead accumulate in the RPE and become esterified and subsequently stored in lipid droplets.

In addition, a reduced total RPE consumption capacity for rhodopsin was observed in *PEX1^-/-^*and *PEX6^-/-^* iRPE compared to wildtype, and a reduction in RPE cellular health demonstrated by reduced TEER in *PEX1^-/-^* and *PEX6^-/-^* iRPE (Figure 7). Interestingly, while the uptake of POS at baseline was similar across all three iRPE lines, there was a significant reduction in the proportion of *PEX1^-/-^* and *PEX6^-/-^* iRPE engulfing POS relative to wildtype following a 7-day POS challenge. This coincided with an apparent defect in the trafficking of POS to terminal phagolysosomal compartments and evidence of lysosomal dysfunction (Figure 7).

Peroxisomal phosphatidylinositol 4,5-bisphosphate (PI(4,5)P2) and lysosomal synaptotagmin VII interact to establish membrane contact sites and facilitate the transfer of cholesterol from lysosomes to peroxisomes.^61^ Given that *PEX1* and *PEX6* knockdown in HeLa cells reduces peroxisomal PI(4,5)P2, and cholesterol accumulates in PBD patient fibroblasts,^61^ *PEX1^-/-^* and *PEX6^-/-^* iRPE may have disrupted peroxisome-lysosome interactions leading to lipid accumulation and lysosomal dysfunction, which may contribute to the impairment of POS uptake following a continuous POS challenge. Kocherlakota and colleagues identified lipid droplet accumulation and lysosomal dysfunction in the RPE of both global and RPE-specific *Mfp2*^-/-^ mice,^4^ supporting this hypothesis. Our iRPE models offer a platform to further investigate the coordination and regulation of peroxisomes, lipid droplets, and lysosomes in the RPE.

While RPE degeneration occurs in patients with biallelic pathogenic loss-of-function variants in *PEX1* and *PEX6*,^39,40^ our *PEX1^-/-^* and *PEX6^-/-^* iRPE did not demonstrate any overt signs of degeneration at baseline. However, when these iRPE were burdened with a physiological lipid load in the form of POS, both *PEX1^-/-^* and *PEX6^-/-^* iRPE had a greater than 50% reduction in TEER, indicating disrupted integrity of the RPE barrier and reduced cellular health.^62^ We hypothesize that the *PEX1^-/-^*and *PEX6^-/-^* iRPE may have altered metabolic pathways that compensate for peroxisome dysfunction, but that these processes become overwhelmed during POS phagocytosis. Intriguingly, while an RPE-specific deletion of *Mfp2* led to RPE dysfunction and secondary photoreceptor degeneration in mice, RPE degeneration did not occur in *Mfp2*/*rd1* double mutants that do not develop POS.^4^ Consistent with our *PEX1^-/-^*and *PEX6^-/-^* iRPE, this implies that the RPE is highly dependent on β-oxidation reactions housed inside functional peroxisomes for POS phagocytosis.

DHA is the most abundant ω3 PUFA in the retina and modulates outer segment fluidity and supports phototransduction.^63,64^ Peroxisomal β-oxidation is required to generate DHA through the retroconversion of C24:6ω3 (THA) to DHA, a pathway called Sprecher’s shunt.^65–67^ DHA can be further retroconverted to EPA, a process that also depends on intact peroxisomes,^68^ and an analogous retroconversion pathway exists for ω6 PUFAs.^69^ The physiological relevance of these pathways, however, is poorly understood. Boeck and colleagues reported substantially increased C22:5ω6 in the retinas of *Acox1^-/-^*mice that was attributed to DHA deficiency, but the precise mechanism was difficult to ascertain.^70^ Following a challenge with bovine POS with a defined fatty acid profile consistent with the published lipid composition of rod outer segments,^63,71^ we identified an increase in ω3 and ω6 retroconversion products EPA (C20:5ω3) and AA (C20:4ω6), respectively, in wildtype iRPE but not in *PEX1^-/-^*and *PEX6^-/-^* iRPE (Figure 6). The accumulation of EPA and AA in wildtype iRPE occurred despite these fatty acids not being substantially abundant in POS. Moreover, wildtype iRPE, but not *PEX1^-/-^* and *PEX6^-/-^* iRPE, accumulated EPA and AA when cultured with 30 μM DHA and 30 μM DPAω6 (C22:5ω6), respectively, indicating that the source of EPA and AA is likely from peroxisome-dependent retroconversion (Supplementary Figure 3). Consistent with *Acox1^-/-^*murine retinas, we identified a significant increase in DPAω6, the second most abundant LC-PUFA in POS, in *PEX1^-/-^* and *PEX6^-/-^* iRPE following a POS challenge (Figure 6). We conclude that impaired peroxisome-dependent retroconversion pathways are responsible for DPAω6 accumulation, rather than a deficiency of DHA *per se*. Precisely why the RPE prioritizes retroconverting DHA to EPA (and the ω6 PUFA equivalents) is unknown, but one explanation may be for the generation of retinal VLC-PUFAs, as EPA is the preferred substrate of ELOVL4, an elongase in photoreceptors.^69^ Finally, while the *PEX1^-/-^* and *PEX6^-/-^* iRPE had reduced DHA levels at baseline, following a POS challenge, wildtype, *PEX1^-/-^*, and *PEX6^-/-^* iRPE all accumulated DHA to a similar level (Figure 6), suggesting that RPE may regulate or buffer specifically intracellular DHA levels, but further experiments are required to evaluate this possibility.

Our study has several limitations. First, the pigmentation of the iRPE limited our ability to acquire confocal microscopy images of our cells in a completely unbiased manner (i.e. capturing random fields of view). For example, for determining peroxisome abundance, we imaged iRPE where we could visualize PMP70-positive puncta, which was biased towards less pigmented cells that could differ physiologically from more pigmented iRPE. Second, we lacked a robust means of standardizing and comparing POS uptake precisely across our iRPE as the outer segment fragments differed in size and thus lipid and protein amounts. Lastly, all of our iPSC lines had a chromosome 20q copy number gain (Supplementary Figure 4), a common iPSC karyotype abnormality that may confer a growth advantage in culture.^72^

In summary, our work has established the first human iRPE models to study the effects of peroxisome dysfunction in a disease-relevant cell type. *PEX1^-/-^* and *PEX6^-/-^* iRPE develop normally, but demonstrate significant abnormalities following POS exposure, including impaired DHA retroconversion, reduced outer segment phagocytosis capacity, lipid droplet accumulation, and diminished TEER. Our findings suggest that peroxisomes in the RPE have a critical role in the metabolism of photoreceptor lipids, and disruption of this process may underlie retinal degeneration in patients with peroxisomal disorders.

## Methods

### Sex as a biological variable

Our study utilized iRPE that were all derived from isogenic control iPSCs from a biological female (XX). Sex-specific phenotypic differences have not been described in patients with PBDs. As a result, we expect our results to be generalizable to biological males and females.

### Cell culture

All cells were incubated at 37 °C and 5% CO2 in a Heracell VIOS 160i incubator (Thermo Scientific, Cat. No. 51033546). Matrix-coated culture dishes were prepared using Matrigel (Corning, Cat. No. 354234) that was diluted in DMEM/F12 (Thermo Scientific, Cat. No. 11330032) to yield 10μg/cm^2^ or 100μg/mL according to manufacturer protocols.

A normal human iPSC line (Gibco^TM^ episomal hiPSC, Thermo Scientific, Cat. No. A18945) was used to derive single isogenic *PEX1^-/-^* and *PEX6^-/-^* iPSC clones (generated by Synthego, USA using CRISPR/Cas9-mediated genome editing). All lines were cultured in TeSR™-E8™ media (StemCell Technologies, Cat. No. 05990, Canada), and passaged using ReLeSR™ (StemCell Technologies, Cat. No. 100-0484, Canada).

iRPE were maintained in RPE Maturation Media (RPE-MM) composed of Minimal Essential Medium α (Thermo Scientific, Cat. No. 41061029) supplemented with 2% fetal bovine serum (Millipore Sigma, Cat. No. F1051), 2% KnockOut serum replacement (Thermo Scientific, Cat. No. 10828028), 1x non-essential amino acids (Cytiva, Cat. No. SH3023801 -100x), 1x penicillin/streptomycin (Cytiva, Cat. No. SV30010 -100x), 0.5x N-2 supplement (Thermo Scientific, Cat. No. 175020DPBS48 -100x), 250 µg/mL taurine (Sigma Aldrich, Cat. No. T8691-25G), 14 pg/mL triiodo-L-thyronine (Sigma Aldrich, Cat. No. T5516), and 20 ng/mL hydrocortisone (Sigma Aldrich, Cat. No. H0888) dissolved in Dulbecco’s Phosphate Buffered Saline (DPBS) (Cytvia, Cat. No. 12 SH30028.02). The formulation was derived from a published protocol.^73^ iRPE were passaged using TrypLE™ (Thermo Scientific, Cat. No. 12563011).

### iPSC-RPE differentiation

iPSC-RPE differentiation media recipe was derived from a published protocol^74^ and consisted of HyClone Dulbecco’s Modified Eagle Medium (DMEM) with high glucose (Cytiva, Cat.

No. SH30243.01) supplemented with 1x non-essential amino acids (Cytiva, Cat. No. SH3023801 -100x) and 20% KnockOut serum (Thermo Scientific, Cat. No. 10828028). The media was filtered through a 0.2 μm pore size vacuum filter (Thermo Scientific, Cat. No. 568-0020). The media was changed 3 times per week. At each media change, 50 μM of 2-mercaptoethanol (BioRad, Cat. No. 1610170) was added.

iPSC cultures were harvested using TrypLE™ (Thermo Scientific, Cat. No. 12563011), and single-cell suspension of iPSCs were seeded at 40,000 cells/cm^2^ with TeSR™-E8™ medium supplemented with 2.5 µM blebbistatin (SigmaAldrich, Cat. No. B0560). The following day, TeSR™-E8™ medium without blebbistatin was used until the cells reached 70% confluency. Differentiation then commenced (Figure 1, Day 0) by incubating the cells with differentiation media as outlined below:

For the first week, 10 mM nicotinamide (StemCell Technologies, Cat. No. 07154) was added to the differentiation medium between Day 0 and Day 7. For the second week, 100 ng/mL Activin-A (StemCell Technologies, Cat. No. 78001) was added to the differentiation medium between Day 7 and Day 14. For the following 4 weeks, 3 μM CHIR99021 (StemCell Technologies, Cat. No. 72054) was added to the differentiation medium between Day 14 and Day 42, after which the cells reached a committed RPE stage. At Day 43, cells were harvested using TrypLE™ (Thermo Scientific, Cat. No. 12563011), and either pooled for cryopreservation or RPE maturation proceeded by generating a single-cell suspension, seeding 300,000 cells/cm^2^, and maintaining the cells in RPE-MM for 3 additional weeks. At Day 63, iRPE cells were similarly harvested and seeded at 300,000 cells/cm^2^ while supplementing the media with 10 μM Y-27632 (StemCell Technologies, Cat. No. 72304, Canada) during seeding. The cells were considered mature RPE three weeks after their final seeding. iRPE from independent wells were used for experimental replicates.

### Validating knockout of PEX1 and PEX6 in iRPE

Sanger sequencing performed for *PEX1^-/-^* iRPE confirmed a homozygous variant c.178dup; p.(W60Lfs*8) in *PEX1* (NM_000466.3) in exon 2 of 24 (Supplementary Figure 1A). Sanger sequencing performed for *PEX6^-/-^* iRPE confirmed a homozygous variant c.197dup; p.(Q67Afs*11) in *PEX6* (NM_000287.4) in exon 1 of 17 (Supplementary Figure 1B).

### iPSC transfection

iPSC colonies were cultured on Matrigel-coated chamber slides (Sarstedt, Cat. No. 946170802). Each well was transfected with 300 ng of GFP-SKL (Addgene, plasmid #53450) or pmaxGFP® (Lonza) using Lipofectamine 2000 (Invitrogen, Cat. No. 11668019) for 48 hours. The cells were fixed with 4% paraformaldehyde (Sigma Aldrich, Cat. No. P6148-500G), incubated with 1.67 μM DAPI (Millipore Sigma, Cat. No. D8417) for 5 minutes at room temperature, and mounted with ProLong™ Glass Antifade Mountant (Thermofisher, Cat. No. P36984) following DPBS washes between steps.

### Western blotting

Cells were harvested mechanically with cell scrapers in radioimmunoprecipitation (RIPA) buffer (Thermo Scientific, Cat. No. 89900) supplemented with cOmplete™ Protease Inhibitor Cocktail (Sigma Aldrich, Cat. No. 11697498001).

Protein concentration was determined using Pierce™ BCA Protein Assay Kit (Thermo Scientific, Cat. No. 23225). Multiskan™ GO Microplate Spectrophotometer (Thermo Scientific, Cat. No. 51119300) was used to measure absorbance at 562 nm.

Samples were mixed with 1x Laemmli Sample Buffer (Bio-Rad, Cat. No. 1610747 – 4x) supplemented with 10% 2-mercaptoethanol (Sigma Aldrich, Cat. No. M3148) and incubated at 70 °C for 10 minutes. Samples were then loaded onto 4-15% Mini-PROTEAN® TGX™ Precast Protein Gels precast (Bio-Rad, Cat. No. 4561084) housed in Mini-PROTEAN® Tetra Cell (Bio-Rad, Cat. No. 1658037) submerged with 1 L of running buffer (25 mM Tris, 190 mM glycine, 0.1% SDS). Following electrophoresis at 100 V for 120 minutes, proteins were transferred to a nitrocellulose membrane (Thermo Scientific, Cat. No. IB23002) at 23 V for 6 minutes with the iBlot™ 2 dry blotting system (Thermo Scientific, Cat. No. IB21001).

Membranes were protected from light and incubated with Intercept® PBS Blocking Buffer (Li-Cor, Cat. No. 927-70001) for 1 hour at room temperature. All incubations and washes were performed on a rocker. A 1:1 blocking buffer:PBS-T (PBS containing Tween-20 0.05%) solution was prepared to dilute antibodies for incubations. Primary antibodies (Supplementary Table 1) were applied overnight at 4 °C. After the incubation, membranes were washed with PBS-T three times for 5 minutes each. Secondary antibodies (Supplementary Table 1) were applied for 1 hour at room temperature. Following the incubation, membranes were washed with PBS-T twice for 10 minutes each and then once with PBS for 10 minutes.

Membranes were scanned using the Odyssey CLx imaging system (Li-Cor) and visualized on Image Studio (LiCor, V.5.2). Blots were captured with a high-quality setting at 169 μm resolution. Regions of interest were manually selected and densitometry values were recorded. Values were adjusted using their respective loading controls and normalized to wildtype using Microsoft® Excel® (Version 2401 Build 16.0.17231.20236)

### Immunofluorescence microscopy

Cells were grown on Matrigel-coated chamber slides (Ibidi, Cat. No.81817) and were washed once with DPBS and then incubated on ice with 4% paraformaldehyde (Sigma Aldrich, Cat. No. P6148-500G) for 15 minutes. Cells were then washed 3 times with DPBS and blocked and permeabilized by incubation in PBS supplemented with 0.1% saponin (Millipore Sigma, Cat. No. 558255) and 1% bovine serum albumin (Sigma Aldrich, Cat. No. A9418) for 30 minutes at room temperature. Incubation with primary antibodies occurred for 1 hour at room temperature in blocking buffer. Cells were then washed 3 times with DPBS and incubated with secondary antibodies at room temperature for 1 hour in blocking buffer. Cells were again washed 3 times with DPBS and then incubated with 1.67 μM DAPI (Millipore Sigma, Cat. No. D8417) or 6.67 μM Hoechst (Thermo Scientific, Cat. No. 33342) for 5 minutes at room temperature. Cells were finally washed 3 times with DPBS, and the slide was mounted with ProLong™ Glass Antifade Mountant (Thermofisher, Cat. No. P36984). The slides were left to dry at room temperature overnight, protected from light. Alternatively, cells were kept in DPBS and imaged.

### Lipidomics

iRPE cultured on Matrigel-coated 60 mm dishes were harvested using TrypLE™ (Thermo Scientific, Cat. No. 12563011). After creating a single-cell suspension, samples were mixed 1:1 with Trypan Blue (HiMedia, Cat. No. TCL005-100ml) and counted using the Countess 3 (Thermo Scientific, Cat. No. AMQAX2000). Cells were then centrifuged at 400 g for 5 minutes, and the resulting pellets were resuspended in 100 μL of molecular grade water. Gas and liquid chromatography-mass spectrometry were performed by the Peroxisomal Diseases Laboratory at the Kennedy Krieger Institute in Baltimore, MD, USA. Lipid fatty acids were acid hydrolyzed from triglycerides and phospholipids then measured as their pentafluorobenzyl bromide esters by isotope dilution capillary gas chromatography negative chemical ion GC-MS according to the method of Lagerstedt et al 2001^75^. Acylcarnitines and phosphatidylethanolamine plasmalogens were methanol extracted and analyzed by LC-MS/MS on an AB Sciex API3200 with either a Phenomenex Kinetex C8 column (2.6μm x 50 x 2.1 mm) using a linear gradient between 0.1% formic acid in H2O and 0.1% formic acid in methanol in positive electrospray ionization mode according to the method van de Beek 2016^76^ or a Phenomenex Kinetex HILIC column (2.6μm x 30 x 3 mm) column using a linear gradient between 0.2% formic acid in 60mM ammonium formate and 0.2% formic acid in acetonitrile according to modified methods of De Biase et al., 2023^77^ and Scherer et al., 2010^78^.

### Bovine photoreceptor outer segments

Isolated bovine photoreceptor outer segments (POS) were acquired from InVision BioResources (22/2/2024 lot). POS pellets were gently resuspended in 10 mL of 8.3 pH wash buffer containing 20 g of sucrose (Fisher Scientific, Cat. No. BP2201) and 1.68 g of sodium bicarbonate (Fisher Scientific, Cat. No. BP3281), dissolved in a total volume of 200 mL

Milli-Q water (Millipore Sigma). The resuspended POS were centrifuged at 600 g for 20 minutes at 4 °C. The resulting pellet was resuspended in 10 mL of wash buffer. Aliquots were mixed 1:1 with RIPA buffer (Thermo Scientific, Cat. No. 89900) supplemented with cOmplete™ Protease Inhibitor Cocktail (Sigma Aldrich, Cat. No. 11697498001) for determining the protein concentration using Pierce™ BCA Protein Assay Kit (Thermo Scientific, Cat. No. 23225). For fluorescent probe conjugation, 1 mg of pHrodo™ Red (Thermo Scientific, Cat. No. P36600), a pH-sensitive dye that fluoresces more intensely in acidic environments, was dissolved in 150 uL of dimethyl sulfoxide (Thermo Scientific, Cat. No.D12345) and combined with 10 mg of POS suspended in the wash buffer for 1 hour at room temperature. After the conjugation reactions, the labeled POS were washed three times with 5 mL of the wash buffer and centrifuged at 600 g for 20 minutes at 4 °C between steps. For use in immunoblotting experiments, approximately 0.5 POS per cell was delivered in RPE-MM supplemented with 4 μg/mL human protein S (Sino Biologicals, Cat. No. 12179-H08H-100) and 1.5 μg/mL human MFG-E8 (Sino Biologicals, Cat. No. 10853-H08B-1).

### Flow cytometry

Suitable gating strategies and other technical aspects of flow cytometry were developed in consultation with the University of Alberta Flow Cytometry Core Facility. A minimum of 10,000 events were captured for each sample on the flow cytometer. Cells were gated based on the FSC/SSC (forward scatter/ side scatter) profile to exclude debris and cell clumps. FSC area versus FSC height was then used to remove doublets. Single cells were gated based on fluorescence minus one (FMO) controls. Data including median fluorescence intensities were analyzed using FlowJo v10.10.0 (BD Biosciences).

*Assessing Expression of Signature RPE Proteins:* iRPE were harvested using TrypLE™ then fixed with 4% paraformaldehyde. The cells were washed with flow cytometry buffer (DPBS supplemented with 2% fetal bovine serum (Sigma Aldrich, Cat. No. F1051-500ml)) between steps. To permeabilize cells, 0.2% Triton X-100 (Bio-Rad, Cat. No. 1610407) was added to the flow cytometry buffer. Antibodies were incubated overnight at 4 °C on a rocker. The next day, cells were washed and incubated in secondary antibodies with the permeabilization buffer for 30 minutes at room temperature. The cells were then incubated in the flow cytometry buffer containing 1.67 μM DAPI (Millipore Sigma, Cat. No. D8417) for 5 minutes at room temperature. After washes, the cells were finally resuspended in the flow cytometry buffer. Ten thousand events per sample were acquired on a 5-laser Cytek Aurora spectral flow cytometer (RRID:SCR_019826, Cytek Biosciences, Fremont, CA, USA) running SpectroFlo 3.1.0 with following panels: DAPI, MITF Alexa Fluor 488, BEST1 Alexa Fluor 546, and PMEL17 Alexa Fluor 647; or DAPI, PAX6 Alexa Fluor 488 and TYRP1 Alexa Fluor 647. Threshold was on FSC (50,000) in both cases.

*Measuring Intracellular Neutral Lipids and Phagocytosis with pHrodo-Labeled POS:* iRPE were harvested using trypsin (Thermo Scientific, Cat. No. 25200072) at baseline and following a POS challenge consisting of the addition of 3 POS/cell daily for 7 days. Live cells were resuspended in DPBS and incubated with 2 μM Calcein (Thermo Scientific, Cat. No. L32250) for 30 minutes at room temperature. LipidTOX Deep Red neutral lipid stain (Fisher Scientific, Cat. No. H34477) (1:100 dilution in DPBS) mean fluorescent intensity (MFI) was used to measure intracellular neutral lipid accumulation. Internalized pHrodo Red labeled POS MFI was used to measure POS uptake and trafficking efficiency to intracellular acidic compartments (Supplementary Figures 5 and 6). Cells were treated with 1 μM bafilomycin (Abcam, Cat. No. 120497), a specific inhibitor of V-ATPase that prevents lysosomal acidification, for three hours to facilitate gating of the presumed lysosomal pHrodo fraction. The frequency of live cells that were positive for any pHrodo signal was used to determine uptake. At least 100,000 total events were acquired using an Attune NxT Acoustic Focusing Cytometer (RRID:SCR_019590, ThermoFisher Scientific, Burlington, ON, Canada) equipped with 4 lasers employing the indicated bandpass filters: 488 nm BL1 530/30 for Calcein, 561 nm YL1585/16 for pHrodo, and 634 nm RL1 670/14 for LipidTOX Deep Red. Compensation was applied among fluorochromes. Threshold was on FSC (25,000). Voltages used were FSC-A 70 V, SSC-A 280 V, YL1-A 250 V, RL1-A 220 V, and VL1-A 220 V.

### Transepithelial electrical resistance

The EVOM3 (World Precision Instruments) was used to measure the transepithelial electrical resistance (TEER) of iRPE cultured on Matrigel-coated transwells (Corning, Cat. No. 3470) at baseline and following 150 μM palmitate treatment or a 24-hour POS (10 POS/cell) challenge.

### Statistical Analysis

To quantify PMP70 positive puncta, captured z-stacks were analysed using IMARIS (Oxford Instruments, V. 10.1). The spots detection tool was employed with a 0.3 μm diameter parameter to automatically detect PMP70 positive puncta. Cell borders were manually drawn using the draw tool. The filter tool was used to determine the number of spots inside each cell by selecting all spots that were less than 0 μm away from the cell’s three-dimensional surface. Data was managed using Microsoft® Excel® (Version 2401 Build 16.0.17231.20236).

Maximum intensity projections were developed using ImageJ (NIH, V1.53t). All statistical analysis and graphing were executed in GraphPad Prism version 10.6.0. Statistical comparisons between wildtype, *PEX1^-/-^* and *PEX6^-/-^* iRPE were performed using one- or two-way ANOVA with Dunnett’s post-hoc test. Illustrations were created using BioRender or Microsoft® PowerPoint® (Version 2512 Build 16.0.19530.20184).

### Study Approval

Ethics approval was obtained for the use of human iPSCs for research (Pro00074451).

## Data Availability

All data reported in this paper will be shared by the lead contact, Matthew D. Benson upon request.

## Author Contributions

C.M. designed and conducted the experiments, analyzed the data, and drafted the original manuscript. C.B.F, S.H., J.H., E.T., Q.Z., A.R., H.H., B.C.H., E.M., and A.R. conducted experiments and analyzed the data. T.D.F., R.B.H., J.M.L.M., and M.D.B. conceptualized experiments, developed methodology, and supervised research activities. M.D.B. reviewed and edited the manuscript and generated the final draft of the manuscript.

## Funding Support

This research is funded by Fighting Blindness Canada (grant #RG2401). Acknowledgement is made to the donors of the Postdoctoral Fellowship Program in Macular Degeneration Research (MDR), grant # M2024001F, a program of the BrightFocus Foundation, for support of this research. J.M.L.M. is supported by the James Grosfield Initiative for Dry AMD, an RPB Career Development Award, and a National Eye Institute Career Development Award (K08EY033420). M.D.B. is supported by a Bayer Professorship in Translational Research in Ophthalmology.

## Acknowledgments

The authors would like to thank all former and current members of the Benson lab. We thank Drs. Andrew Simmonds, Brittany Carr, Ian MacDonald, and Robin Clugston for their technical expertise, experimental advice, and sharing of reagents. We thank Dr. Davide Ortolan for sharing his expertise with REShAPE, and Dr. Mitra Farnoodian for sharing her pHrodo conjugation protocol. Lai Xu and Dr. Gabrielle Siegers assisted with experiments performed at the University of Alberta Flow Cytometry Facility (RRID:SCR_019195, which receives financial support from the Faculty of Medicine & Dentistry (FoMD) and Canada Foundation for Innovation (CFI) awards to contributing investigators) and Cell Imaging Core (RRID:SCR_019200, which receives financial support from the FoMD, the University Hospital Foundation, Striving for Pandemic Preparedness – The Alberta Research Consortium, and CFI awards to contributing investigators).

## List of Abbreviations

AAA: ATPases Associated with Diverse Cellular Activities
ACAA1: Acetyl-Coenzyme A Acyltransferase 1
ACOX1: Acyl-Coenzyme A Oxidase 1 BCFAs Branched-Chain Fatty Acids
DPBS: Dulbecco’s Phosphate Buffered Saline
FAs: Fatty Acids
iPSCs: Induced Pluripotent Stem Cells
iRPE: iPSC-Derived RPE
IRDs: Inherited Retinal Dystrophies
LDs: Lipid Droplets
MFP2: Multifunctional Protein 2
PBDs: Peroxisome Biogenesis Disorders
POS: Photoreceptor Outer Segments
PTS1: Peroxisomal Targeting Signal-1
PTS2: Peroxisomal Targeting Signal-2
PUFA: Polyunsaturated Fatty Acids
RPE: Retinal Pigment Epithelium
RPE-MM: Retinal Pigment Epithelium Maturation Media
VLCFAs: Very Long-Chain Fatty Acids

## Supplementary Tables and Figures

**Supplementary Table 1.**
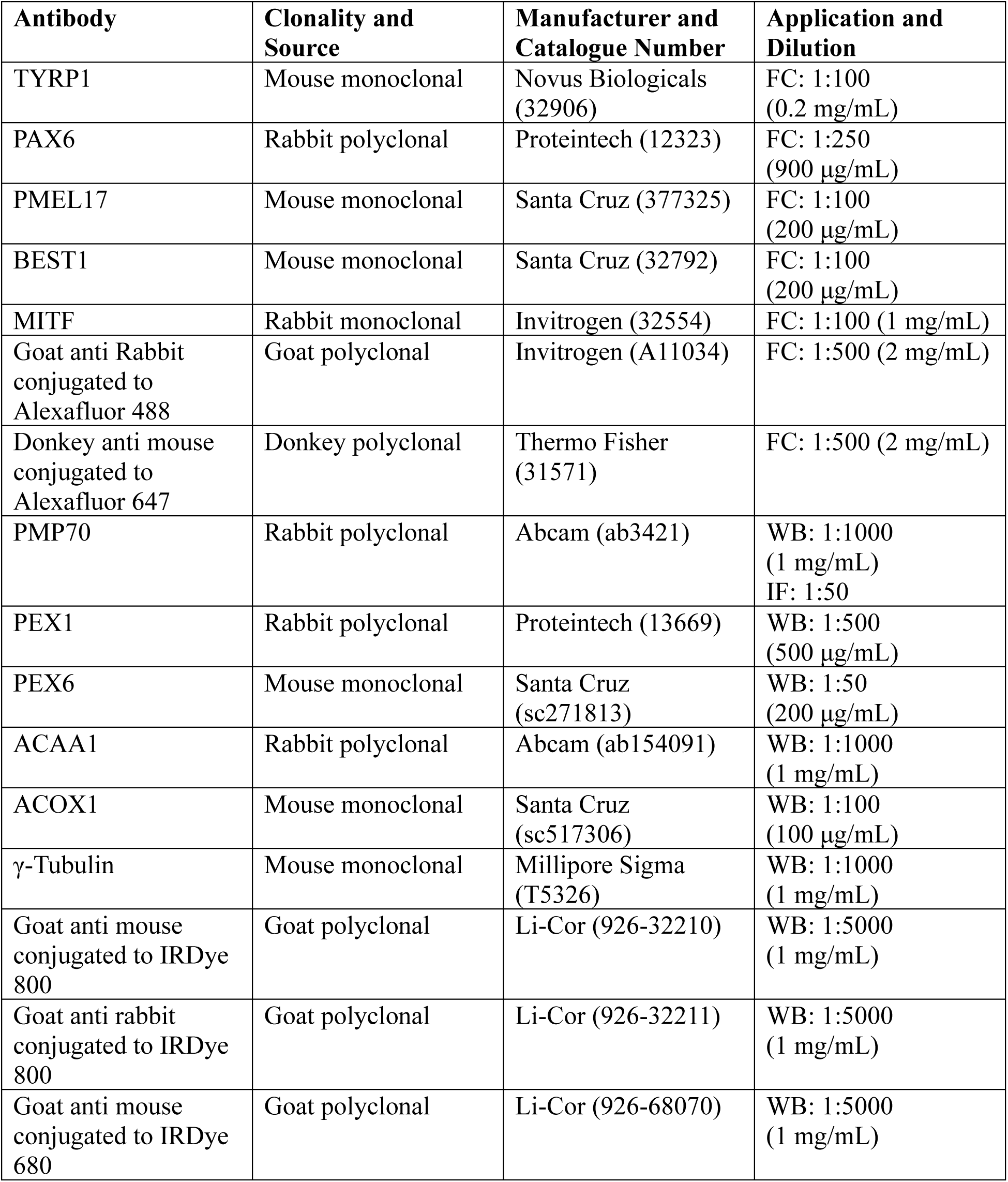
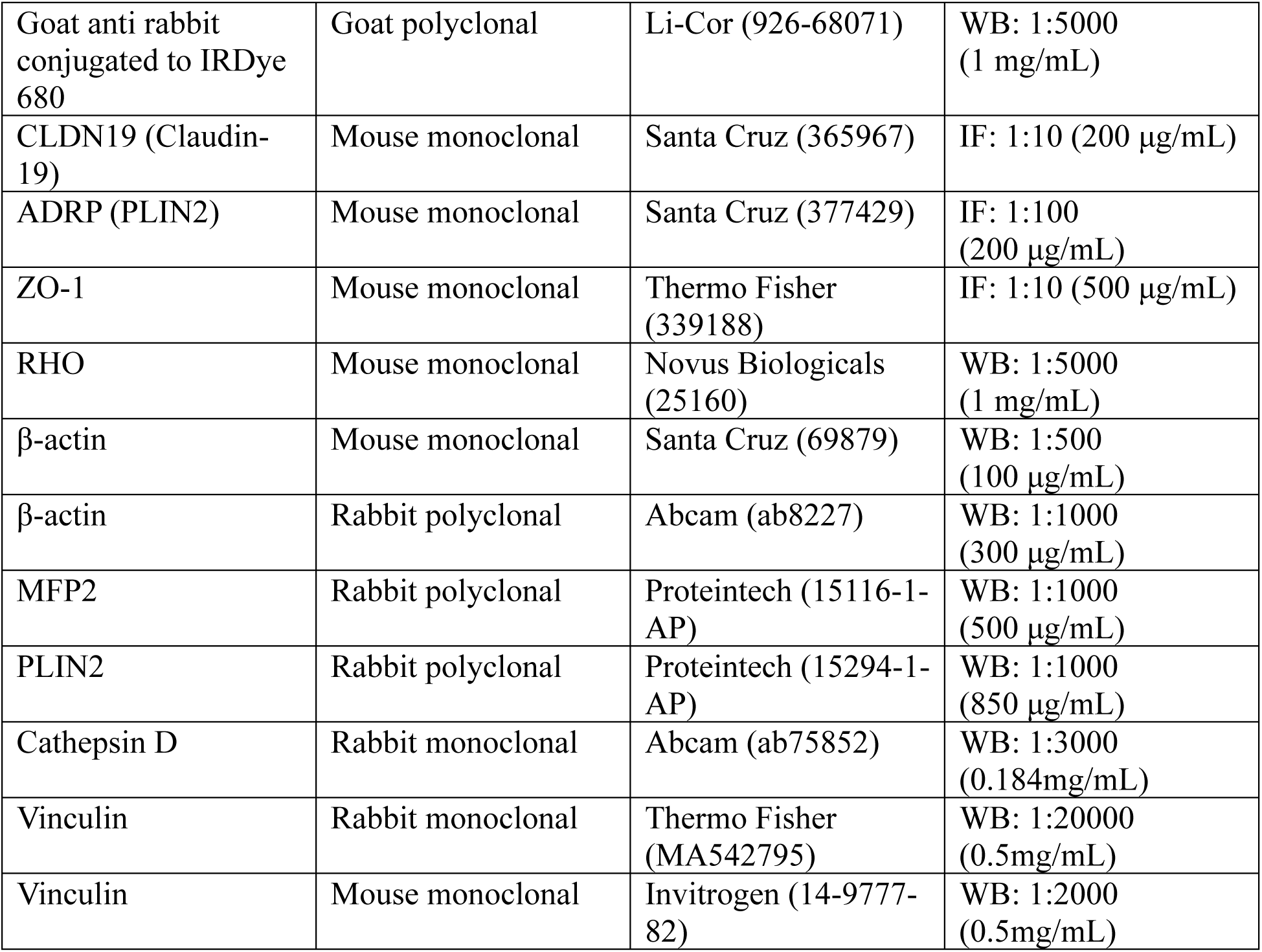
List of antibodies with corresponding application and concentration. FC = flow cytometry; WB = western blot; IF = immunofluorescence.

**Supplementary Table 2.**
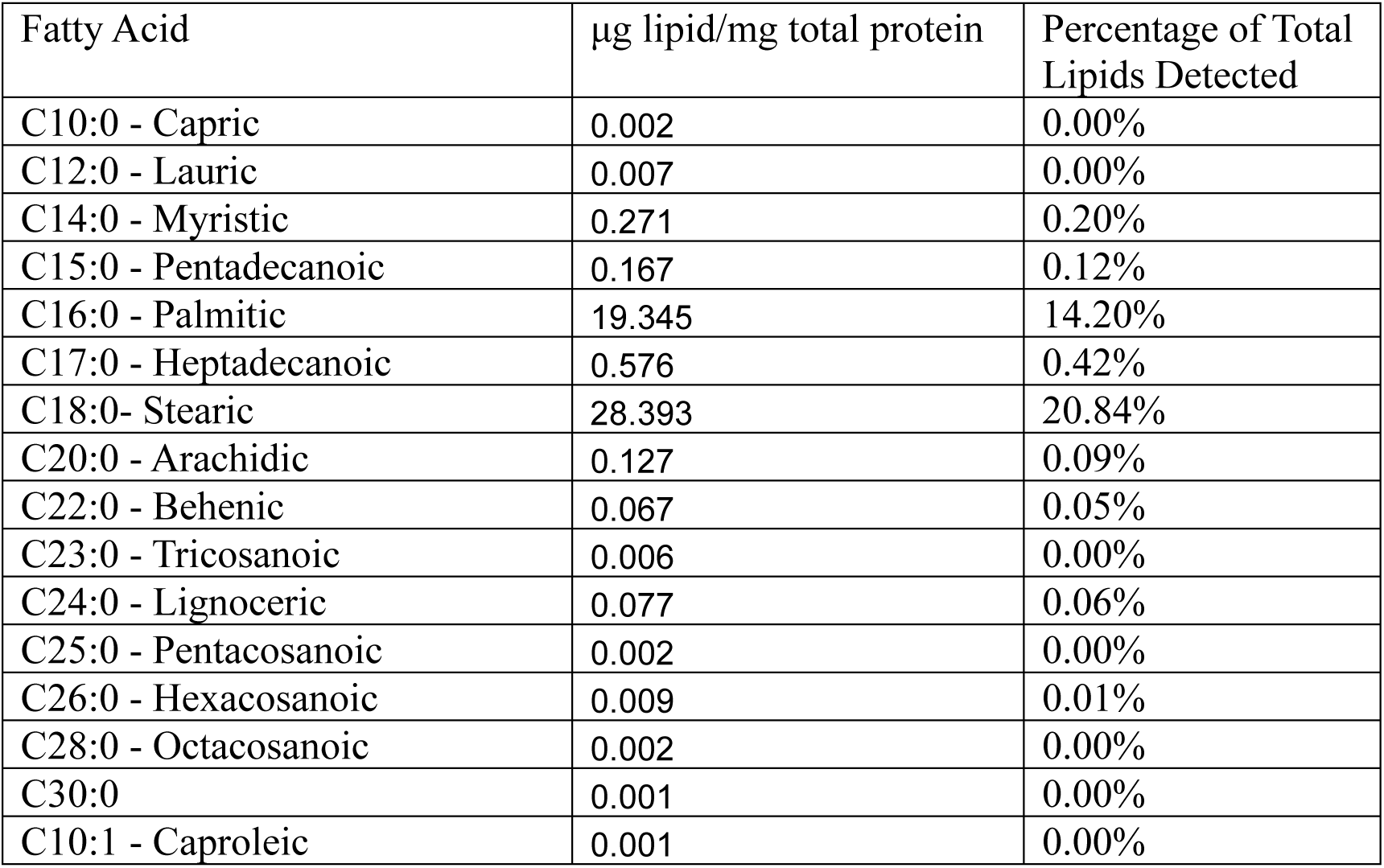
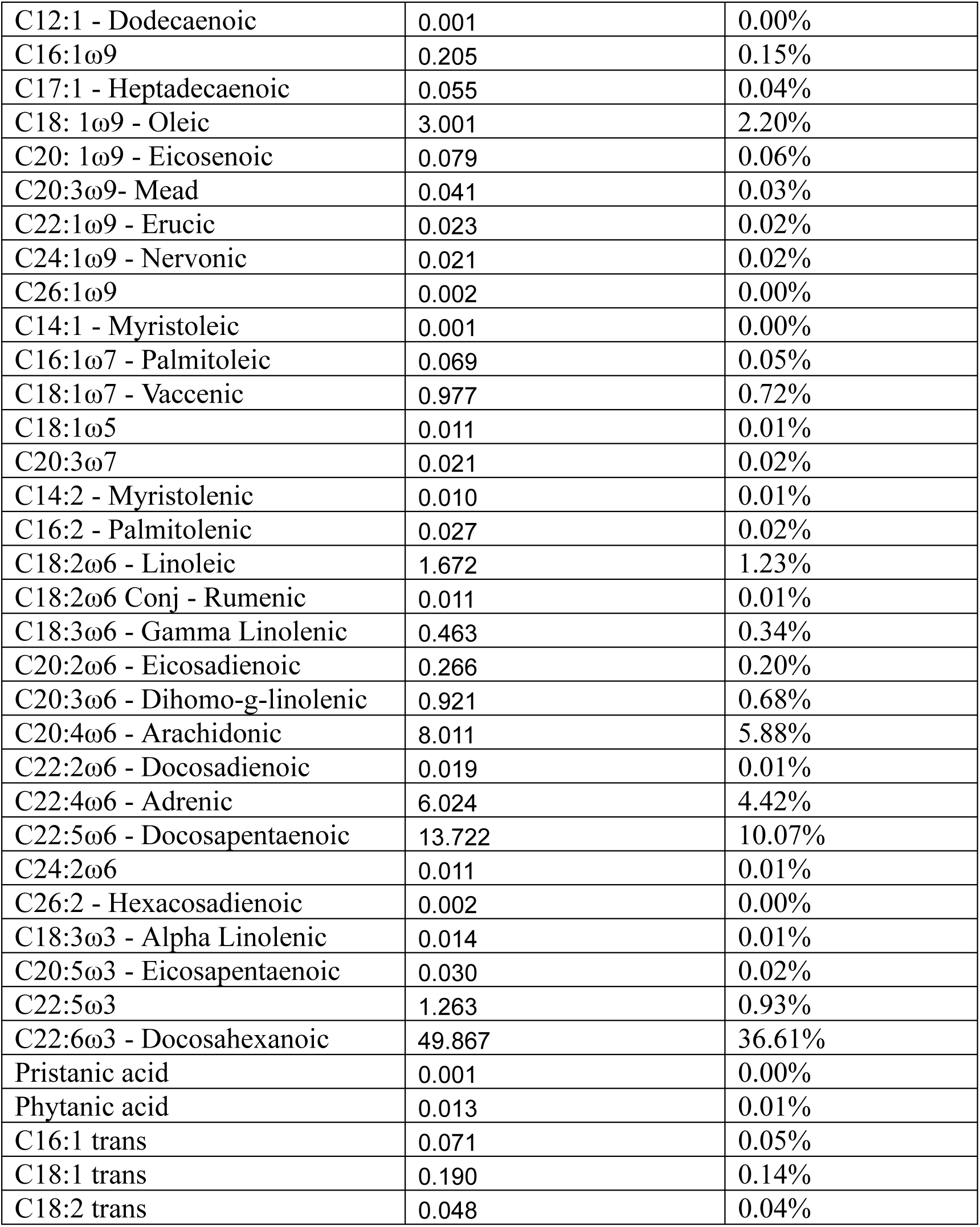
Comprehensive list of total fatty acid species detected in bovine POS using GC-MS. The mean value for each fatty acid species is presented (n=3).

### DNA Sanger Sequencing

DNA was isolated from iRPE using GeneJET Genomic DNA Purification Kit (Thermo Scientific, Cat. No. K0722) and eluted in molecular grade water. DNA concentration was determined using a μDrop™ plate (Thermo Scientific, Cat. No. N12391) and Multiskan™ GO Microplate Spectrophotometer (Thermo Scientific, Cat. No. 51119300).

To confirm the genotype of *PEX1^-/-^* iRPE, 50 μL PCR reactions were performed using Taq DNA polymerase (New England Biolabs, Cat. No. M0273L) with forward (5’-GAACTCTTTTTGGACATGTGAATTG-3’) and reverse (5’- GCAAGTAGGGAGTATGGTAAACT-3’) primers (0.2 μM). Reactions were subject to one denaturation step for 30 seconds at 95 °C, followed by 30 cycles of 15 seconds duration at 95 °C, 15 seconds at 49 °C, and 27 seconds at 68 °C, followed by one extension step for 5 minutes at 68 °C.

To confirm the genotype of *PEX6^-/-^*iRPE, 50 μL touchdown PCR reactions were performed using Phusion DNA polymerase and GC buffer (Thermo Scientific, Cat. No. F534S) with forward (5’-AGAAACCGCAAAGGAGGAC-3’) and reverse (5’-ACTAGTCGTCTGGCTCTCTG-3’) primers (0.2 μM). Reactions were subject to one denaturation step for 30 seconds at 98 °C, followed by 10 touchdown cycles of 10 seconds at 98 °C, 20 seconds at 66 °C (decreasing by 0.5 °C each cycle), and 21 seconds at 72 °C. This was followed by 25 cycles of 30 seconds at 98 °C, 20 seconds at 61 °C, and 21 seconds at 72 °C, followed by one extension step for 5 minutes at 72 °C.

**Supplementary Figure 1:**
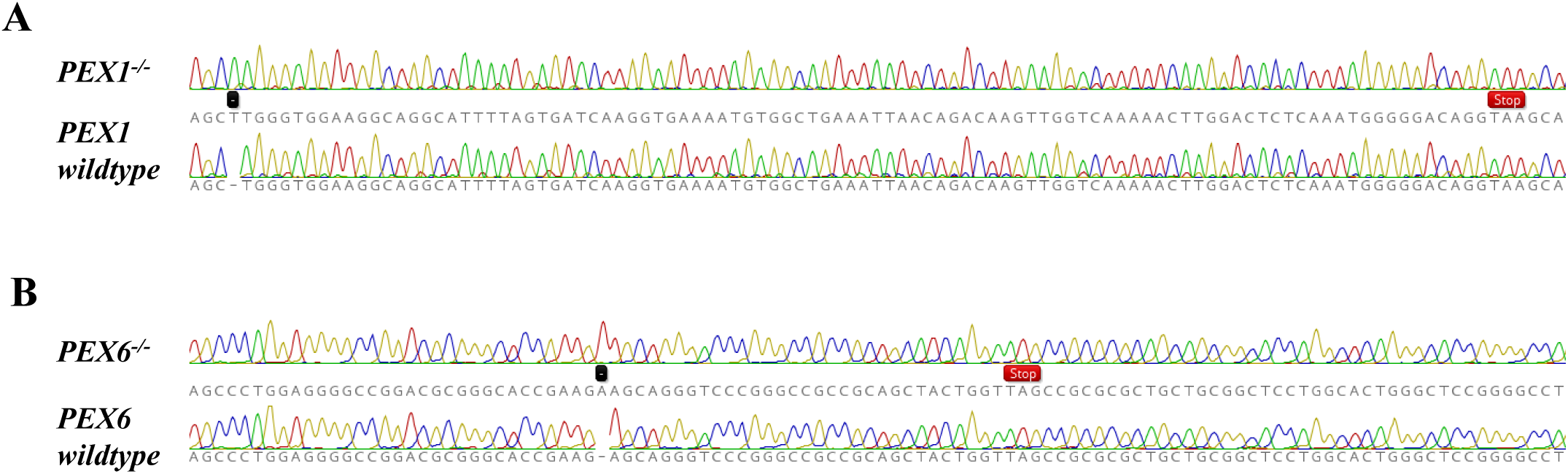
Sanger sequencing of iRPE DNA. Frameshift mutations caused by single base pair duplications (indicated by a black square) resulted in stop codons (indicated by red rectangles).

**Supplementary Figure 2:**
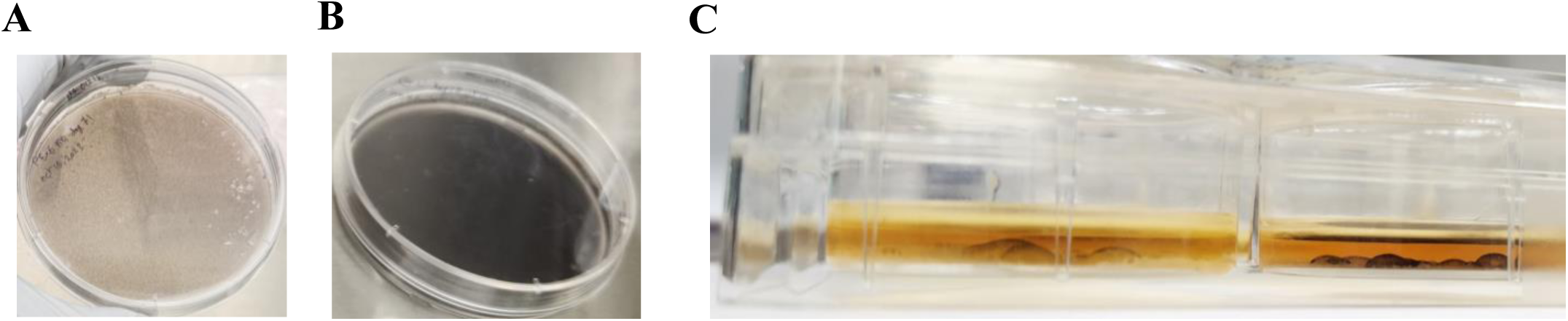
Pigmentation and focal monolayer detachments in iRPE. A. Maturing iRPE cells at day 74 demonstrate mild pigmentation. B. Mature iRPE at day 95 demonstrate significant pigmentation. C. Cross-sectional image of differentiating iRPE cells demonstrate focal monolayer detachments (day 37).

**Supplementary Figure 3:**
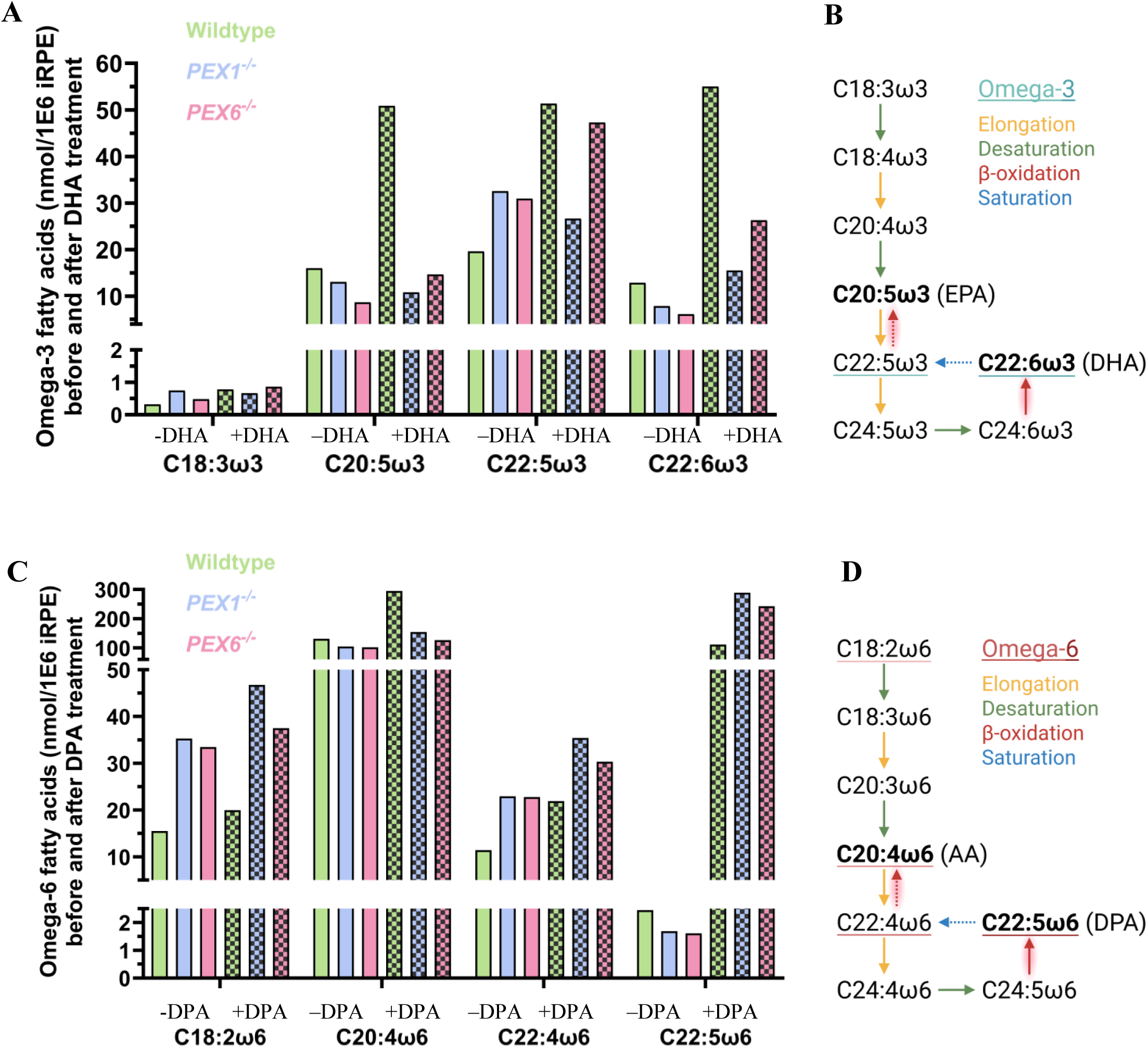
Quantitative analysis of fatty acid profiles shifts in iRPE following DHA and DPA treatments. A. Quantification of omega-3 fatty acids in iRPE lysates by GC-MS before and after a 24-hour treatment with 30 μM DHA (n=1). B. The omega-3 fatty acid metabolic pathway adapted from Yu et al., 2012.^76^ Reactions that are presumed to require intact peroxisome β-oxidation are highlighted in red. C. Quantification of omega-6 fatty acids in iRPE lysates by GC-MS before and after a 24-hour treatment with 30 μM omega-6 DPA (n=1). D. The omega-6 fatty acid metabolic pathway adapted from Yu et al., 2012.^76^ Reactions that are presumed to require intact peroxisome β-oxidation are highlighted in red.

**Supplementary Figure 4:**
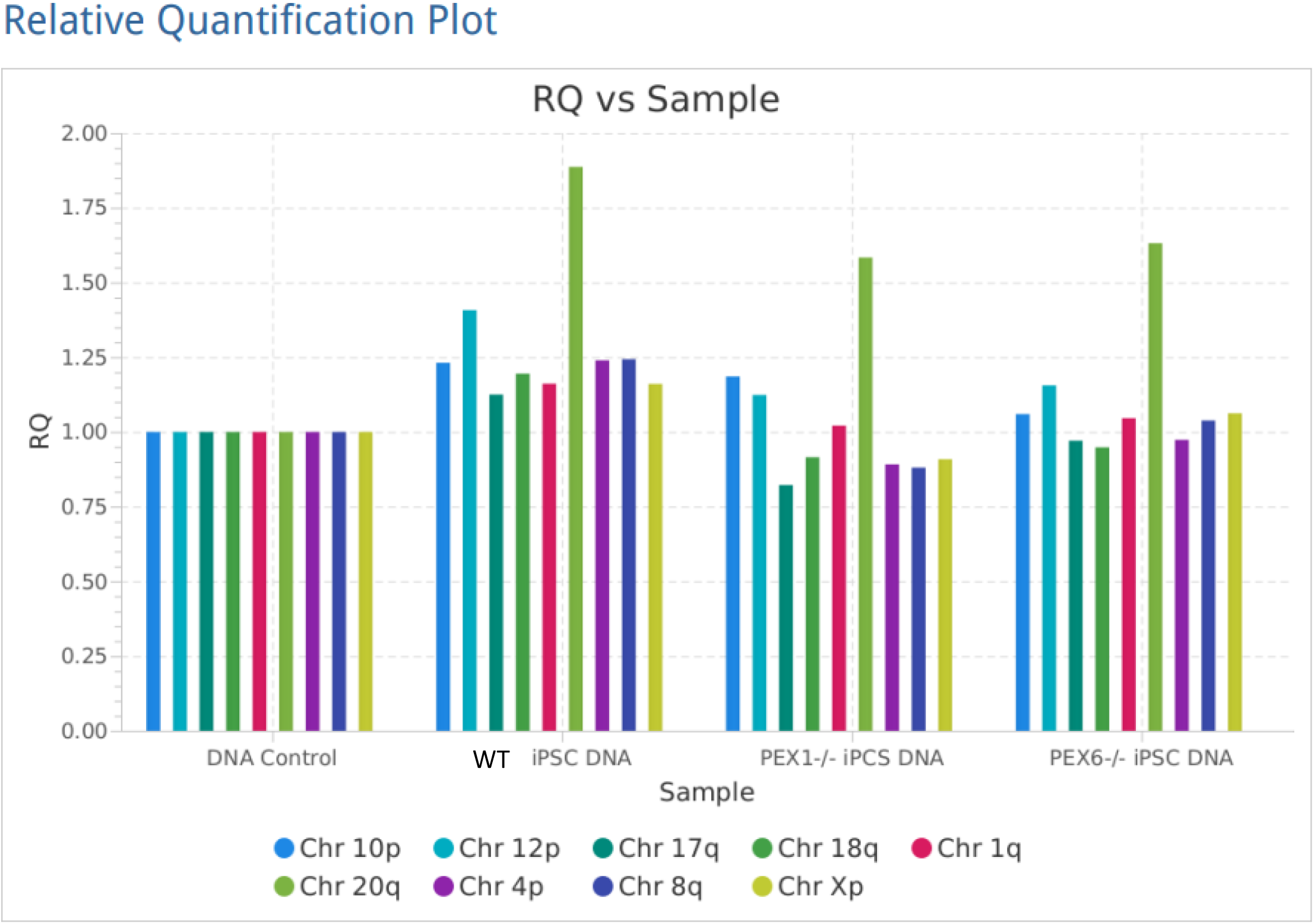
Quantitative PCR in wildtype, *PEX1^-/-^* and *PEX6^-/-^* iPSCs for commonly reported karyotypic abnormalities. Chr20q has an amplification in a minimal critical region in wildtype, *PEX1^-/-^* and *PEX6^-/-^* iPSCs.

**Supplementary Figure 4:**
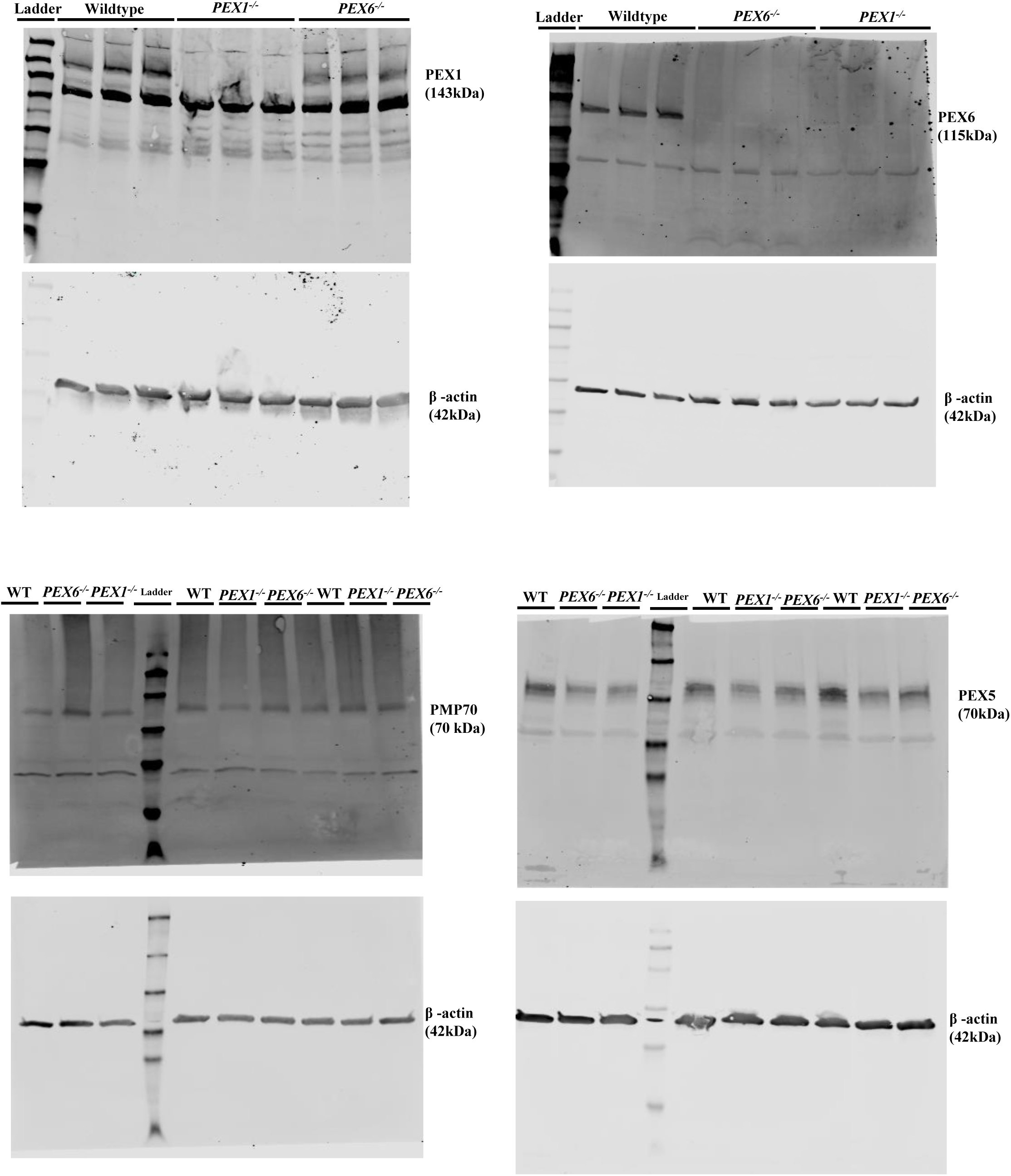
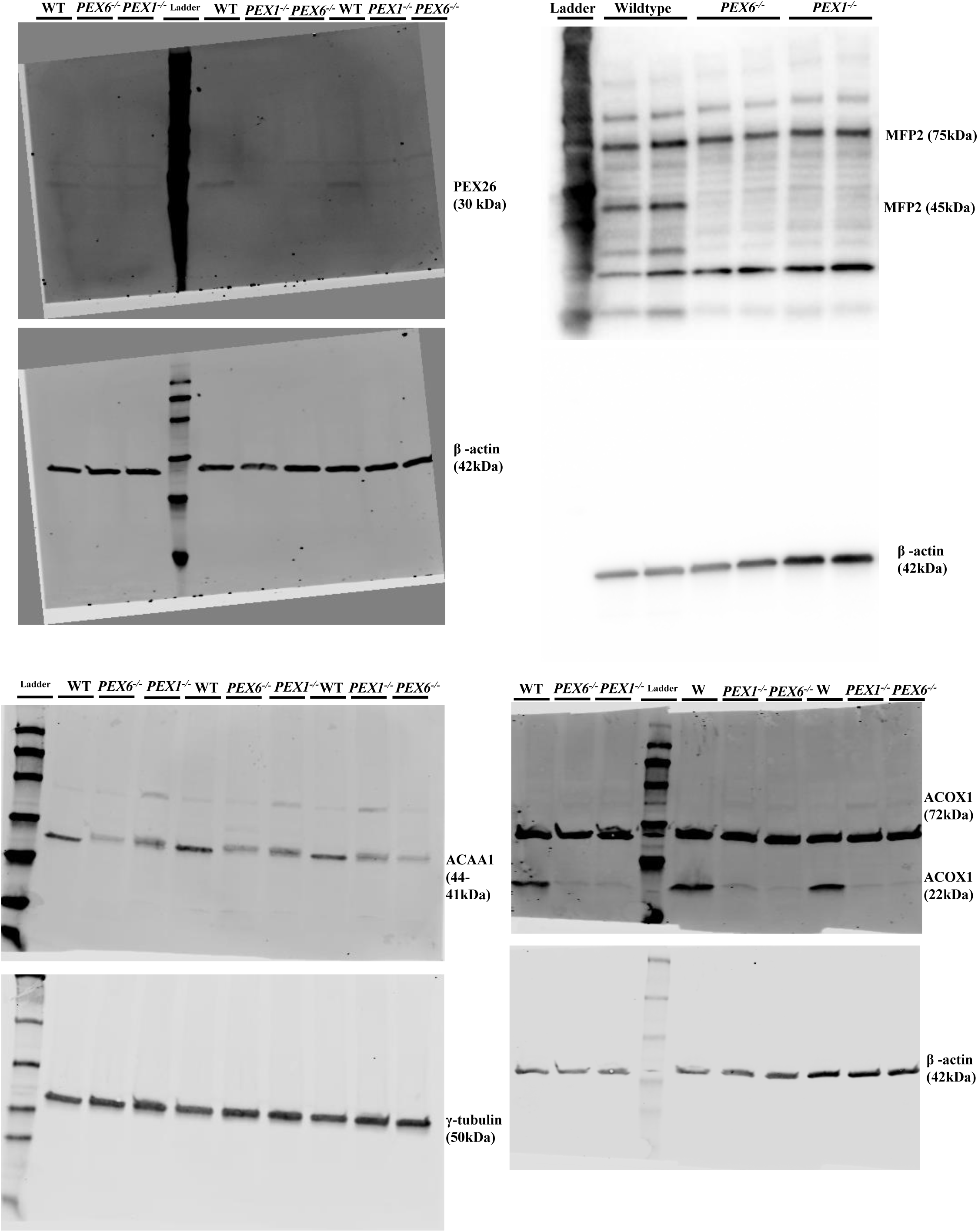
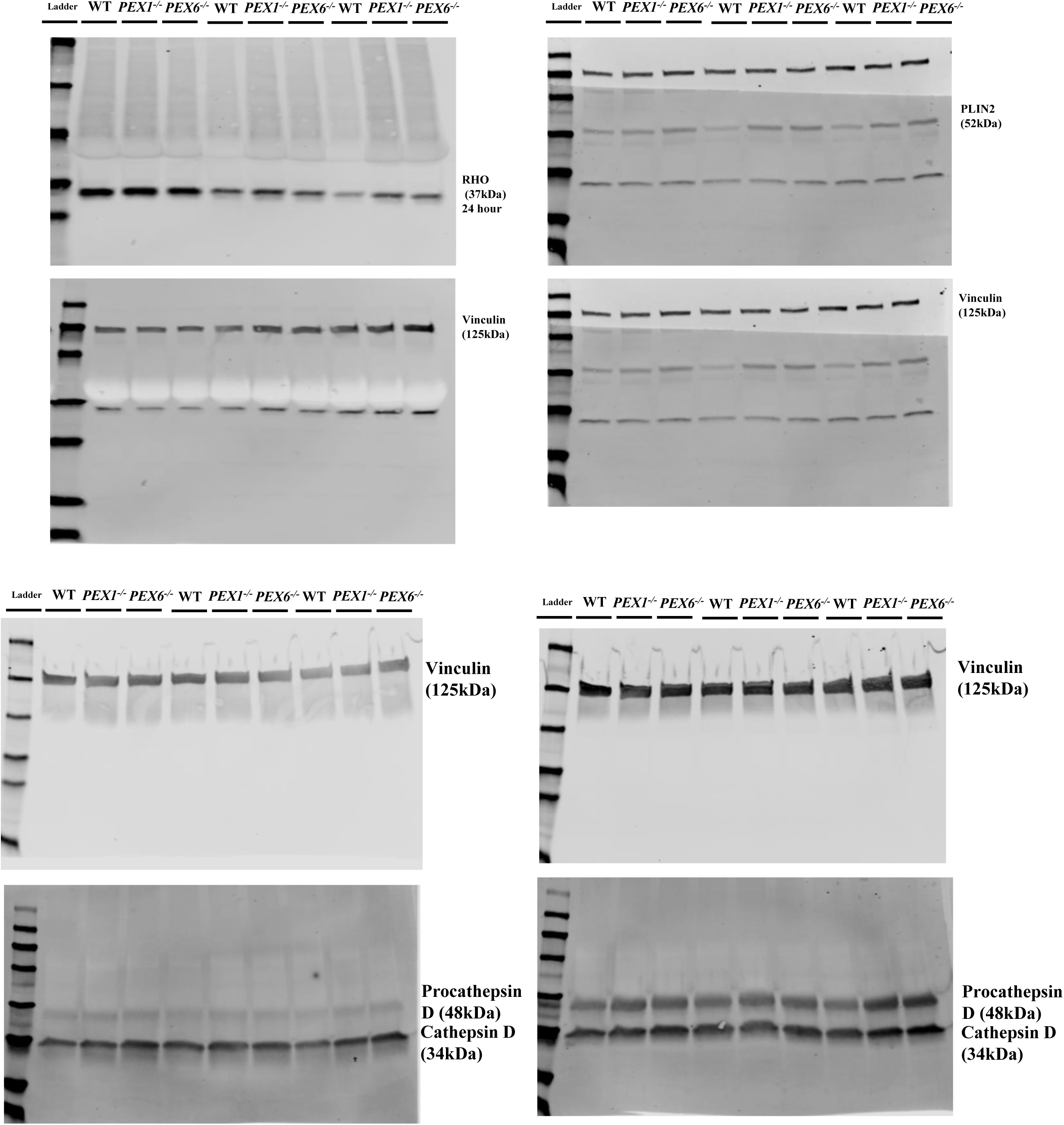
Full membranes of all immunoblot images presented.

**Supplementary Figure 5:**
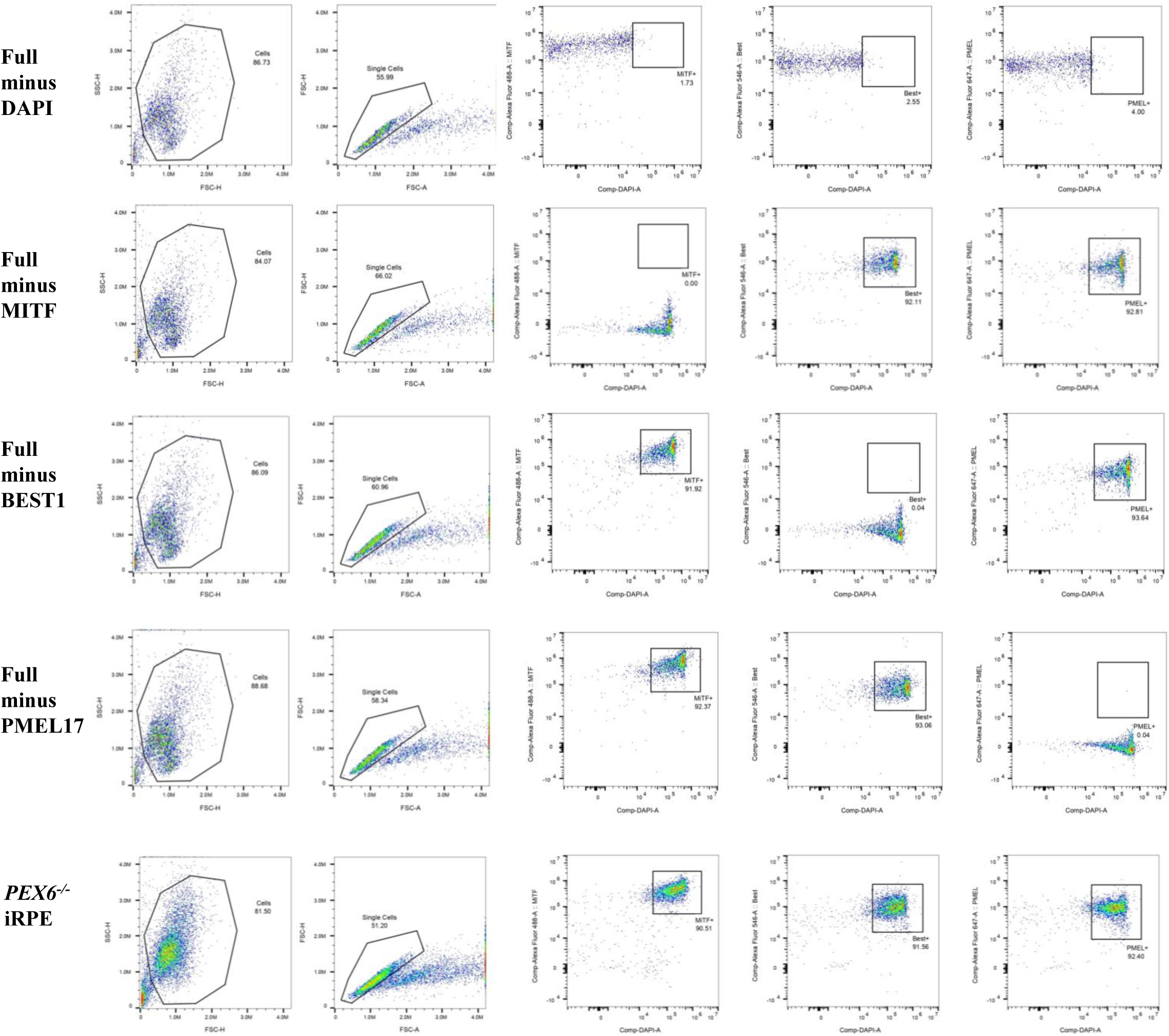
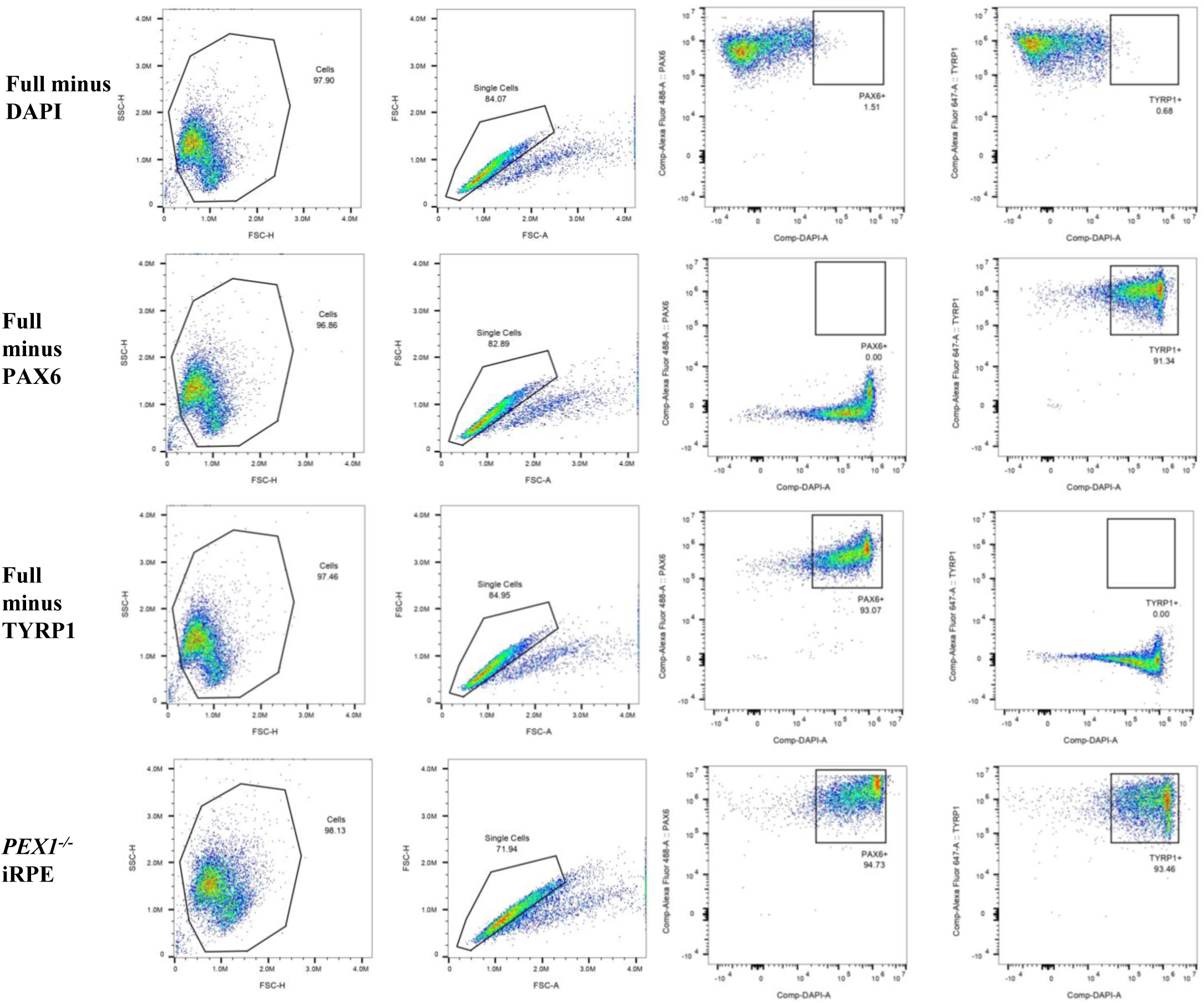
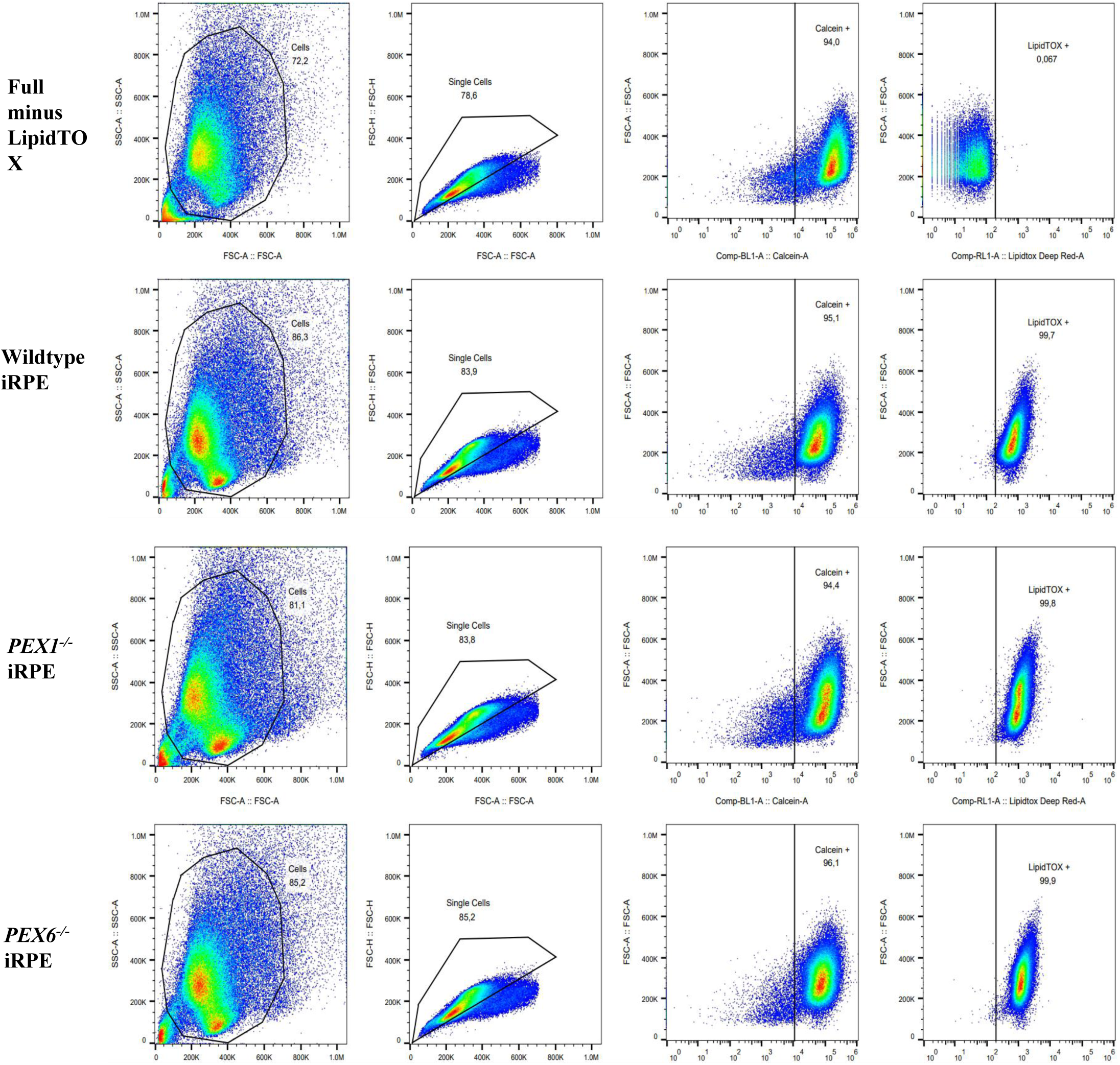
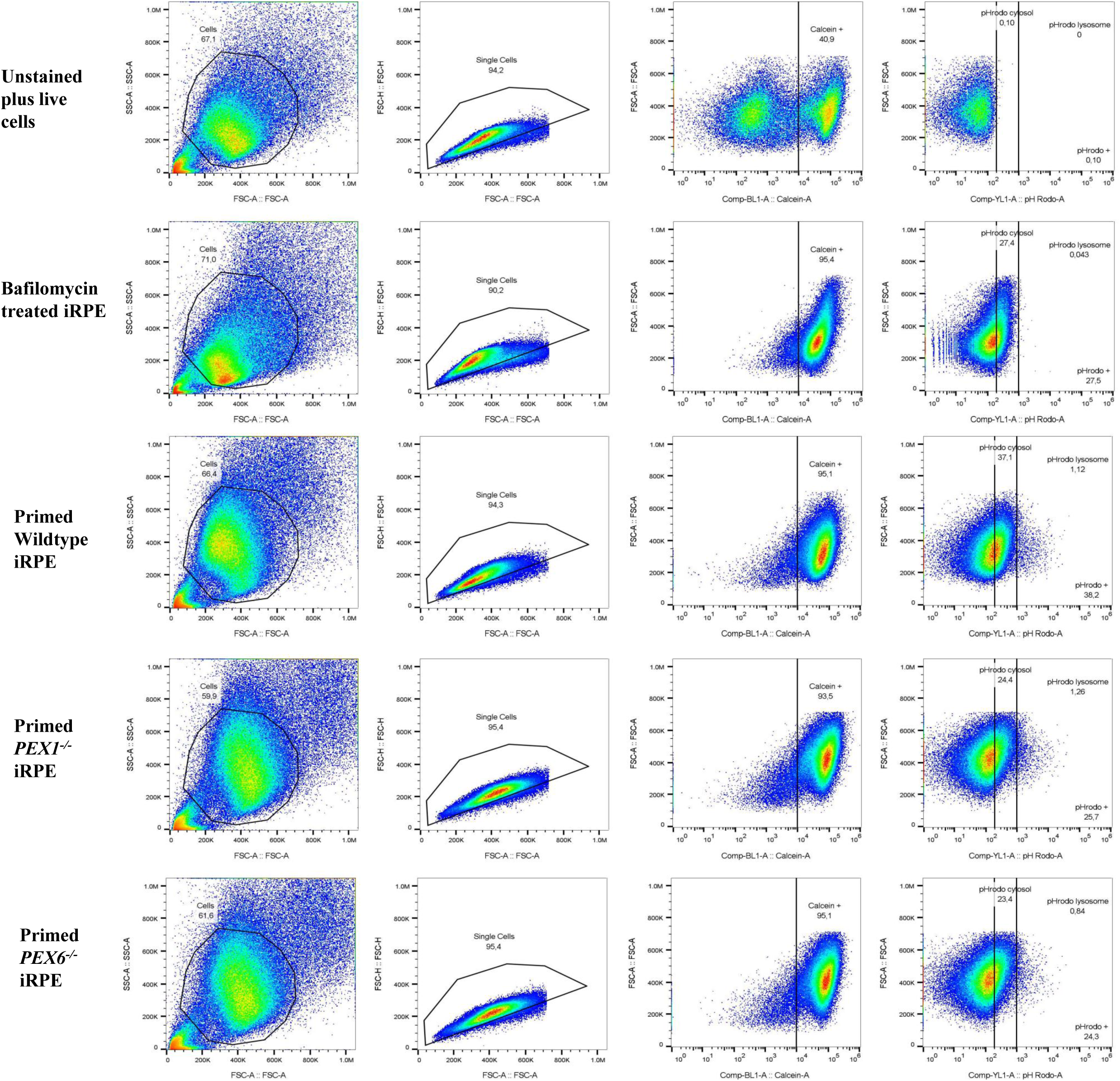
Representative flow cytometry data showing gating strategies.

**Supplementary Figure 6:**
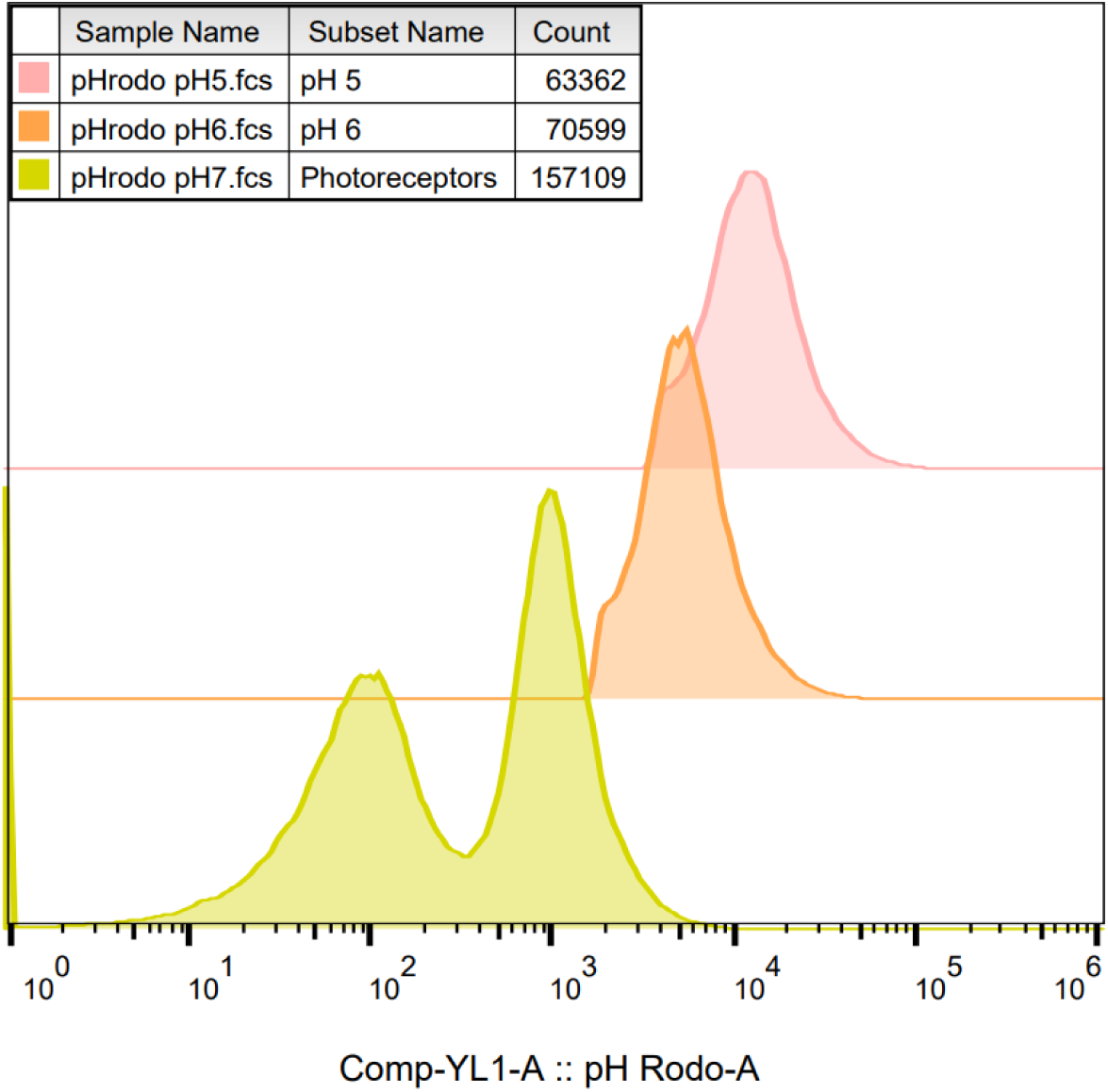
Analysis of pHrodo-labeled POS fluorescence at pH 7, pH 6, and pH 5. The pH 7 population included both pHrodo-labeled POS and unlabeled POS.

## Notes

### Competing Interest Statement

The authors have declared no competing interest.

### Summary of Updates

Figure 7 was updated to include additional data: panels D, E, F, G, and H.

