## Supplementary Tables and Figures for "Peroxisome dysfunction alters metabolism of photoreceptor outer segments in human retinal pigment epithelium"

**Supplementary Table 1. List of antibodies with corresponding application and concentration.**

FC = flow cytometry; WB = western blot; IF = immunofluorescence.

| Antibody | Clonality and Source | Manufacturer and Catalogue Number | Application and Dilution |
|---|---|---|---|
| TYRP1 | Mouse monoclonal | Novus Biologicals (32906) | FC: 1:100 (0.2 mg/mL) |
| PAX6 | Rabbit polyclonal | Proteintech (12323) | FC: 1:250 (900 µg/mL) |
| PMEL17 | Mouse monoclonal | Santa Cruz (377325) | FC: 1:100 (200 µg/mL) |
| BEST1 | Mouse monoclonal | Santa Cruz (32792) | FC: 1:100 (200 µg/mL) |
| MITF | Rabbit monoclonal | Invitrogen (32554) | FC: 1:100 (1 mg/mL) |
| Goat anti Rabbit conjugated to Alexafluor 488 | Goat polyclonal | Invitrogen (A11034) | FC: 1:500 (2 mg/mL) |
| Donkey anti mouse conjugated to Alexafluor 647 | Donkey polyclonal | Thermo Fisher (31571) | FC: 1:500 (2 mg/mL) |
| PMP70 | Rabbit polyclonal | Abcam (ab3421) | WB: 1:1000 (1 mg/mL)<br>IF: 1:50 |
| PEX1 | Rabbit polyclonal | Proteintech (13669) | WB: 1:500 (500 µg/mL) |
| PEX6 | Mouse monoclonal | Santa Cruz (sc271813) | WB: 1:50 (200 µg/mL) |
| ACAA1 | Rabbit polyclonal | Abcam (ab154091) | WB: 1:1000 (1 mg/mL) |
| ACOX1 | Mouse monoclonal | Santa Cruz (sc517306) | WB: 1:100 (100 µg/mL) |
| γ-Tubulin | Mouse monoclonal | Millipore Sigma (T5326) | WB: 1:1000 (1 mg/mL) |
| Goat anti mouse conjugated to IRDye 800 | Goat polyclonal | Li-Cor (926-32210) | WB: 1:5000 (1 mg/mL) |
| Goat anti rabbit conjugated to IRDye 800 | Goat polyclonal | Li-Cor (926-32211) | WB: 1:5000 (1 mg/mL) |
| Goat anti mouse conjugated to IRDye 680 | Goat polyclonal | Li-Cor (926-68070) | WB: 1:5000 (1 mg/mL) |

| | | | |
|---|---|---|---|
| Goat anti rabbit conjugated to IRDye 680 | Goat polyclonal | Li-Cor (926-68071) | WB: 1:5000 (1 mg/mL) |
| CLDN19 (Claudin-19) | Mouse monoclonal | Santa Cruz (365967) | IF: 1:10 (200 µg/mL) |
| ADRP (PLIN2) | Mouse monoclonal | Santa Cruz (377429) | IF: 1:100 (200 µg/mL) |
| ZO-1 | Mouse monoclonal | Thermo Fisher (339188) | IF: 1:10 (500 µg/mL) |
| RHO | Mouse monoclonal | Novus Biologicals (25160) | WB: 1:5000 (1 mg/mL) |
| β-actin | Mouse monoclonal | Santa Cruz (69879) | WB: 1:500 (100 µg/mL) |
| β-actin | Rabbit polyclonal | Abcam (ab8227) | WB: 1:1000 (300 µg/mL) |
| MFP2 | Rabbit polyclonal | Proteintech (15116-1-AP) | WB: 1:1000 (500 µg/mL) |
| PLIN2 | Rabbit polyclonal | Proteintech (15294-1-AP) | WB: 1:1000 (850 µg/mL) |
| Cathepsin D | Rabbit monoclonal | Abcam (ab75852) | WB: 1:3000 (0.184mg/mL) |
| Vinculin | Rabbit monoclonal | Thermo Fisher (MA542795) | WB: 1:20000 (0.5mg/mL) |
| Vinculin | Mouse monoclonal | Invitrogen (14-9777-82) | WB: 1:2000 (0.5mg/mL) |

**Supplementary Table 2. Comprehensive list of total fatty acid species detected in bovine POS using GC-MS.**

The mean value for each fatty acid species is presented (n=3).

| Fatty Acid | µg lipid/mg total protein | Percentage of Total Lipids Detected |
|---|---|---|
| C10:0 - Capric | 0.002 | 0.00% |
| C12:0 - Lauric | 0.007 | 0.00% |
| C14:0 - Myristic | 0.271 | 0.20% |
| C15:0 - Pentadecanoic | 0.167 | 0.12% |
| C16:0 - Palmitic | 19.345 | 14.20% |
| C17:0 - Heptadecanoic | 0.576 | 0.42% |
| C18:0- Stearic | 28.393 | 20.84% |
| C20:0 - Arachidic | 0.127 | 0.09% |
| C22:0 - Behenic | 0.067 | 0.05% |
| C23:0 - Tricosanoic | 0.006 | 0.00% |
| C24:0 - Lignoceric | 0.077 | 0.06% |
| C25:0 - Pentacosanoic | 0.002 | 0.00% |
| C26:0 - Hexacosanoic | 0.009 | 0.01% |
| C28:0 - Octacosanoic | 0.002 | 0.00% |
| C30:0 | 0.001 | 0.00% |
| C10:1 - Caproleic | 0.001 | 0.00% |

| | | |
|---|---|---|
| C12:1 - Dodecaenoic | 0.001 | 0.00% |
| C16:1ω9 | 0.205 | 0.15% |
| C17:1 - Heptadecaenoic | 0.055 | 0.04% |
| C18: 1ω9 - Oleic | 3.001 | 2.20% |
| C20: 1ω9 - Eicosenoic | 0.079 | 0.06% |
| C20:3ω9- Mead | 0.041 | 0.03% |
| C22:1ω9 - Erucic | 0.023 | 0.02% |
| C24:1ω9 - Nervonic | 0.021 | 0.02% |
| C26:1ω9 | 0.002 | 0.00% |
| C14:1 - Myristoleic | 0.001 | 0.00% |
| C16:1ω7 - Palmitoleic | 0.069 | 0.05% |
| C18:1ω7 - Vaccenic | 0.977 | 0.72% |
| C18:1ω5 | 0.011 | 0.01% |
| C20:3ω7 | 0.021 | 0.02% |
| C14:2 - Myristolenic | 0.010 | 0.01% |
| C16:2 - Palmitolenic | 0.027 | 0.02% |
| C18:2ω6 - Linoleic | 1.672 | 1.23% |
| C18:2ω6 Conj - Rumenic | 0.011 | 0.01% |
| C18:3ω6 - Gamma Linolenic | 0.463 | 0.34% |
| C20:2ω6 - Eicosadienoic | 0.266 | 0.20% |
| C20:3ω6 - Dihomo-g-linolenic | 0.921 | 0.68% |
| C20:4ω6 - Arachidonic | 8.011 | 5.88% |
| C22:2ω6 - Docosadienoic | 0.019 | 0.01% |
| C22:4ω6 - Adrenic | 6.024 | 4.42% |
| C22:5ω6 - Docosapentaenoic | 13.722 | 10.07% |
| C24:2ω6 | 0.011 | 0.01% |
| C26:2 - Hexacosadienoic | 0.002 | 0.00% |
| C18:3ω3 - Alpha Linolenic | 0.014 | 0.01% |
| C20:5ω3 - Eicosapentaenoic | 0.030 | 0.02% |
| C22:5ω3 | 1.263 | 0.93% |
| C22:6ω3 - Docosahexanoic | 49.867 | 36.61% |
| Pristanic acid | 0.001 | 0.00% |
| Phytanic acid | 0.013 | 0.01% |
| C16:1 trans | 0.071 | 0.05% |
| C18:1 trans | 0.190 | 0.14% |
| C18:2 trans | 0.048 | 0.04% |

DNA Sanger Sequencing

DNA was isolated from iRPE using GeneJET Genomic DNA Purification Kit (Thermo Scientific, Cat. No. K0722) and eluted in molecular grade water. DNA concentration was determined using a µDrop™ plate (Thermo Scientific, Cat. No. N12391) and Multiskan™ GO Microplate Spectrophotometer (Thermo Scientific, Cat. No. 51119300).

To confirm the genotype of *PEX1*$^{-/-}$ iRPE, 50 µL PCR reactions were performed using Taq DNA polymerase (New England Biolabs, Cat. No. M0273L) with forward (5'-GAACTCTTTTGGACATGTGAATTG-3') and reverse (5'-GCAAGTAGGGAGTATGGTAAACT-3') primers (0.2 µM). Reactions were subject to one denaturation step for 30 seconds at 95 °C, followed by 30 cycles of 15 seconds duration at 95 °C, 15 seconds at 49 °C, and 27 seconds at 68 °C, followed by one extension step for 5 minutes at 68 °C.

To confirm the genotype of *PEX6*$^{-/-}$iRPE, 50 µL touchdown PCR reactions were performed using Phusion DNA polymerase and GC buffer (Thermo Scientific, Cat. No. F534S) with forward (5'-AGAAACCGCAAAGGAGGAC-3') and reverse (5'-ACTAGTCGTCTGGCTCTCTG-3') primers (0.2 µM). Reactions were subject to one denaturation step for 30 seconds at 98 °C, followed by 10 touchdown cycles of 10 seconds at 98 °C, 20 seconds at 66 °C (decreasing by 0.5 °C each cycle), and 21 seconds at 72 °C. This was followed by 25 cycles of 30 seconds at 98 °C, 20 seconds at 61 °C, and 21 seconds at 72 °C, followed by one extension step for 5 minutes at 72 °C.

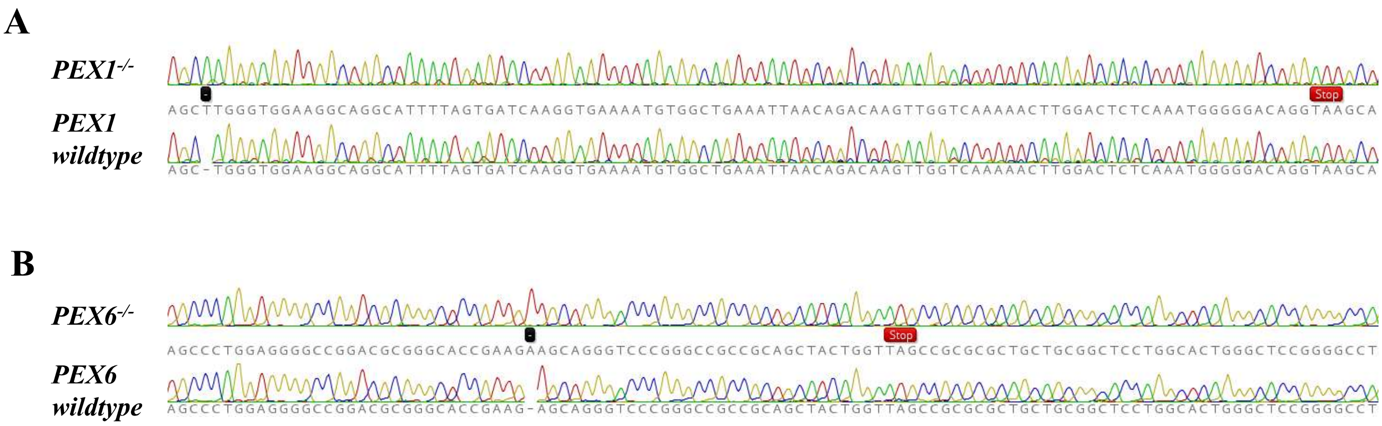


**Supplementary Figure 1: Sanger sequencing of iRPE DNA. Frameshift mutations caused by single base pair duplications (indicated by a black square) resulted in stop codons (indicated by red rectangles).**

*A.* *PEX1*$^{-/-}$ iRPE DNA compared to wildtype.
*B.* *PEX6*$^{-/-}$ iRPE DNA compared to wildtype.

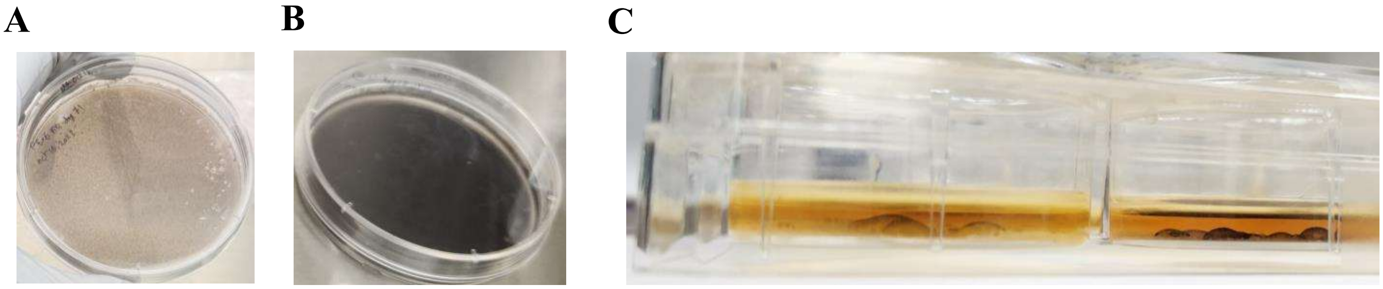


**Supplementary Figure 2: Pigmentation and focal monolayer detachments in iRPE.**

A. Maturing iRPE cells at day 74 demonstrate mild pigmentation.
B. Mature iRPE at day 95 demonstrate significant pigmentation.
C. Cross-sectional image of differentiating iRPE cells demonstrate focal monolayer detachments (day 37).

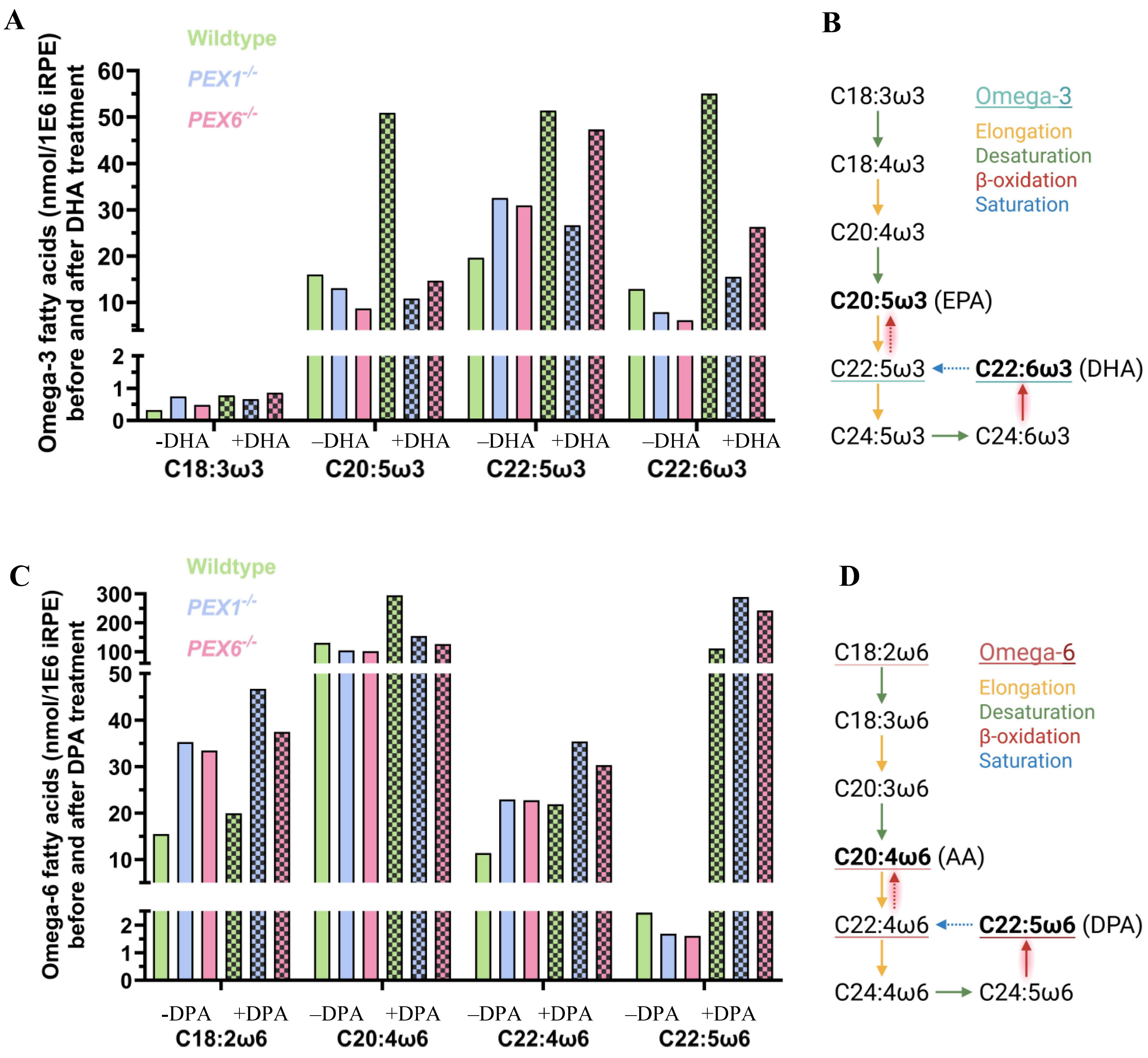


**Supplementary Figure 3: Quantitative analysis of fatty acid profiles shifts in iRPE following DHA and DPA treatments.**

A. Quantification of omega-3 fatty acids in iRPE lysates by GC-MS before and after a 24-hour treatment with 30 µM DHA (n=1).
B. The omega-3 fatty acid metabolic pathway adapted from Yu et al., 2012.[76] Reactions that are presumed to require intact peroxisome β-oxidation are highlighted in red.
C. Quantification of omega-6 fatty acids in iRPE lysates by GC-MS before and after a 24-hour treatment with 30 µM omega-6 DPA (n=1).
D. The omega-6 fatty acid metabolic pathway adapted from Yu et al., 2012.[76] Reactions that are presumed to require intact peroxisome β-oxidation are highlighted in red.

### Relative Quantification Plot

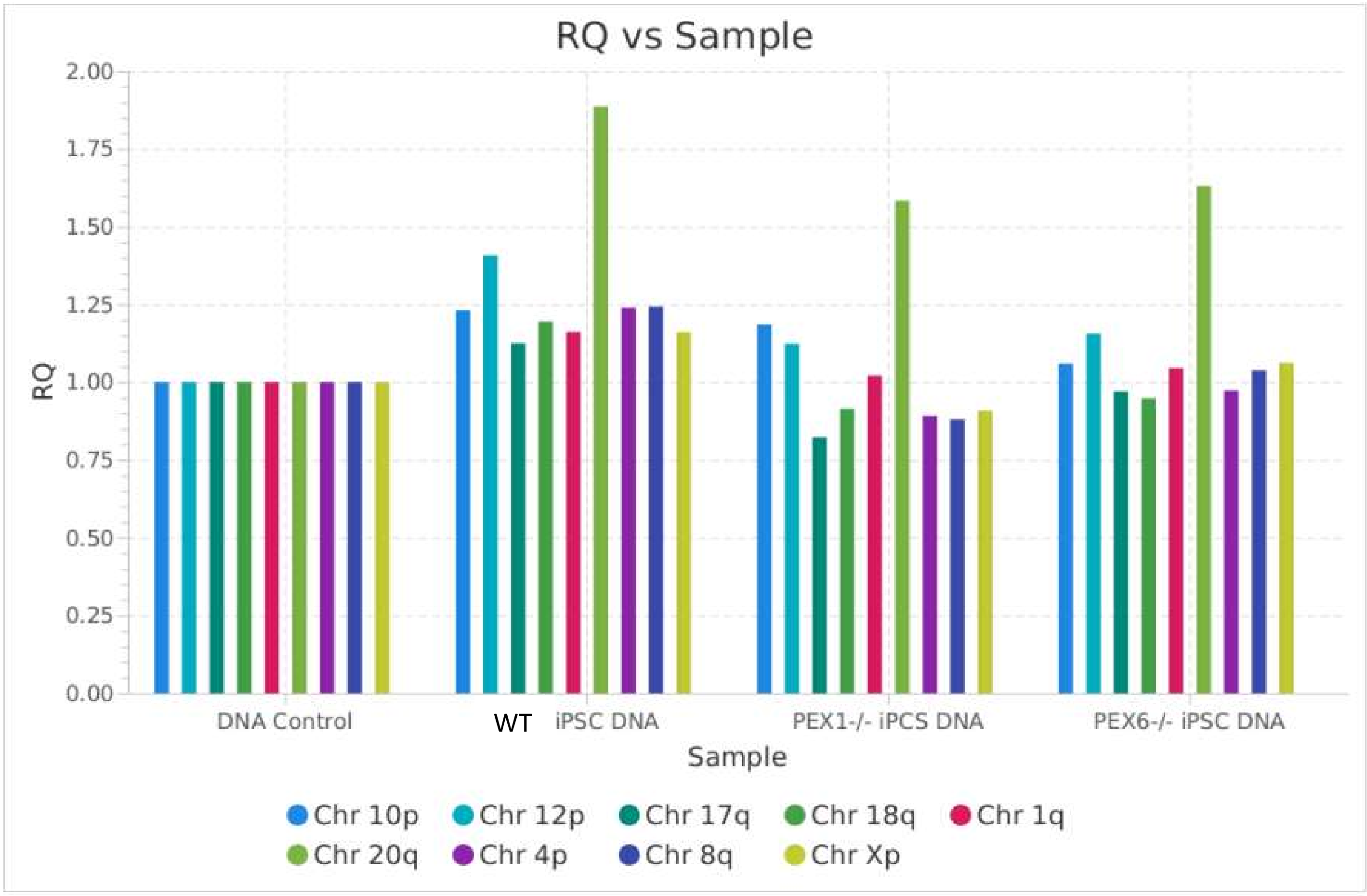


**Supplementary Figure 4: Quantitative PCR in wildtype, *PEX1*$^{-/-}$ and *PEX6*$^{-/-}$ iPSCs for commonly reported karyotypic abnormalities.** Chr20q has an amplification in a minimal critical region in wildtype, *PEX1*$^{-/-}$ and *PEX6*$^{-/-}$ iPSCs.

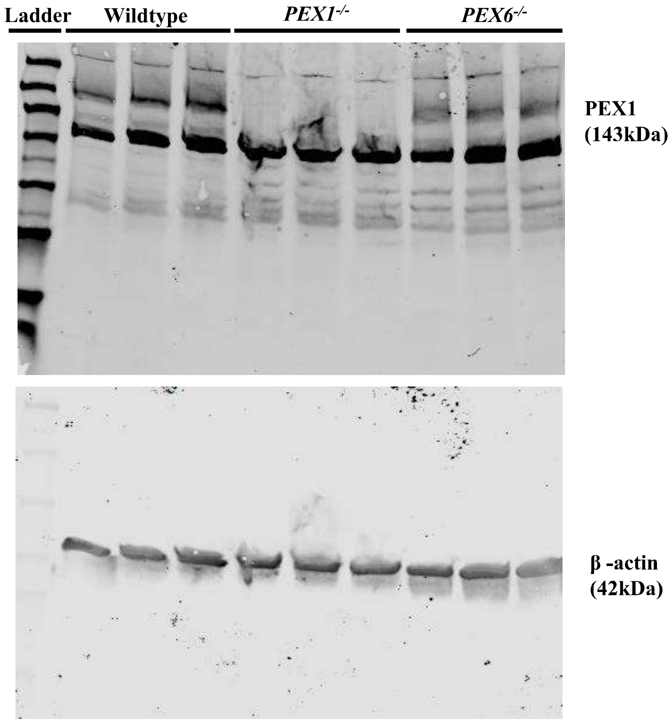
Ladder
Wildtype
PEX1-/-
PEX6-/-
PEX1
(143kDa)
β -actin
(42kDa)

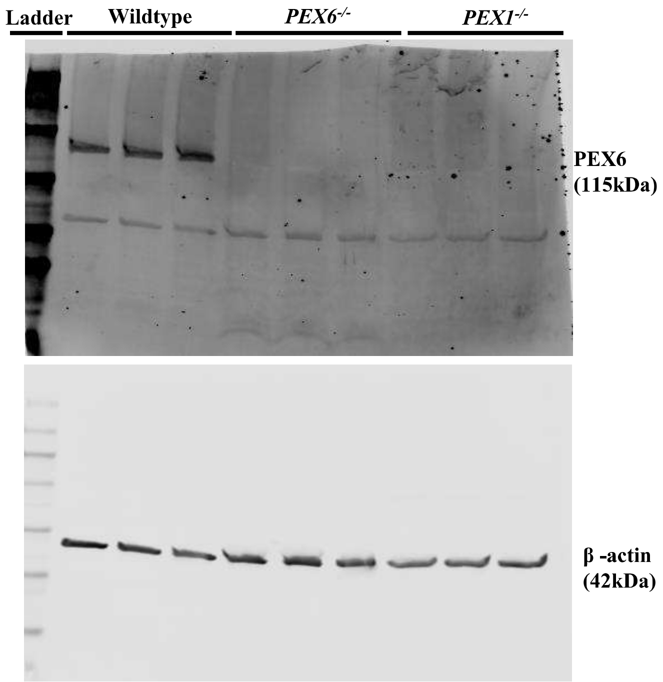
Ladder
Wildtype
PEX6-/-
PEX1-/-
PEX6
(115kDa)
β -actin
(42kDa)

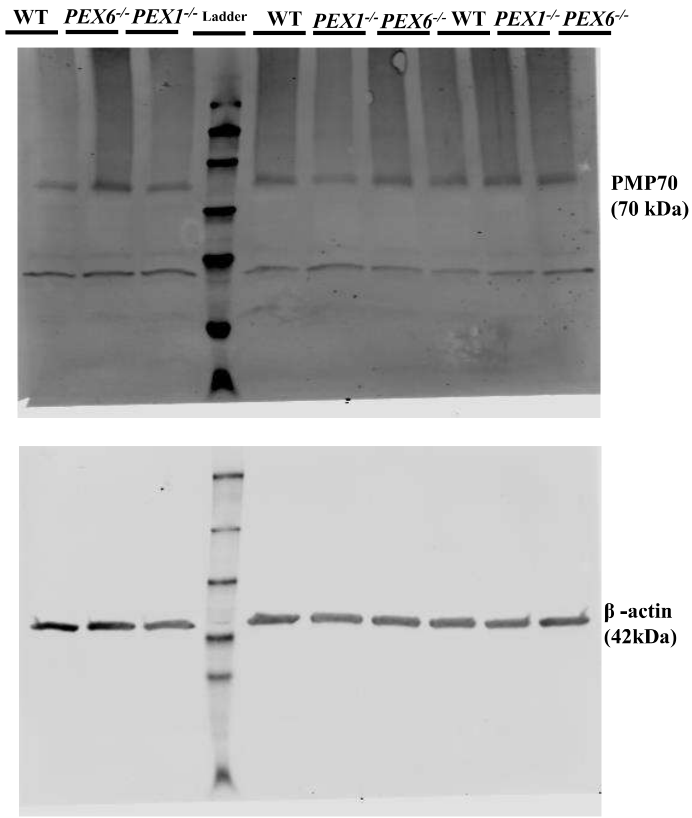
WT
PEX6-/-
PEX1-/-
Ladder
WT
PEX1-/-
PEX6-/-
WT
PEX1-/-
PEX6-/-
PMP70
(70 kDa)
β -actin
(42kDa)

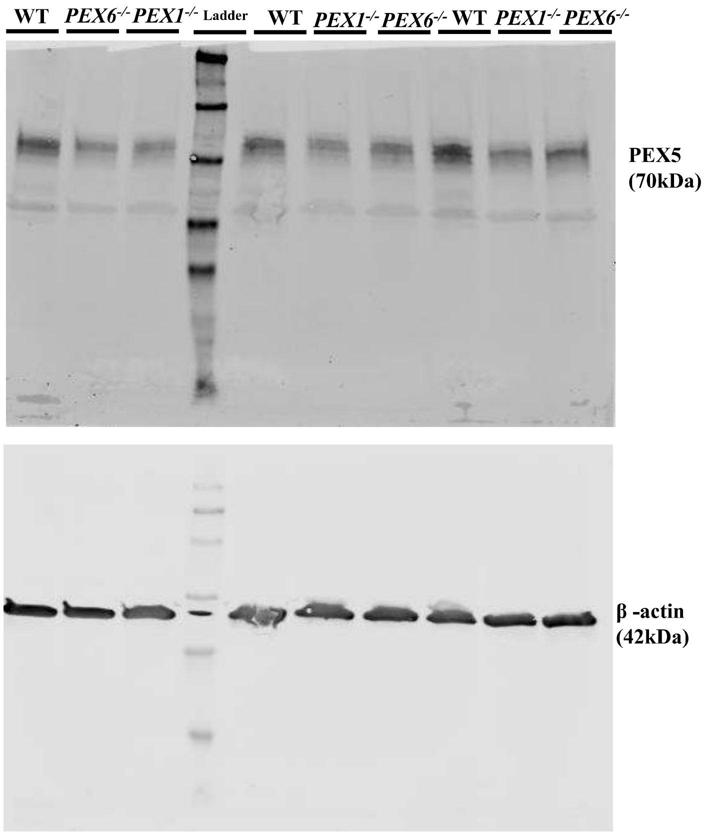
WT
PEX6-/-
PEX1-/-
Ladder
WT
PEX1-/-
PEX6-/-
WT
PEX1-/-
PEX6-/-
PEX5
(70kDa)
β -actin
(42kDa)

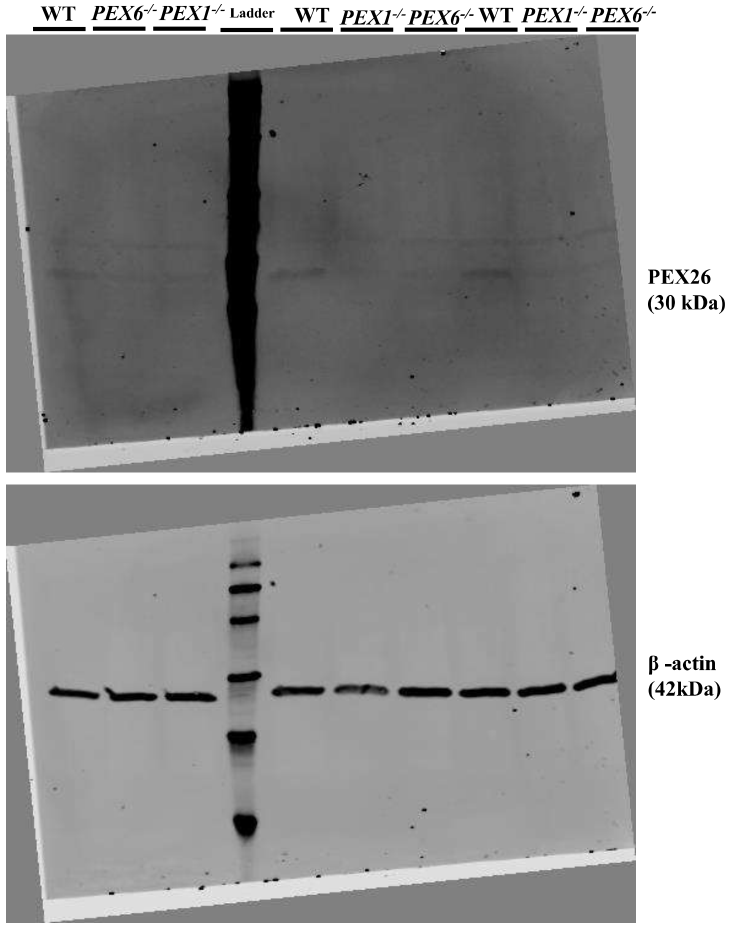
WT PEX6-/- PEX1-/- Ladder WT PEX1-/- PEX6-/- WT PEX1-/- PEX6-/-
PEX26 (30 kDa)
β -actin (42kDa)

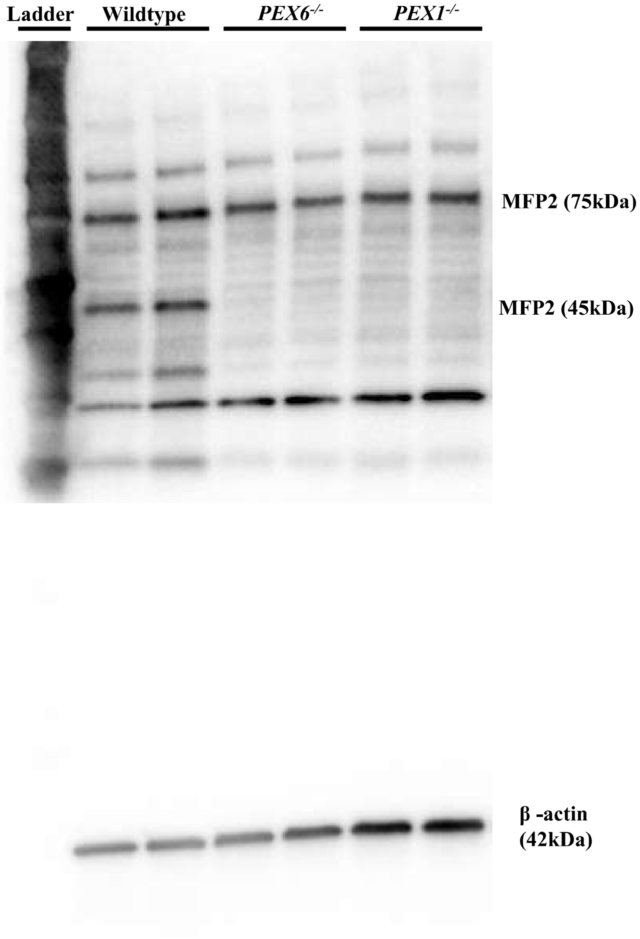
Ladder Wildtype PEX6-/- PEX1-/-
MFP2 (75kDa)
MFP2 (45kDa)
β -actin (42kDa)

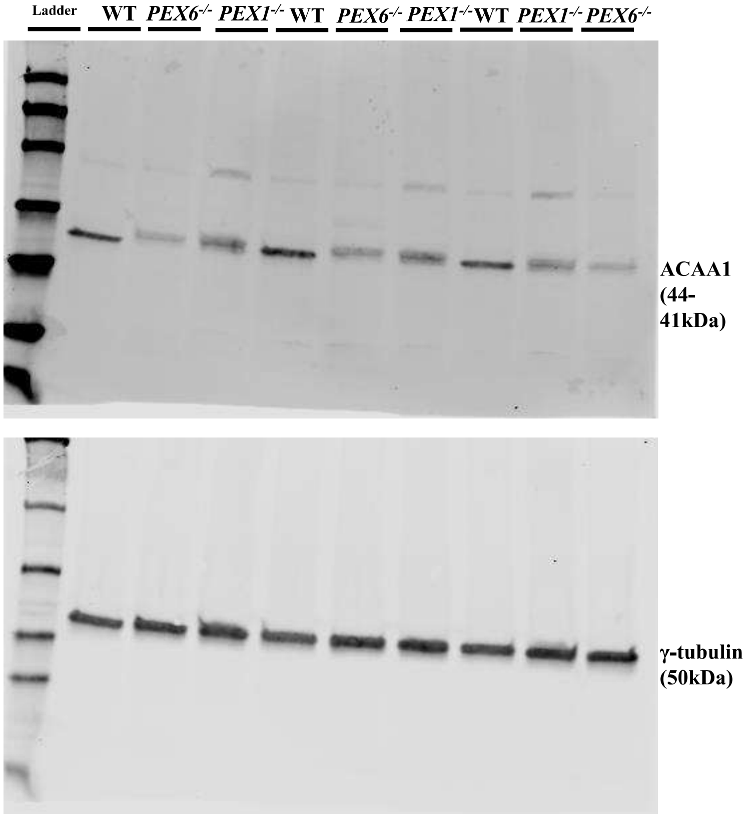
Ladder WT PEX6-/- PEX1-/- WT PEX6-/- PEX1-/- WT PEX1-/- PEX6-/-
ACAA1 (44-41kDa)
γ-tubulin (50kDa)

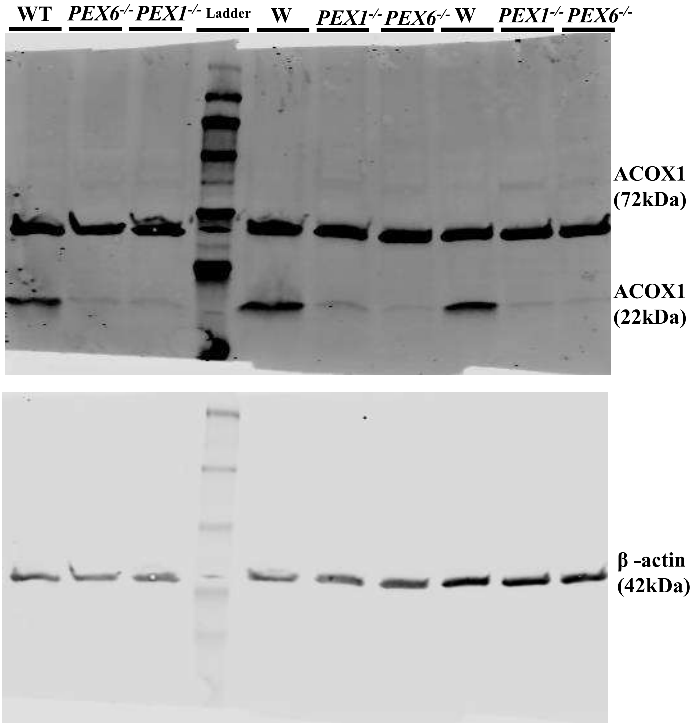
WT PEX6-/- PEX1-/- Ladder W PEX1-/- PEX6-/- W PEX1-/- PEX6-/-
ACOX1 (72kDa)
ACOX1 (22kDa)
β -actin (42kDa)

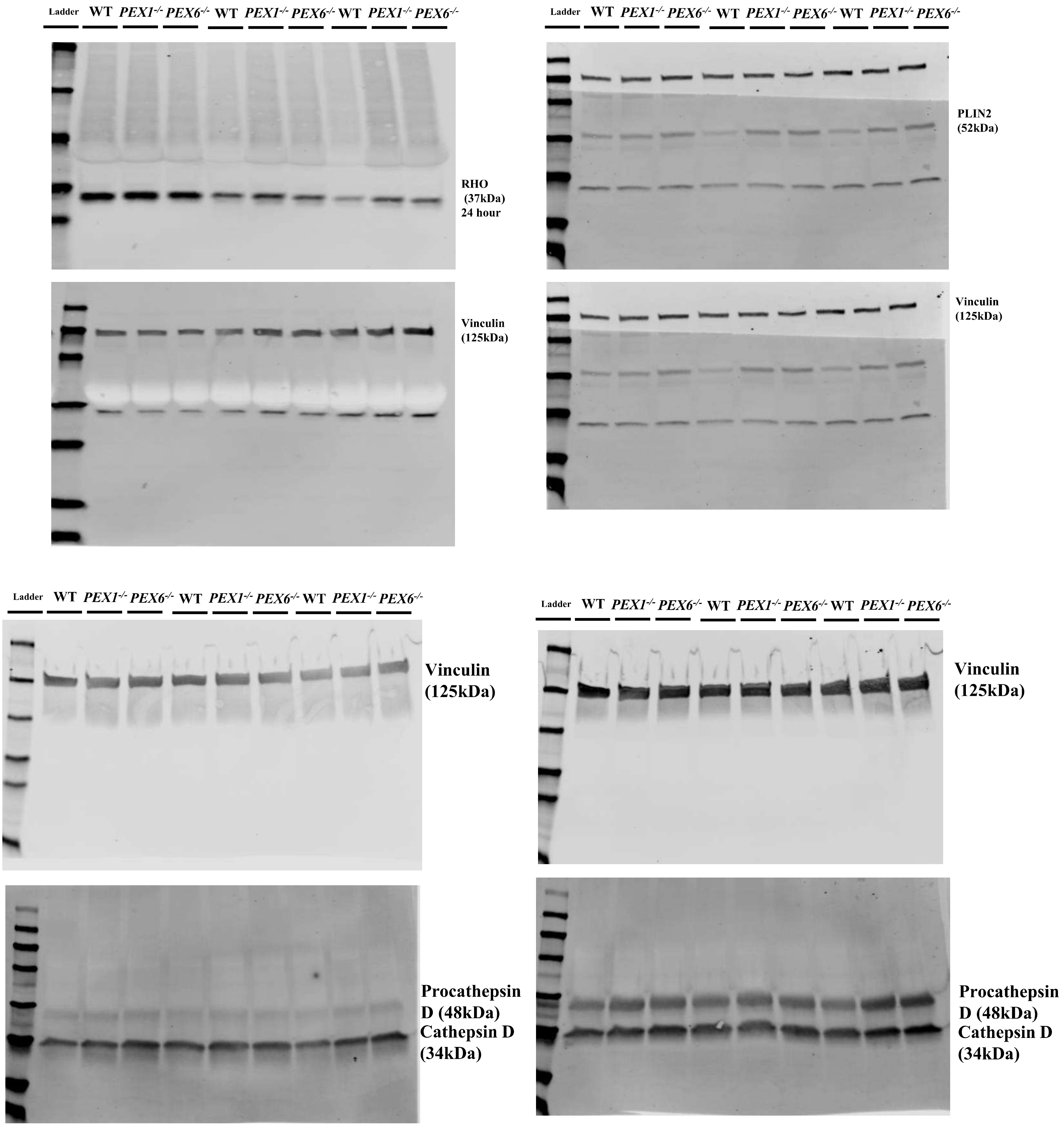


**Supplementary Figure 4: Full membranes of all immunoblot images presented.**

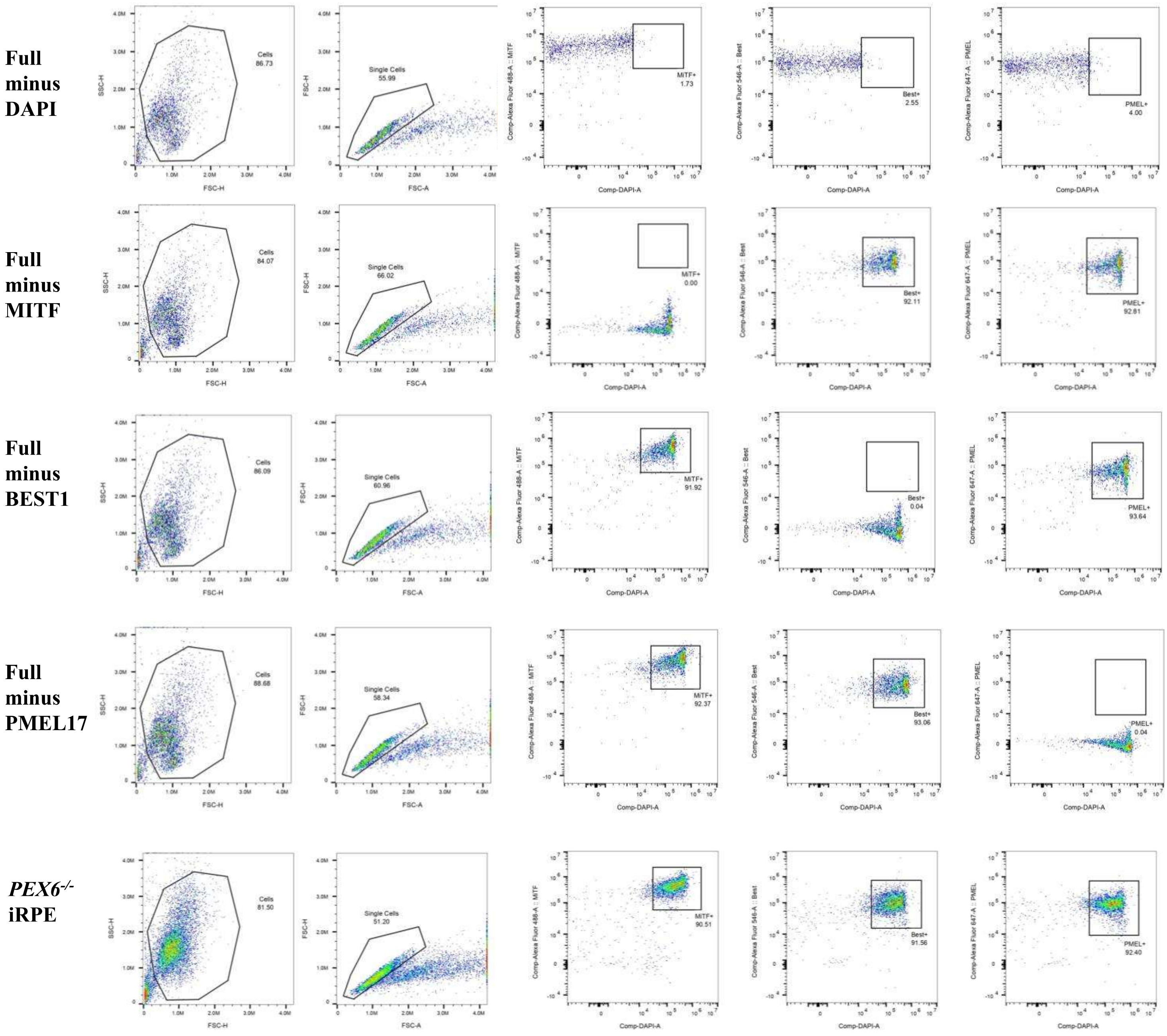
Full minus DAPI
Full minus MITF
Full minus BEST1
Full minus PMEL17
PEX6-/- iRPE
Cells 86.73
Single Cells 55.99
MITF+ 1.73
Best+ 2.55
PMEL+ 4.00
Cells 84.07
Single Cells 66.02
MITF+ 0.00
Best+ 92.11
PMEL+ 92.81
Cells 86.09
Single Cells 60.96
MITF+ 91.92
Best+ 0.04
PMEL+ 93.64
Cells 88.68
Single Cells 58.34
MITF+ 92.37
Best+ 93.06
PMEL+ 0.04
Cells 81.50
Single Cells 51.20
MITF+ 90.51
Best+ 91.56
PMEL+ 92.40
FSC-H
SSC-H
FSC-A
Comp-DAPI-A
Comp-Alexa Fluor 488-A :: MITF
Comp-Alexa Fluor 546-A :: Best
Comp-Alexa Fluor 647-A :: PMEL

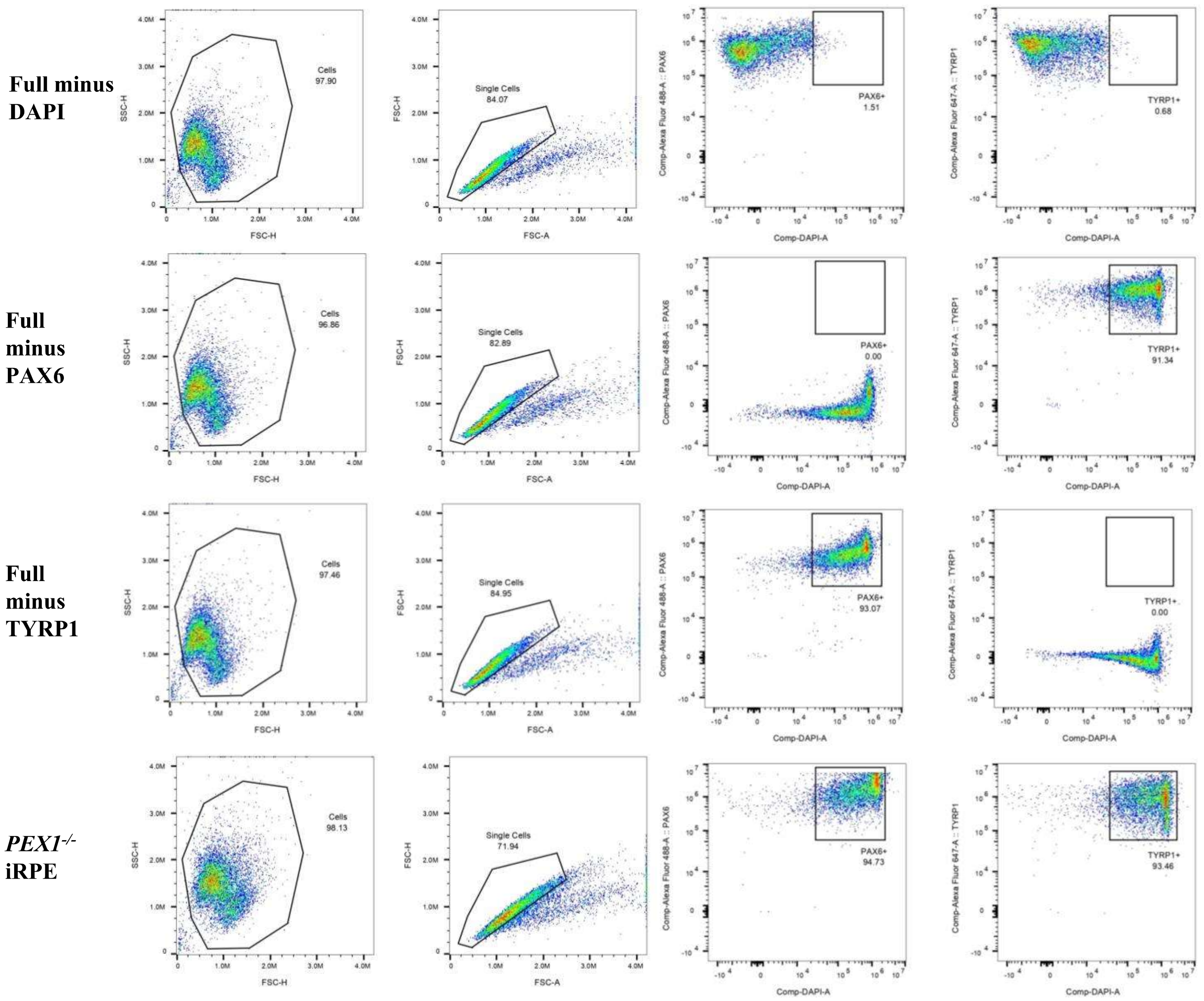
Full minus DAPI
Full minus PAX6
Full minus TYRP1
*PEX1*$^{-/-}$ iRPE
Cells
97.90
Single Cells
84.07
PAX6+
1.51
TYRP1+
0.68
Cells
96.86
Single Cells
82.89
PAX6+
0.00
TYRP1+
91.34
Cells
97.46
Single Cells
84.95
PAX6+
93.07
TYRP1+
0.00
Cells
98.13
Single Cells
71.94
PAX6+
94.73
TYRP1+
93.46
SSC-H
FSC-H
FSC-A
Comp-Alexa Fluor 488-A :: PAX6
Comp-Alexa Fluor 647-A :: TYRP1
Comp-DAPI-A

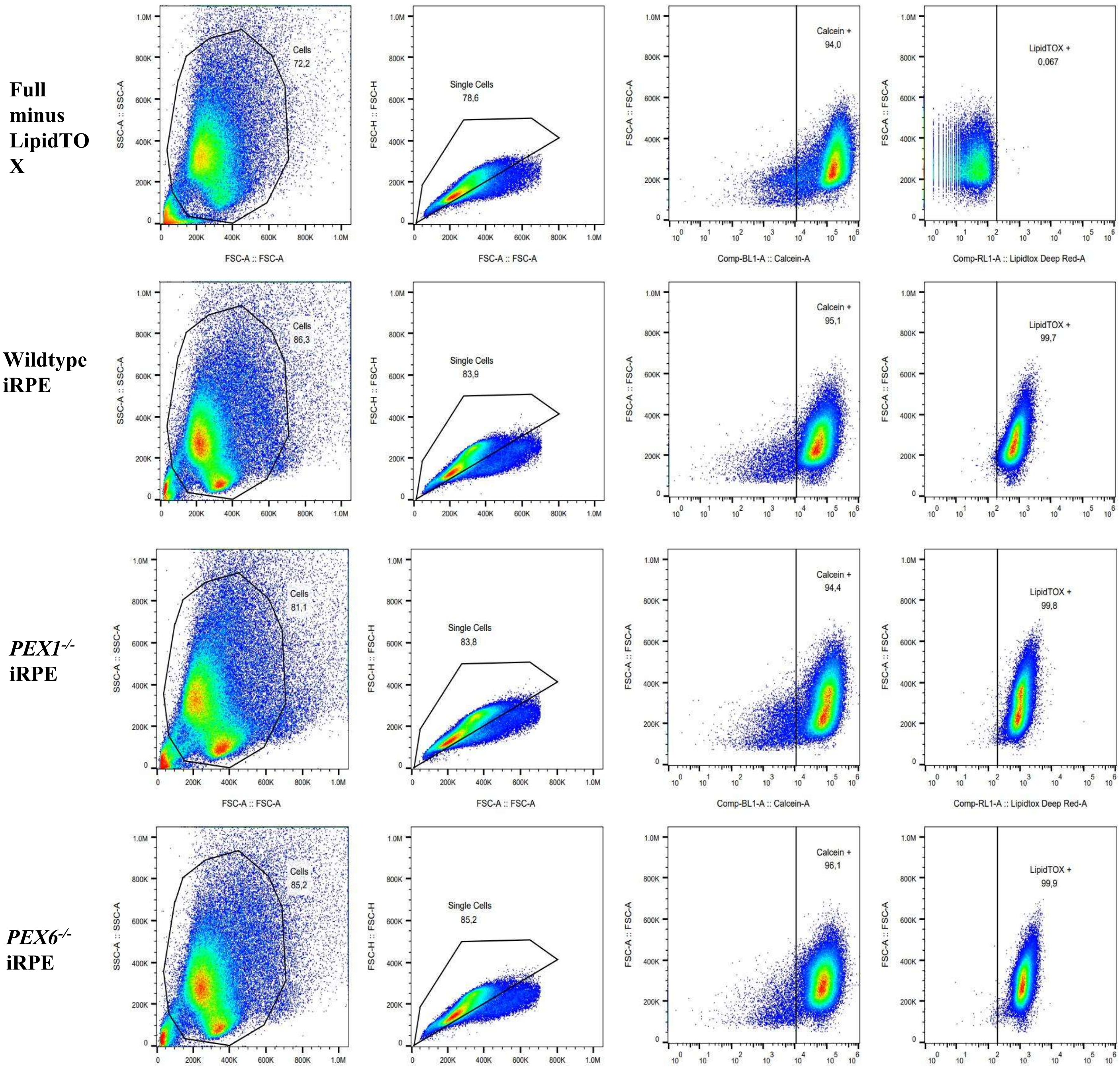
Full minus LipidTOX
Wildtype iRPE
PEX1-/- iRPE
PEX6-/- iRPE
Cells 72,2
Single Cells 78,6
Calcein + 94,0
LipidTOX + 0,067
Cells 86,3
Single Cells 83,9
Calcein + 95,1
LipidTOX + 99,7
Cells 81,1
Single Cells 83,8
Calcein + 94,4
LipidTOX + 99,8
Cells 85,2
Single Cells 85,2
Calcein + 96,1
LipidTOX + 99,9
SSC-A :: SSC-A
FSC-A :: FSC-A
FSC-H :: FSC-H
Comp-BL1-A :: Calcein-A
Comp-RL1-A :: Lipidtox Deep Red-A

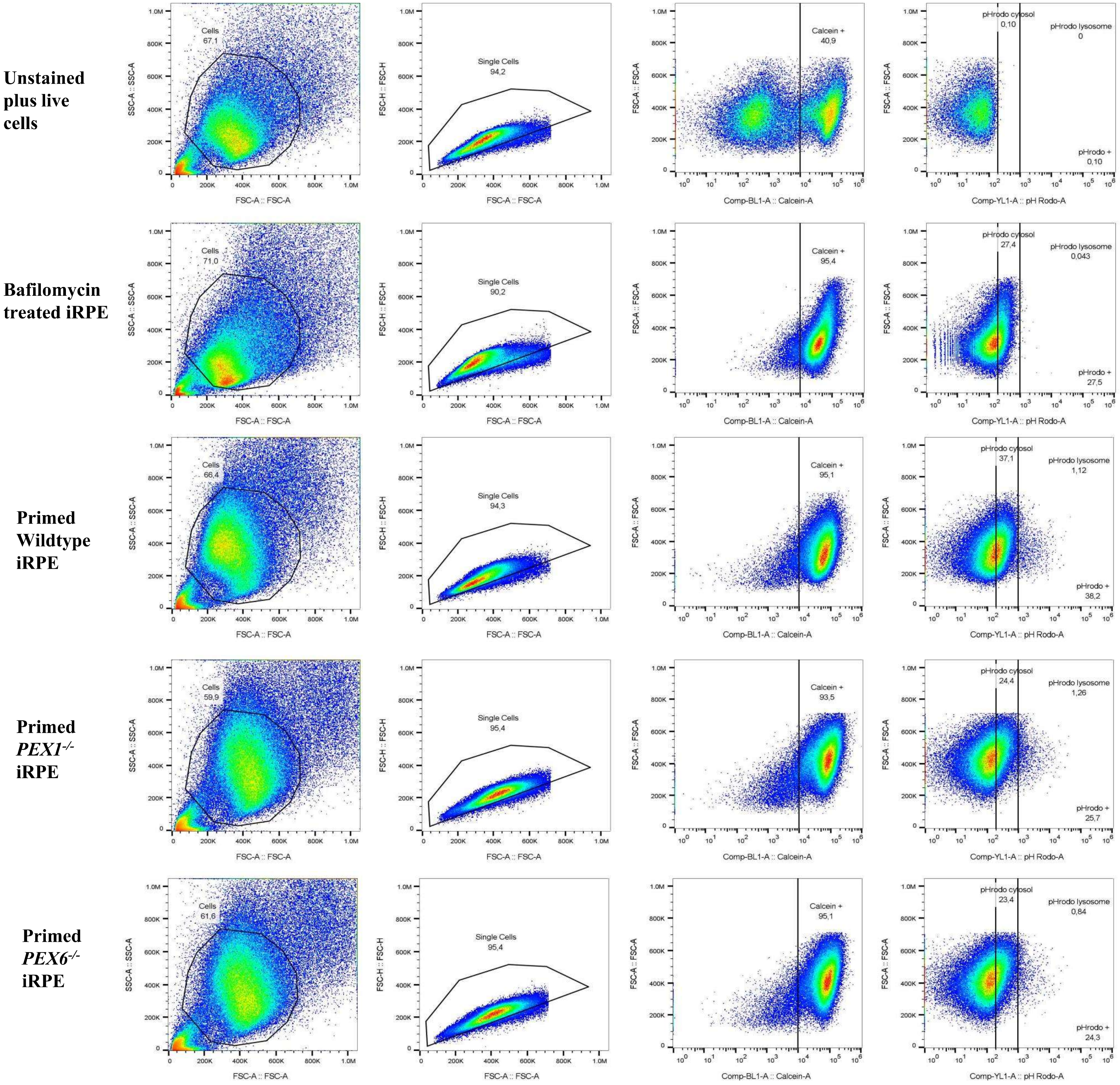


**Supplementary Figure 5: Representative flow cytometry data showing gating strategies.**

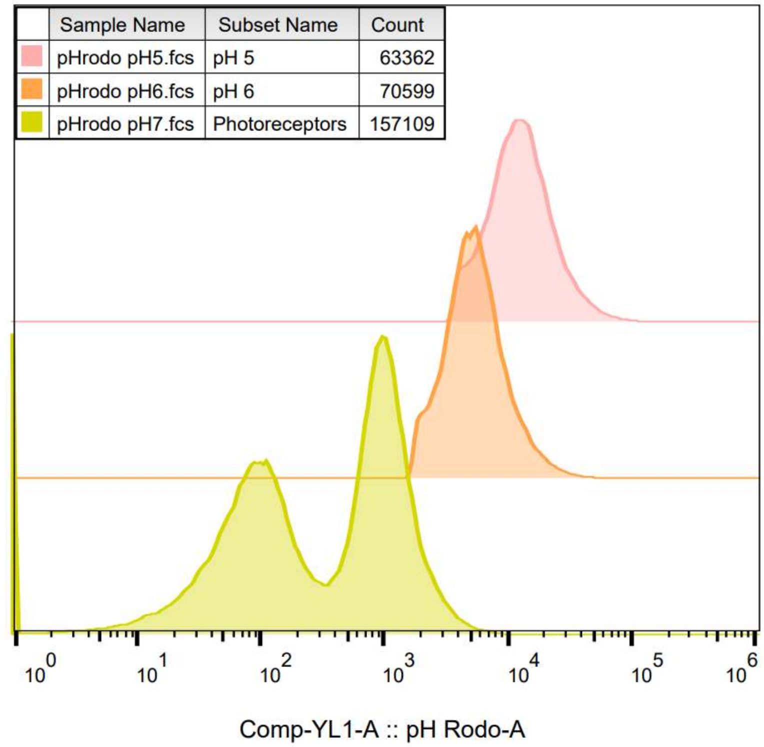


**Supplementary Figure 6: Analysis of pHrodo-labeled POS fluorescence at pH 7, pH 6, and pH 5. The pH 7 population included both pHrodo-labeled POS and unlabeled POS.**
